# Non-consensus flanking sequence of hundreds of base pairs around in vivo binding sites: statistical beacons for transcription factor scanning

**DOI:** 10.1101/2025.05.28.656598

**Authors:** Kateřina Faltejsková, Josef Šulc, Jiří Vondrášek

## Abstract

It was long suspected that for specific DNA binding by a transcription factor, the flanks of the binding motifs can play an important role. By a thorough analysis of the DNA sequence in the broad context (± 5000 bp) of in vivo binding sites (as identified in a ChIP-seq or a Cut&Tag experiment), we show that the average GC content is in most cases statistically significantly increased around the binding site in a patch spanning 1000–1500 bp. This increase was observed consistently in experiment targeting the same TF in different cell lines. The surrounding of binding sites of certain TFs like MYC display a directional alteration of dinucleotide frequencies. We attempt to explain these preferences by alteration in DNA shape features as well as by potential cooperation with other TFs. We observed differences in sequence affinity to various potential cooperating TFs between cell lines. Altogether, we propose that the observed feature distortion is indicative of a coarse scanning mechanism that helps TFs find the target binding site.

**Graphical Abstract:** 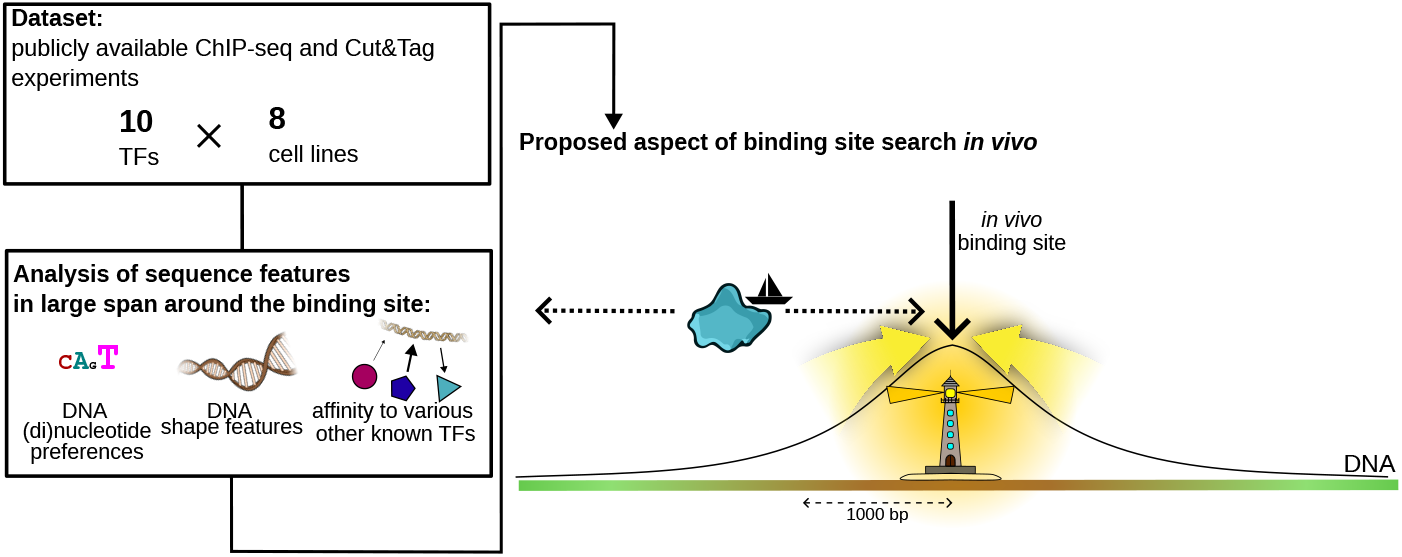

**Key Messages:**

- We prove an increase in GC content 1000–1500 bp both upstream and downstream of the binding site.
- We observed a funnel-like sequence signature of thousands of bp that is shared between most cell lines.
- We propose an explanation based on helix shape and binding affinities of potential cooperating TFs, supported by crosslinking the ChIP-seq (Cut&Tag) experiments with ATAC-seq experiments.

## Introduction

Transcription factors (TFs) facilitate the transmission of information from DNA to downstream molecular processes. These molecules specifically bind motifs of mostly 6-12 bp [Kim and Wysocka, 2023, Slattery et al., 2014]. Such a length (averaging ~10 bp) balances binding specificity and the ability of TFs to regulate multiple genomic loci efficiently, although longer motifs have been observed [Stewart et al., 2012].

It has been known for some time that only a fraction of motifs capable of being bound are truly bound in vivo at a given point in time [Kim and Wysocka, 2023]. There appear to be several reasons for this selectivity. Firstly, many TFs do not function as monomers. The binding motifs of monomeric TFs and TFs in dimers can differ significantly [Jolma et al., 2013, Xie et al., 2025]. These composite motifs are important for acquiring specificity during cell differentiation [Xie et al., 2025]. Additionally, the affinity and biological role of TFs can change depending on cooperating molecules, highlighting a flexible functionality [Martinez-Corral et al., 2024, Mahendrawada et al., 2025]. For instance, yeast contains more TFs with a wide activity range than those with uniquely defined functions [Mahendrawada et al., 2025].

When describing TF binding, increasing attention is being paid to the structure of the binding site. Describing the shape of the binding site can be more informative for binding predictions than the motif sequence alone [Mathelier et al., 2016, Samee et al., 2019, Chen et al., 2024]. The structure and flexibility of DNA flanking the motif also significantly influence the overall affinity of a TF for its binding site [Ghoshdastidar and Bansal, 2022, Yella et al., 2018, Chiu et al., 2022]. Recent studies emphasized flexibility of DNA flanking regions as crucial determinants of TF binding specificity [Ghoshdastidar and Bansal, 2022].

By gradually shifting from a detailed to a broad perspective, we can conceptualize the recognition process as comprising several overlapping levels. At the lowest level, base readout involves direct recognition through chemical contacts, including hydrogen bonds, electrostatic interactions, water-mediated bonds, and hydrophobic contacts, all contributing directly to binding specificity and affinity. Namely, the preference for a given nucleotide at a specific position is mainly determined by interactions between the amino acid side chains of the TF and the accessible edges of the base pairs that are contacted.

Other modes of readout, especially minor groove recognition, rely on detecting variations its width and electrostatical potential of its surface [Rohs et al., 2010]. This readout mechanism is often shown as an example of shape readout. Shape readout refers to TF recognition based on structural alterations of the DNA helix (e.g., roll, twist, tilt, groove geometry) that vary with sequence context [Mathelier et al., 2016]. For example, Hox proteins promote bending through such perturbations, generating unique conformations that favor binding [Ghoshdastidar and Bansal, 2022].

Protein–DNA backbone interactions are pervasive: approximately two thirds of direct hydrogen bonds and approximately three quarters of van der Walls contacts involve the sugar–phosphate backbone, with a strong bias toward the highly exposed negatively charged phosphates. These contacts are largely sequence-independent and mainly stabilize binding (and can contribute to indirect readout via DNA shape/structure). In contrast, direct amino acid–base interactions are the main drivers of sequence-specific readout [Sathyapriya and Vishveshwara, 2004].

Incorporating shape data into previously sequence-only predictors significantly improves TF-binding site predictions [Rohs et al., 2010, Zhou et al., 2015, Wang et al., 2021, Mathelier et al., 2016]. Indeed, it has been shown on numerous occasions that integrating DNA-shape information into previously sequence-only based predictors gives consistent improvement in binding site prediction in a significant number of cases [Mathelier et al., 2016, Rohs et al., 2010, Zhou et al., 2015, Wang et al., 2021].

Several studies have also indicated that TF preferences for specific DNA shapes extend beyond core motifs, with sequences immediately flanking TF-binding sites contributing substantially to binding specificity [Gordân et al., 2013, Dror et al., 2015, Yella et al., 2018]. Clearly, incorporating the sequence and structural context surrounding TF binding sites improves computational predictions and better aligns with observed biological outcomes. Additionally, it improves the fit between our models and *in vivo* reality [Yang et al., 2017].

Apart from the precise interplay of base and shape readout, less local influences help guide TF diffusion kinetics, optimizing search efficiency for target DNA sequences. These indirect interactions optimize the search time needed to find the right target DNA element. Considering random naive 3D diffusion within the crowded nuclear environment, the likelihood of locating a 1 bp target (approximately 0.34 nm) is extremely low. Experimentally measured association rates are ~100-fold faster than the maximum allowed by simple 3D diffusion [Slutsky and Mirny, 2004].

To accelerate target-site localization, TFs utilize facilitated diffusion, combining 1D scanning (sliding, hopping, intersegmental transfer) with short-range 3D diffusion, significantly improving search efficiency [Halford and Marko, 2004, Winter et al., 1981, Slutsky and Mirny, 2004]. This is clearly seen in some larger organizational elements in the genome. Noticeably, repeating oligomeric poly(dA:dT) and poly(dC:dG) tracts seem to attract or repulse TFs to/from (non)specific binding sites, as has also been observed *in vitro* [Afek and Lukatsky, 2013a,b, Sela and Lukatsky, 2011].

Positive design elements—extended poly(dA:dT) tracts—create low-energy ‘highways’ that statistically enhance nonspecific TF–DNA affinity and funnel TFs toward nearby cognate sites [Sela and Lukatsky, 2011]. Homo-oligonucleotide sequence correlations, such as these tracts, broaden the TF–DNA binding energy distribution, thus lowering overall nonspecific binding free energy. In contrast, DNA with alternating correlations narrows the energy spectrum and raises the nonspecific TF–DNA binding free energy [Sela and Lukatsky, 2011]. These factors allow for both positive and negative non-consensus design, greatly expanding the targeting system of the DNA transcription [Afek and Lukatsky, 2013b]. Collectively, these elements shape DNA’s geometry and electrostatic landscape, forming regions that attract or repel TFs over longer distances. This finely tuned structure optimizes the speed and specificity of TF binding. Combined with base-specific and shape-specific DNA recognition, as well as facilitated diffusion, these chromatin characteristics sculpt a multiscale free-energy landscape, directing TF movement from individual base pairs to higher-order genome structures.

In this work, we attempt to thoroughly describe the sequence features of the *in vivo* binding sites for 10 various TFs and several different cell lines. Building on the work of Dror et al., who proposed that TF binding sites are surrounded by a patch with altered GC content (increased around binding sites of TFs from *e. g*. SP1 and ELF1, decreased around binding sites of homeodomain TFs), we describe the sequence preferences and derivable features within the vast surrounding region of the binding sites. We discover meaningful distortion of GC content and dinucleotide frequencies spanning thousands of base pairs, as well as distortions of some DNA shape features and affinities to known TFs. We propose that these relatively far-reaching distortions are indicative of the 1D scanning, a coarse search mechanism that guides the TF towards a general region. Once a TF arrives to the vicinity of the target binding site, short-range 3D diffusion assumes a leading role in binding the target DNA site.

## Methods

### Data curation

For our analysis, we curated a set of human TF ChIP-seq experiments from the ENCODE portal [Luo et al., 2020]. We chose experiments targeting one of the following TFs: CTCF, FOXK2, IRF1, MEF2A, MYC, NANOG, NFRKB, RUNX1, SPI1, p53. These TFs were chosen (1) because of the sufficient data availability (ChIP-seq experiments available for multiple cell lines) and (2) because of the variability in both the structure of the DNA-binding domain and the TF functionality. For example, CTCF regulates the 3D organization of the genome [Kim et al., 2015]. MYC and NANOG are known pluripotency factors, regulating gene expression in embryonic stem cells [Povolotskii et al., 2025], while the IRF TFs activate the genes involved in immune response. Structurally, these TFs are different as well: no two TFs from our list share a DNA-binding domain protein family. The aforementioned CTCF is a zinc finger, while NANOG contains a homeodomain DNA-binding domain. MYC TF contains the helix-loop-helix DNA-binding domain [Kim et al., 2015, UniProt Consortium, 2018].

We limited our search to experiments without red flags in the audit category. From this set of experiments, we selected only the experiments associated with a biosample that appears at least three times (*i. e*., H1, K562, HepG2, A549, MCF-7, GM12878, WTC11 and GM23338). For each available combination of TF and biosample, we included a single experiment. In most cases, no more data was available.

As a representation of each experiment, we took the “conservative IDR thresholded peaks” file. If no such file was available, we represented the experiment with the “IDR thresholded peaks” file. There was only one such case (ENCFF692RPA, CTCF, H1 biosample). The files were downloaded on 17 January 2025. The final identifiers can be found in Supplementary Table S1.

To validate our results using a different experimental method, we included a Cut&Tag experiment GSM3560258 [Kaya-Okur et al., 2019] (9864 peaks). Using this data source, we replicated the peak calling process using MACS2 [Feng et al., 2012] as per the original work [Kaya-Okur et al., 2019].

### Feature extraction

From each ChIP-seq and Cut&Tag experiment, 10 000 peaks were uniformly randomly sampled (if there were fewer peaks, all were included). For each of the acquired peaks, its center was identified. Each of the peaks was then prolonged 5 000 bp both upstream and downstream of the peak center, resulting in a 10 000 bp–long patch of DNA sequence for each sampled peak. We denoted this patch a “prolonged peak”. We assume the binding site to be in the middle of a ChIP-seq peak [Gheorghe et al., 2018]. The construction of prolonged peaks is illustrated in Figure 1, steps A and B.

**Figure 1.**
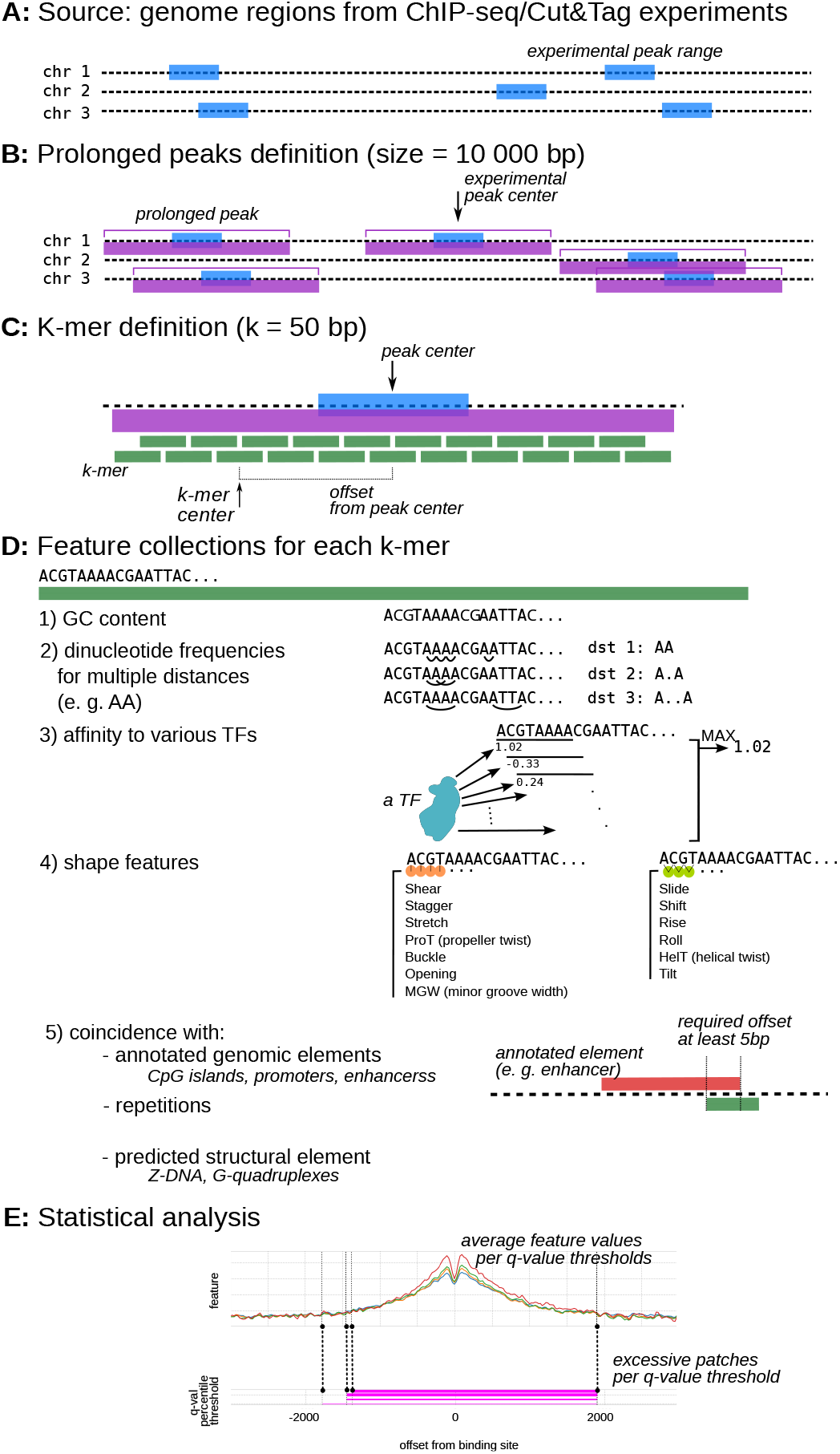
Illustration of feature extraction steps. An experimental peak range is indicated in blue, a prolonged peak in violet and a k-mer in green. **A:** illustration of peak coordinates as captured in a ChIP-seq or Cut&Tag experiment. **B:** definition of the prolonged peaks. The center of the experimental peak range is identified, then, a region spanning 5000 bp upstream and downstream is taken a the prolonged peak. **C:** a prolonged peak is then divided to k-mers (k=50 bp) which overlap for 25 bp (by half). To each k-mer, an offset value is associated as the distance from the original peak center to the k-mer center. **D:** various features are measured for each k-mer: GC content (number of G or C bases in the k-mer), dinucleotide frequencies, maximum TF affinity to the k-mer for every TF in the non-redundant set of human TFs from HOCOMOCO [Kulakovskiy et al., 2017], vector of shape features as measured by deepDNAshape [Li et al., 2024] and an indication on whether the k-mer intersects with one or multiple previously annotated genome elements such as promoters, enhancers or CpG islands (uses NIAID Visual & Medical Arts. (10/7/2024). NF-*κ*B. NIAID NIH BIOART Source: bioart.niaid.nih.gov/bioart/387). **E:** illustration of statistical analysis. In the top plot average feature value is plotted (in this case, GC content per k-mer) calculated in k-mers from peaks of Q-value above the indicated threshold. The bottom plot shows the corresponding excessive patch found in the one-dimensional cluster-based permutation test for each Q-value threshold (multiple patches can be found). In the following work, we depict excessive patches linked with an increase in feature value with respect to the neutral edge in magenta. We depict those linked with decrease in teal (not depicted here).

From each prolonged peak, we extracted 50 bp windows that span the entire length of the prolonged peak, with each window overlapping the previous one by 25 bp. Separation into 50 bp–long overlapping k-mers is illustrated in Figure 1, step C. To each of the k-mers, we assign the following values (each is illustrated in Figure 1, step D):

- **GC content**: the number of guanine (G) and cytosine (C) bases in the k-mer (Figure 1, D1).
- **Frequencies of dinucleotides**: the number of a particular dinucleotide (*i. e*. CG or AC) in the window sequence. We probe the occurences of neighboring dinucleotides as well as those separated by one to six arbitrary base (Figure 1, D2, inspired by the approach of Povolotskii et al.).
- **TF affinity information**: given a TF, we calculate its maximum possible affinity to a k-mer. We calculate the affinity value by sliding the k-mer sequence by the HOCOMOCO scoring model of the TF (both the forward sequence and its reverse complement). The maximum affinity seen in the k-mer is then kept (illustrated in Figure 1, D3). We calculate this value for every human transcription factor from the HOCOMOCO v12 non-redundant set [Kulakovskiy et al., 2017].

To verify if the affinity to various TFs from the HOCOMOCO datasets changes around the binding site, the affinity score is normalized with regards to affinity to sequences 5000 bases upstream of annotated transcription starts of RefSeq genes (hg19 genome assembly, downloaded from UCSC Genome Browser database [Perez et al., 2025], hgdownload.soe.ucsc.edu/goldenPath/hg19/bigZips/). We will refer to these affinity values as “upstream-normalized”.

Finally, in order to tell whether the affinity is changed beyond the change in GC content, we developed the following process: we randomly sample 500 k-mers at each offset from the original peak center. For a random permutation of each sampled k-mer the TF affinity is then calculated as above and used for z-score normalization. This way, we keep the GC content distribution as it changes with the proximity to the peak center. We will refer to these affinity values as “scramble-normalized”.

- **DNA shape features**: for each window, we predicted DNA shape features from the sequence using deepDNAshape [Li et al., 2024] (Figure 1, D4). For prediction, we set the number of layers used by deepDNAshape to 7. The prediction on the window edges were excluded so that every prediction has the context sequence available. The calculated DNA shape features can be classified to two groups: first, features that correspond to a single base pair (shear, stagger, stretch, propeller twist (ProT), buckle, opening and minor groove width (MGW)). Secondly, shape features corresponding to two adjacent base pairs include slide, shift, rise, roll, helical twist (HelT) and tilt [Li et al., 2017].
- **intersection of a k-mer with repetitive genomic regions, predicted Z-DNA patches and/or predicted G-quadruplexes** (a yes/no indicator): coordinates of these features were gathered from non-B DB [Cer et al., 2012] (hg38 genome assembly, nonb-abcc.ncifcrf.gov/apps/ftp/). A k-mer is reported to intersect such a feature if it has at least a 5 bp overlap with it.
- **intersection of a k-mer with CpG islands** (a yes/no indicator). CpG island can be associated with promoter regions. The annotation was gathered from the UCSC genome annotation database (hg38, GRCh38 Genome Reference Consortium Human Reference 38 (GCA 000001405.15)), where these areas are defined as tracks longer than 200 bp, with over 50 % GC-content and containing more 0.6 times the expected number of CG dinucleotide assuming base independence (genome.ucsc.edu/cgi-bin/hgTrackUi?g=cpgIslandExt).
- **intersection of a k-mer with ENSEMBL regulatory features** (a yes/no indicator). For each k-mer, we report intersection with annotated regulatory features (enhancer, promoter, open chromatin region). The information on regulatory features was dowloaded from ENSEMBL [Zerbino et al., 2015], release 113. Again, a k-mer is reported to intersect such a feature if it has at least a 5 bp overlap with it.

### Statistical analysis

To identify areas with increased or decreased feature values in the prolonged peaks, we defined an “edge” of the prolonged peak as the first and last 500 bp of the prolonged peak. We assumed this “edge” to be neutral, without any sequence bias associated with the presence of the binding site. Then, we adopted a one-dimensional cluster-based permutation approach to identify continuous runs of significant sequence-feature deviation in each prolonged peak. To perform this analysis, we subtract this edge mean from each peak’s profile to yield a peak×offset matrix of deviations. A sparse adjacency matrix was constructed linking each offset to its immediate neighbors, and MNE’s permutation cluster 1samp test was applied (1,000 permutations; cluster-forming threshold —t— ¿ 2.0; two-sided) [Gramfort et al., 2013]. Clusters whose summed t-mass exceeded the 95th percentile of the permutation null distribution (p ¡ 0.05) were called significant, and their start/end offsets were recorded. We label these ranges as “excessive patches”, as they show an excess compared to what is observed in the “edge” (the neutral area).

As stated above, for each feature and for each ChIP-seq or Cut&Tag experiment, we first summarized the 10 kb prolonged peak as a series of equally spaced windows (50 bp with 25 bp overlap).

Then, we attempted to identify excessive patches in the individual prolonged peaks. For every feature and every window, we calculated the respective z-score using the “edges” defined above as background distribution. We declared a window/feature pair excessive if it had absolute z-score of more than a predefined threshold, gaining a zero-one vector for every feature. On this vector, we then applied rolling mean on every five windows. Based on this vector, we declared a window/feature pair excessive after smoothing if its value is bigger than 0.5, *i. e*. if there is at least 3 out of 5 window/feature pairs around the given window that have z-score higher than 3. We consider a window *w* excessive if there is any feature *f* for which window *w*/feature *f* pair is excessive after smoothing. This evaluation was used for visualization only (in Figure 2); we did not derive further inference beside simple demonstration.

**Figure 2.**
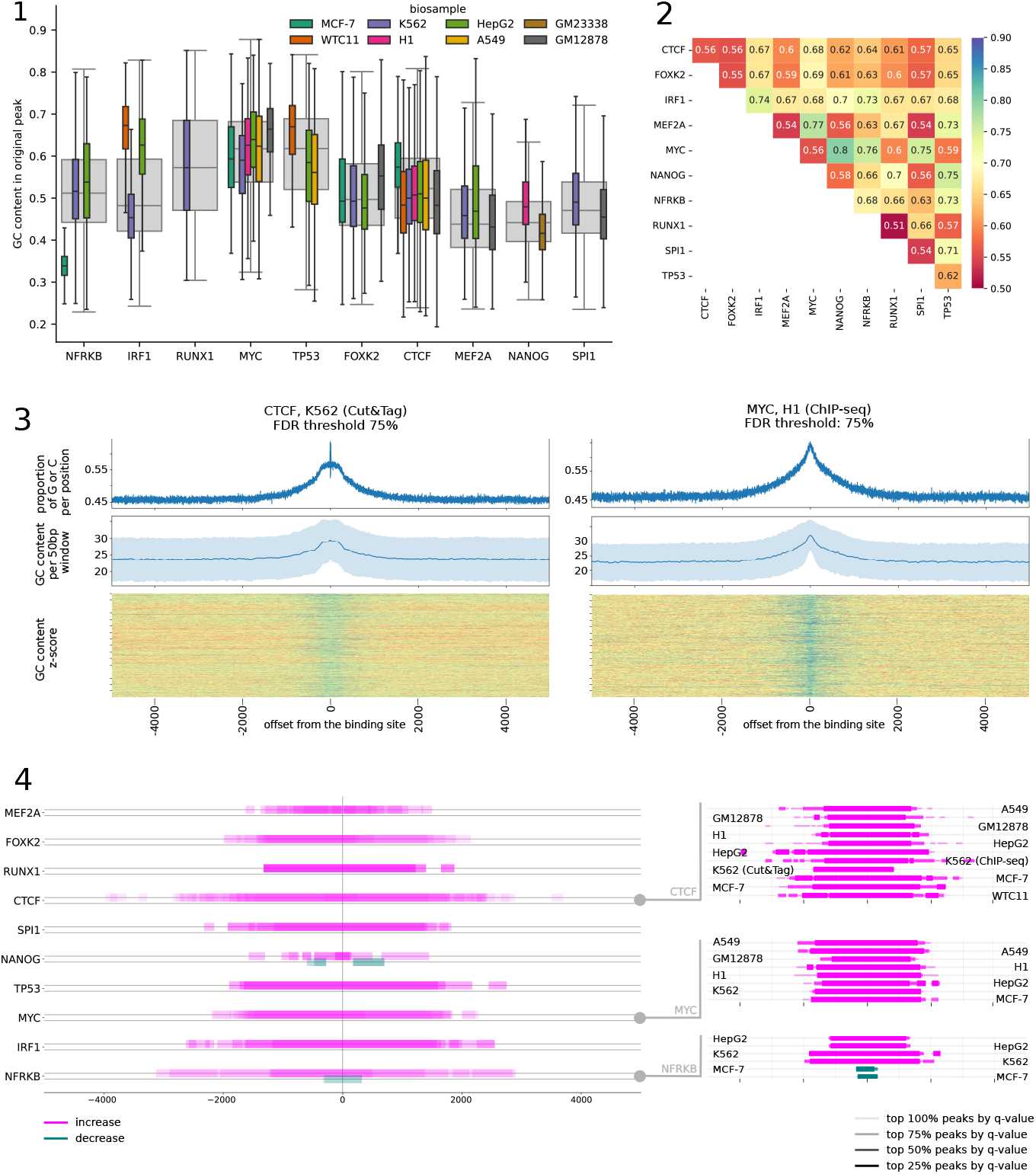
**(1):** Distribution of GC content in peaks identified by ChIP-seq experiment, grouped by cell line and target TF. The grey boxplot describes the distribution of GC content in all the peaks captured in a TF ChIP-seq experiment targeting the corresponding TF. The peak size is kept as found in the experiment. **(2):** Average accuracy of agglomerative clustering based on trivial GC content difference. On the left, grouped and averaged by target TF. On the right, grouped and averaged by cell line. For a pair of peak sets, we measured accuracy by uniformly sampling 1000 peak sequences from each file and clustering them using an agglomerative approach to put together sequences from the same experiment. The best outcome among several tested linkage functions (single, average, complete) was reported. **(3):** Overview of GC content on the most relevant 25 % of peaks in the Cut&Tag experiment targeting CTCF in K562 (left) and in the ChIP-seq experiment targeting MYC in H1 (ENCFF372RQZ, right). In each of these plots, the top plot depicts G or C base fraction per site. The middle plot shows average GC content per 50 bp window *±* its standard deviation. The bottom plot shows GC content z-score (edges were used as the background distribution) per window in individual prolonged peaks (calculation detailed in methods). **(4):** On the left, areas with significant increase or decrease (excessive patches) in GC content w. r. to the putative binding site (middle of the experimental peak) for the tested TFs, using the 50 % most relevant experimental peaks. Decrease is depicted in teal, increase in magenta. Color intensity is proportional to the number of experiments corresponding to the TF – if the color if fully opaque, the excess (increase or decrease) was seen in all experiments for the given TF. Detailed annotation on average patch sizes as well as the number of experiments displaying this effect can be found in Table 1. On the right, a detailed look for experiments targetting CTCF, MYC and NFRKB. Available experiments are depicted individually. Areas with significant increase or decrease (excessive patches) in GC content w. r. to the putative binding site are distinguished by line width and opacity for different FDR percentile thresholds.

We tried to estimate the effect of the immediate sequential context and position on the various shape parameters using a linear regression model using the *statsmodels* Python package [Seabold and Perktold, 2010]. Each shape feature was modeled as a function of sequential context (either a single base pair or a pair of neighboring base pairs), position relative to the binding site and their interaction. Using the *anova lm* function in the statsmodels package, we generated ANOVA tables for the fitted models and calculated effect size coefficient *η*^2^ as the ratio between the variable sum of squares and total sum of squares.

**Table 1.** Number of experiments in which an excessive patch was detected and average total size of the excessive patches, derived from the 50 % most relevant experimental peaks in the ChIP-seq/Cut&Tag experiments. The average size of covered area is calculcated as the mean area covered by excessive patches in the given excess direction. If there are no excessive patches found for a direction and experiment, it is not included in the average. Positioning of the excessive patches can be found in Figure 2.4.

|  |  |  |  |
| --- | --- | --- | --- |
| MEF2A | increase | seen in 5/5 experiments | avg. size $1.9 \cdot 10^3$ bp (s. d. $7.7 \cdot 10^2$ bp) |
|  | decrease | undetected |  |
| FOXK2 | increase | seen in 8/8 experiments | avg. size $2.8 \cdot 10^3$ bp (s. d. $8.9 \cdot 10^2$ bp) |
|  | decrease | undetected |  |
| RUNX1 | increase | seen in 2/2 experiments | avg. size $2.8 \cdot 10^3$ bp (s. d. $2.8 \cdot 10^2$ bp) |
|  | decrease | undetected |  |
| CTCF | increase | seen in 11/11 experiments | avg. size $4.2 \cdot 10^3$ bp (s. d. $1.4 \cdot 10^2$ bp) |
|  | decrease | undetected |  |
| SPI1 | increase | seen in 4/4 experiments | avg. size $3.3 \cdot 10^3$ bp (s. d. $6.0 \cdot 10^2$ bp) |
|  | decrease | undetected |  |
| NANOG | increase | seen in 4/4 experiments | avg. size $9.8 \cdot 10^2$ bp (s. d. $9.0 \cdot 10^2$ bp) |
| | decrease | seen in 2/4 experiments | avg. size $7.9 \cdot 10^2$ bp (s. d. $8.8 \cdot 10^1$ bp) |
| TP53 | increase | seen in 3/3 experiments | avg. size $3.7 \cdot 10^3$ bp (s. d. $5.9 \cdot 10^2$ bp) |
|  | decrease | undetected |  |
| MYC | increase | seen in 8/8 experiments | avg. size $3.4 \cdot 10^3$ bp (s. d. $6.8 \cdot 10^2$ bp) |
|  | decrease | undetected |  |
| IRF1 | increase | seen in 4/4 experiments | avg. size $3.9 \cdot 10^3$ bp (s. d. $6.9 \cdot 10^2$ bp) |
|  | decrease | undetected |  |
| NFRKB | increase | seen in 4/6 experiments | avg. size $4.0 \cdot 10^3$ bp (s. d. $2.1 \cdot 10^2$ bp) |
| | decrease | seen in 2/6 experiments | avg. size $6.4 \cdot 10^2$ bp (s. d. $1.8 \cdot 10^1$ bp) |

### Assessing biological relevance

In recent years, the representation of binding site sequences has gained significant nuance. To evaluate the long-range effects of DNA sequence context, we chose not to incorporate predefined motif models into our analysis. Instead, we relied on experimental evidence: assuming that ChIP-seq or Cut&Tag peaks provide a credible description of the TF’s binding landscape within the cell [Gheorghe et al., 2018, Worsley Hunt et al., 2014]. We assume the binding site to be at the center of the experimental peak [Gheorghe et al., 2018]. To ensure our long-range sequence-bias signals are not driven by low-confidence peaks, we ranked all binding events by their MACS2 FDR (q-value) and then re-ran the analysis above on increasingly stringent subsets: a dataset consisting of (1) the top 75%, (2) the top 50% and (3) the top 25% of peaks.

### Clustering of peaks based on GC content

For each peak sequence in the ChIP-seq experiment results, we assigned the GC content as the percentage of guanine or cytosine bases. No peak size normalization was used.

Then, for each pair of experiments, we uniformly randomly sampled 1000 peak sequences from each experiment and we tried to separate them using agglomerative clustering (scikit-learn implementation [Pedregosa et al., 2011]) and a trivial GC content difference distance function (see Results). For clustering, we try out single, complete and average linkage function and we report the result with the best accuracy. For each experiment pair, we repeat this process ten times.

### Assessing the presence of conserved sequence motifs by sequence alignment

As a control experiment, we constructed two multiple sequence alignments (MSA) for each BED file in our input dataset. For the first one, we randomly sampled 1000 peaks and took 150 bp upstream and downstream from each peak center (resulting in 300 bp sequences). The second one was constructed by sampling 100 prolonged peaks (see above). For alignment, we used the Clustal Omega command-line tool in default setting for DNA sequences [Sievers et al., 2011]. We will refer to these two alignments as the “MSA on peaks” and “MSA on prolonged peaks”.

### Combining prolonged peaks with information on chromatin accessibility

We also examined whether open chromatin data correlates with TF binding and other genomic features. To do that, we downloaded all available ATAC-seq experiments in the ENCODE database for any of the cell lines in out original dataset. For each cell line, we calculated the average ATAC-seq peak size and normalize all the peak to that size. Then, we then assessed, for each k-mer found in the prolonged peaks, whether it intersected with a normalized ATAC-seq peak (from the same cell line) by at least 5 base pairs. Such a k-mer was then labeled as ATAC+ k-mer, otherwise, we denote it as ATAC-k-mer. The list of all ATAC-seq experiments can be found in Supplementary Table S2

### Use of artificial intelligence in preparing this manuscript

We acknowledge the use of OpenAI’s ChatGPT in preparing this manuscript. It provided support in drafting simple code snippets, smoothing English phrasing, and offering basic editing suggestions. All scientific ideas, analyses and conclusions, however, are solely those of the authors.

## Results

As can be seen on the Figure 2.1, there is considerable variability in the nucleotide frequency of a peak captured in different ChIP-seq experiments depending on the target TF as well as the biosample used in the experiment. In particular, for NFRKB and IRF1 the difference between median peak GC content is over 10 % depending on which cell line was used in the experiment.

These differences in GC content are, in some cases, sufficient to distinguish between loci bound in two different experiments. We defined a trivial ‘distance’ function as

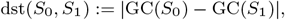

where GC(*S*) denotes the fraction of G or C bases in sequence *S*. Using this function, we were able to cluster the sequences using agglomerative clustering with up to 70–80 % accuracy (see Figure 2.2).

Beyond the GC content fraction, traditional bioinformatic methods reveal no similarity or consensus. In both MSA on peaks and MSA on prolonged peaks, we found little to no shared subsequences. The resulting MSAs have high gap fraction per positions (consistently above 70%). We were not able to retrieve distinct consensus motifs shared between peak sequences bound to the same TF in the same cell line, not even local ones.

### GC content reveals a differently distributed patch around TF binding site

First, we attempted to assess the size of the altered span. To do that, we used sequences spanning 10000 bp centered in the peak center (prolonged peaks). To gain the first insight, we centered all the peaks by the prolonged peak center (putative binding site) and calculated the fraction of G or C bases per position. Consistently, the naive GC proportion per position changes around the peak center. In most cases, it increases for approximately 10%, from 45% to 55% (demonstrated for CTCF and MYC in Figure 2.3, discussed below in depth). Complete set of figures showing GC proportion per position are provided in Supplementary Materials (Supplementary Figures S1 to S10).

**Figure 3.**
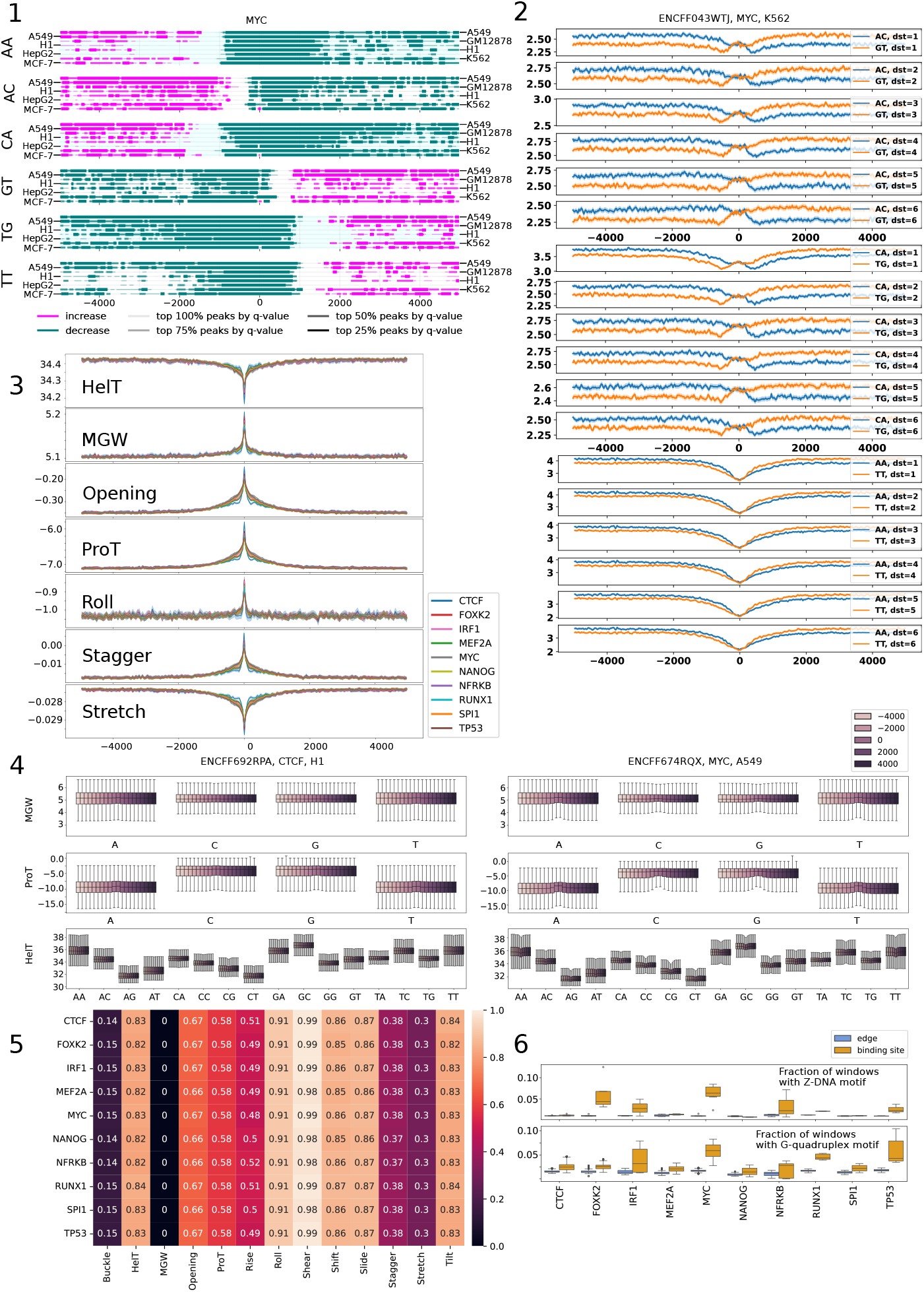
**(1):** Increase and decrease overview for average occurence of AA, AC, CA, GT, TG and TT dinucleotides (neighboring directly). Available experiments are depicted individually. Areas with significant increase or decrease (excessive patches) in GC content w. r. to the putative binding site are distinguished by line width and opacity for different FDR percentile thresholds (no threshold, 25%, 50%, 75%). Increase is depicted in magenta, decrease in teal. **(2):** Average occurence of AC and GT, CA and TG, AA and TT dinucleotides (respectively) with 95% confindence interval. Average dinucleotide frequency for the binding site as well as for the offsets of 1500 and 3000 bp upstream and downstream are presented in Supplementary Table S5.**(3):** Average values of selected shape features predicted by deepDNAshape per TF, with 95% confindence interval. The average shape values for the binding site as well as for the offsets of 1500 and 3000 bp upstream and downstream are presented in Supplementary Tables S6 to S12. **(4):** Distribution of minor groove width, propeller twist and hellical twist values by the sequential context and distance from the putative binding site for an experiment targeting CTCF (left) and for an experiment targeting MYC (right). **(5):** Average *η*^2^ effect size of sequential context acquired from an ordinary least squares model (statsmodels) of the given DNA shape feature as a linear combination of sequential context, position w. r. to the putative binding site and combination of the two. Full effect sizes can be found in Supplementary Figure S23.**(6):** Comparison of the frequency of k-mers incident with Z-DNA and G-quadruplex predicted motifs (non-B DB) at the binding site (offset w. r. to the binding site between −100 and 100) and at the edge of the prolonged peak (3000 bp and further from the binding site). For each experiment and group, the largest observed fraction of the respective DNA feature is taken.

We then proceeded with a more rigorous approach to estimate the extent of the affected region. Using the cluster-based permutation test (described in more detail in the Methods section), we found that the majority of experiments in our dataset show a consistent, statistically significant roughly symmetrical increase in GC content. This area can span around 2000–4000 bp (see Figure 2.4 and Table 1, detailed information in Supplementary Table S3).The only exception we were able to record were experiments targeting NANOG in GM233338 (where GC content around the binding site creates a “W”-shaped curve, being decreased 500 bp to 200 bp (both upstream and downstream) and “bouncing back” ±100 bp from the peak center) and experiments targeting NFRKB in MCF-7 (with a decrease of GC content ±300 bp from the peak center). It is worth noting that there are only minor differences between the patch sizes and excess direction (GC content increase vs. decrease) found in experiments on different cell lines, except for NFRKB (see bottom right plot in Figure 2.4). For this TF, there are immediately apparent differences: positions of increases and decreases directly contradict each other on a 1000 bp–long span.

**Figure 4.**
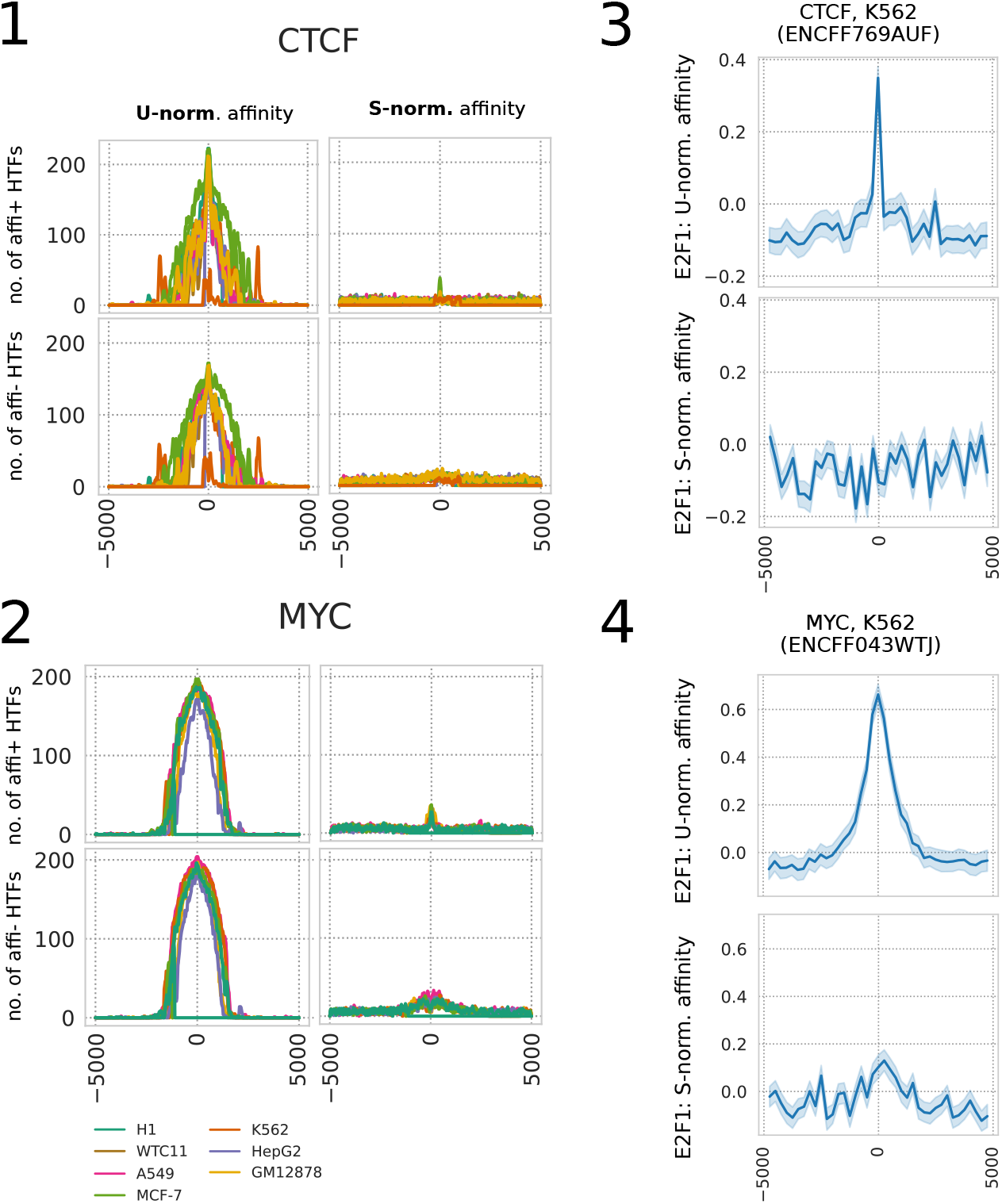
**(1)** and **(2)**: First row: counts of TFs from the HOCOMOCO (HTFs) with increased upstream-normalized (U-norm.) and scramble-normalized (S-norm.) affinity (denoted as affi+ HTFs in the figure). Second row: TF counts with decreased upstream-normalized and scramble-normalized affinity (denoted as affi-HTFs in the figure). For a TF from the HOCOMOCO database, we assess the normalized affinity increases and decreases using the cluster analysis (Methods). The upstream-normalized affinity score is calculated as the k-mer affinity z-score with regards to affinity to sequences 5000 bases upstream of annotated transcription starts of RefSeq genes (hg19). The scramble-normalized affinity score is calculated using a bootstrapped set of k-mers found at a given offset from the putative binding site. Each of these k-mers is then randomly shuffled. **(3)** and **(4)**: Average upstream-normalized affinity (top plot) and scramble-normalized affinity (bottom plot) of E2F1 to the top 25% most relevant sequences captured in experiments ENCFF769AUF and ENCFF043WTJ, respectively, complete with 95% confidence interval.

Notably, assessment of the prolonged peaks against ATAC-seq– defined open chromatin revealed little to no overlap in the case of NFRKB in the MCF-7 cell line (see Supplementary Figure S11.In other cases, most of the prolonged peaks coincide with open chromatin near the binding site. As the distance from the binding site increases, the proportion of ATAC+ k-mers declines, reaching a stable level around 1000 bp (for illustration, see Supplementary Figure S12). In some experiments, ATAC+ k-mers exhibit higher average GC content; however, this trend is not consistently observed across experiments targeting the same TF (see Supplementary Table S4).

It is worth noting that while the (average) GC content increases around the binding site in most of the observed cell lines, the absolute values of GC frequency often differ (only some CTCF binding sites appear interchangeable in terms of GC content near the binding site). We showcase this phenomenon in more detail in Supplementary Figures S13 and S14.

### Distribution of DNA dinucleotides around the binding site

An aspect commonly associated with promoter sites and transcription regulation is the enrichment of CG dinucleotides and CpG islands [Glass et al., 2007]. Near the analysed binding sites, we indeed observed excessive patches of increased CG dinucleotide frequency in the majority of tested experiments: we found it in all experiments but in NFRKB in MCF-7 and NANOG in GM23338. The sizes of excessive patches match the sized of GC content–linked excessive patches (2000–4000 bp, see Supplementary Figure S15). However, the incidence of binding sites with CpG islands varies with the target TF (see Table 2). We detected an increase of CG dinucleotide frequency even if we filtered out the prolonged peaks that intersect a CpG island.

**Table 2.**
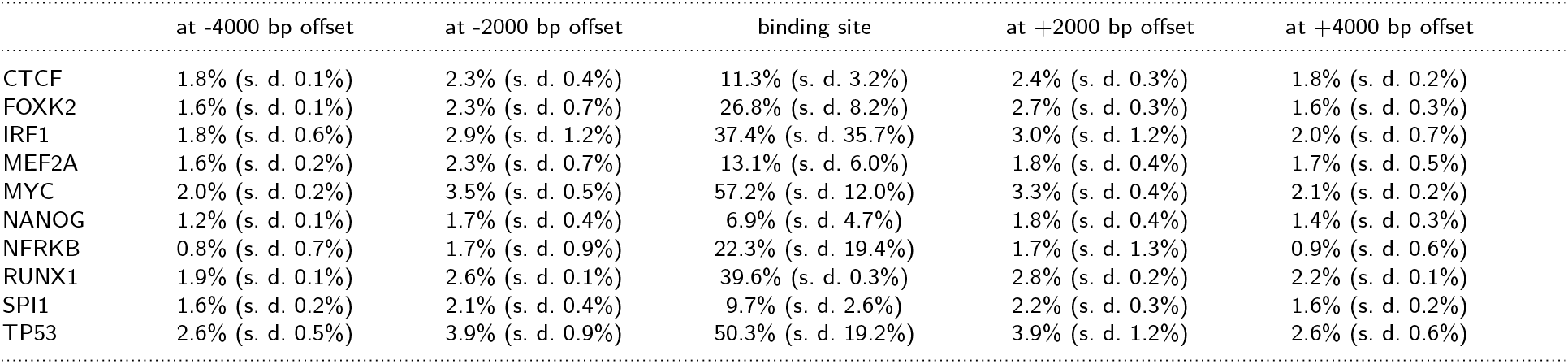
Average portion of k-mers (from all k-mers centered at a given offset from the binding site) that coincide with a CpG island at a given offset from the putative binding site for a given target TF. Standard deviation in parantheses.

While GC content around the binding site appeared roughly symmetrical in the studied experiments, it was not the case for dinucleotide frequency in general. The differences were most prominent in experiments targeting MYC, FOXK2, p53, NFRKB and IRF1. In particular, MYC binding sites are linked with an increase in AA, CA and AC dinucleotide frequencies, ending at −3000 – −1500 bp, followed by a decrease starting about −1000 bp (AA, CA) or about −300 bp (AC). This is mirrored downstream: there is a decrease in TT, TG and GT frequency starting far upstream of the binding site and ending 1000 bp (for TT and TG, 300 bp for GT) downstream of the binding site. Frequency of the GT dinucleotide starts rising 1000 bp from the binding site, frequency of TT and TG as well, 2000 bp from the binding site. The patches are depicted in Figure 3.1. The increases and decreases in the aforementioned dinucleotide frequencies for all tested TFs can be found in Supplementary Figures S16 to S21.This trend is present even for non-neighboring dinucleotides (Figure 3.2, Supplementary Table S5). Patches with increased AA, CA and AC frequency before the binding site and decreased downstream of it, combined with upstream decrease and downstream increase of their reverse complement TT, TG and GT were observed for FOXK2, p53, IRF1 and NFRKB in certain cell lines (not MCF-7).

MYC and p53 are involved in the cell cycle regulation [Bretones et al., 2015, Giono and Manfredi, 2006], while proteins from IRF family including the IRF1 has a role during the immune cell differentiation [Giang and La Cava, 2017]. We find it interesting to note that these TFs with very important roles for the cell fate share similar pattern in dinucleotide frequencies around the binding site. The AA/CA/AC frequencies mirrored by TT/TG/GT frequencies downstream appear to create a funnel around the binding site [Afek and Lukatsky, 2013b], pointing towards it. The binding of MYC has recently been shown to be influenced by the (larger) sequential context of the binding site [Povolotskii et al., 2025].

### Altered shape features around the putative binding site

DNA shape has been shown to affect the TF binding in several studies [Mathelier et al., 2016, Slattery et al., 2014, Chen et al., 2024]. We attempted to assess its role in the context of the prolonged peaks. To that end, we have predicted the DNA shape features using the deepDNAshape [Li et al., 2024] predictor, proccuring predictions for all features this tool offers us. These predictions were done for each 50 bp k-mer separately.

In Figure 3.3, we can clearly see that the average shape of propeller twist (summarized in detail in Supplementary Table S6), minor groove width (Supplementary Table S7), opening (Supplementary Table S8), stretch (Supplementary Table S9), stagger (Supplementary Table S10), roll (Supplementary Table S11) and helical twist (Supplementary Table S12) changes near the putatite binding site. In particular, the average stretch and helical twist decrease, while other parameters increase around the binding site. Increased roll has been associated with greater bendability of base pairs, while decreased helical twist reflects undertwisting of the DNA helix. Both features have been shown to contribute to a more flexible helical structure in shorter helices [Fujii et al., 2007]. Together with the increase in propeller twist [Yella et al., 2018] and other parameters, these parameters paint a picture of a more flexible DNA helix that is relatively more open to intercalators [Lindahl et al., 2017]. In line with the observed shifts in nucleotide composition, the change reaches its maximum at the peak center, indicating a possible link between the broader shape features and binding.

The minimal local sequence context surrounding the examined position in the DNA helix is linked with distinctly different DNA shape predictions for most of the queried shape features(demonstrated in Figure 3.4 for propeller twist and helical twist). Minor groove width is an exception here, as it showed only negligible differences in different (minimal) sequence contexts. To quantify the effect of sequence on the predicted DNA shape parameters rigorously, we trained simple linear regression models for each experiment and shape feature. The shape feature was treated as the dependent variable, with predictors including the immediate sequence context (the contacted base pair or the two nearest neighboring base pairs), the position relative to the binding site, and their interaction. Linear model performance metrics (e.g., coefficient of determination) are provided in Supplementary Figure S22.

Using these models, we summarized the importance of sequence context of the available shape features using effect size *η*^2^ coefficient. For the sequential context, the coefficient values can be found in Figure 3.5 (averaged by TF). All coefficients can be found in the extended visualization in Supplementary Figure S23The effect size coefficients calculated from a linear model (see Materials and methods) highlights strong influence of sequence context for all shape feature except minor groove width. On the other hand, the effect size coefficients of the relative position remain close to zero, suggesting that the position has minimal impact on the shape.

Cross-referencing the extended peaks with ATAC-seq data from the respective cell lines shows that (a) most k-mers near the in vivo binding site are located in open chromatin (ATAC+), and (b) the trends presented in Figure 3.5 remain consistent when ATAC+ and ATAC-k-mers are analyzed separately. The relationship between position relative to the putative binding site and DNA shape, stratified by ATAC+ and ATAC-status for each TF, is presented in Supplementary Figures S24 to S33.

In the section above, we have shown that the nucleotide composition near the *in vivo* binding site differs from surrounding regions. Using DNA shape predictions, we suggest that these compositional changes manifest as a change in DNA shape around the binding site, and that even single nucleotides or dinucleotides can strongly influence base pair shape parameters, compounding into the observed effects. In contrast, the effect of the position relative to the binding site appears to be minor (or almost negligible). The resulting helix shape appears to be more favorable for binding: a flexible, relatively flatter helix which is more accessible to intercalating agents.

### Non-B DNA conformers are predicted near *in vivo* binding site for several TFs

The results presented above are based on the assumption that DNA adopts a right-handed double helical structure. However, prior research suggests this may not always hold true. Several works have shown some involvement of Z-DNA conformation in gene regulation [CernÃ et al., 2004, Shin et al., 2016, Beknazarov et al., 2024]. Similarly, formation of G-quadruplexes was observed to have a role in transcription regulation [Robinson et al., 2021, Huppert and Balasubramanian, 2007]. For this reason, we chose to integrate available data on alternative DNA conformations into our analysis. The information is taken from the non-B DB database [Cer et al., 2012]. It is worth noting that, in the case of Z-DNA, the analysis is based on a sequence-driven thermodynamic prediction using the Z-Hunt-II program [Schroth et al., 1992]. Z-Hunt II predicts Z-DNA solely from the DNA sequence and may overpredict its occurrence, as additional cellular constraints probably influence Z-DNA formation [Beknazarov et al., 2020]. Instead, we interpret this prediction as the sequence’s propensity to adopt a Z-DNA conformation rather than as definitive evidence of the conformer. We consider this consistent with our other approaches, as the aim of this paper is to map sequence-based tendencies in the vicinity of the binding site.

In Figure 3.6, we compare the frequency of k-mers overlapping with a Z-DNA motif or a G-quadruplex motif (as predicted in the non-B DB) in the immediate vicinity of the binding site (± 100 bp) and in distal parts of the prolonged peaks (located 3000 bp from the peak center in both directions).

Restricting ourselves to the top 25 % of peaks in each experiment, we confirm this phenomenon to be statistically significant for FOXK2, MYC and p53 (Mann-Whitney U test, significance level of 5 %, one-sided alternative hypothesis: are the fractions of windows with a Z-DNA motif higher at the binding site (± 100 bp) than on the edge of the prolonged peak?). On average, about ≈ 1% of windows on the edge of the prolonged peak contain Z-DNA motif. At the binding site, it is 2 % (p53) or 6 % (MYC, FOXK2). The highest proportion of k-mers intersecting a predicted Z-DNA patch is consistently observed at the binding site, with this proportion decreasing symmetrically as the distance from the site increases. However, even for these TFs, the change is too small to fully explain the nucleotide frequencies reported above.

For G-quadruplex motifs, we found that CTCF, FOXK2, MEF2A, MYC and p53 are more likely to have a G-quadruplex at the binding site than at the edge of the binding site (Mann-Whitney U test, significance level of 5 %, one-sided alternative hypothesis: are the fractions of windows with a G-quadruplex motif higher at the binding site?). Again, the magnitude of the increase is fairly small: from ≈1.5 % to 3–6 % on average. After cross-linking the peaks with respective ATAC-seq experiments and keeping only offsets that are ATAC+/ATAC-at least from 25 %, a statistically significant difference can only be observed for MYC binding sites.

### Transcription factor affinity around the peak center

To gain insight into possible cooperation of other TFs with the target TF in the given experiment, we scored the prolonged peaks with affinity models of more than 400 TFs from the HOCOMOCO database [Kulakovskiy et al., 2017]. For each window and HOCOMOCO TF combination, we work with the maximum affinity (or maximum affinity z-score) per window. The results reported below were acquired by processing the top 25% prolonged peaks in each experiment.

When assessing the upstream-normalized affinity of TFs from the HOCOMOCO database to the prolonged peaks, we observed that in the binding site vicinity, most TFs exhibited either increased or decreased affinity relative to the edges of the prolonged peaks. In most experiments, few to no TFs were left without changes in affinity by the binding sites. However, when we accounted for the relationship between GC content and position from the putative binding site by using scramble-normalization (see Methods), little HOCOMOCO TFs showed change (approximately 20–30 out of more than 400). This is depicted for CTCF and MYC in Figure 4.1 and 2 and in full in Supplementary Figure S34.More than 150 HOCOMOCO TFs show increased upstream-normalized affinity in several CTCF-targeting experiments, in all but one MYC-targeting experiment, and in nearly all experiments across the tested dataset (see Supplementary Figure S34). In contrast, an increase in scramble-normalized affinity was never observed for more than 30 HOCOMOCO TFs within any single experiment. The same trend was observed for decrease in normalized affinity scores. It appears that for most TFs from the HOCOMOCO database, the observed changes in affinity around the binding site can be largely attributed to variations in GC content.

As a next step, we evaluated whether the increase in normalized affinity is consistently observed across experiments targeting the same TF. We compared the sets of HOCOMOCO TFs that showed increased or decreased affinity scores at specific positions across pairs of experiments. For the majority of experiment pairs targeting CTCF, FOXK2, MYC, SPI1, and TP53, the same TFs exhibited consistent patterns of increased or decreased upstream-normalized affinity. In contrast, greater variation was observed for NANOG and NFRKB, likely due to higher variability in nucleotide composition across the different cell lines in which these TFs were studied. Results are summarized in Supplementary Figure S35.

Only a handful of TFs are shared in all/majority of the comparisons. These are listed in Supplementary Table S13for scramble-normalized affinity scores. The observation that these TFs exhibit altered affinity even after controlling for positional GC content across multiple experiments suggests potential interactions with the target TF; however, further research is needed to confirm this.

## Discussion

As already shown by Dror et al., GC content changes near TF binding sites. In most cases presented above, we detect a statistically significant elevation in average GC content spanning several thousand base pairs. This elevation is largely consistent in size and position relative to the binding site between different cell lines, even though the absolute values of GC frequencies may differ. Although GC-rich regions are thermodynamically more stable [Vinogradov, 2003], our shape predictions suggest that elevated GC content coincides with increased propeller twist values, which have been associated with local structural adaptability rather than rigidity (at least in shorter contexts) [Yella et al., 2018].

Beyond the overall GC enrichment, for MYC and several other TFs listed above, the distribution of dinucleotide frequencies exhibits a pronounced directional asymmetry around the binding site. This arrangement resembles a funnel in sequence space centered almost perfectly at the binding site. Given that sequence tracts like poly(dA:dT) and poly(dC:dG) have been observed to modulate TF affinity to the DNA [Afek and Lukatsky, 2013a], it is plausible that the broader sequence composition patterns contribute to a similar effect: ‘guiding’ the TF along the helix. We have also observed that GC content affects the affinity of several TFs (most of human TFs from the HOCOMOCO database) to the binding site, even if slightly. Our findings suggest that GC content may act as a low-specificity low-strength filtering mechanism guiding potential cofactors, a constraint that could be overridden by sufficiently high TF concentrations [Kribelbauer et al., 2019].

Altogether, our predictions of the DNA shape from the sequence show systematic alterations that result in a flatter, “intercalation-prone” helix. There seems to be little qualitative differences between different target TFs. Our analysis also indicates that the local sequence composition drives the DNA shape, preconfiguring the DNA into conformations that are presumably more conducive to binding. In our opinion, the inherent properties of DNA sequences are likely further modulated by regulatory mechanisms within the cell nucleus. Cross-referencing ChIP-seq with ATAC-seq data shows that most of the excessive patches correspond to open chromatin in the respective cell lines. On the other hand, these results should be interpreted with caution, as they are based solely on DNA shape predictions derived from sequence. We took several steps to ensure the validity of our analyses: shape was predicted for individual k-mers in isolation, with the model having no access to sequence context beyond the k-mer boundaries; positions were analyzed separately and never pooled (we only pooled values for the exactly same position from different prolonged peaks); and sequence-based analyses were limited strictly to the base pair under investigation, to avoid artificially replicating the prediction input. Despite these limitations, the recurrence of these patterns across several TFs and cell lines, along with supporting ATAC-seq evidence showing that the surrounding regions are largely open chromatin, points to an underlying biological mechanism by which DNA sequence encodes structural preparedness for binding. Another open question is the extent to which these shape preferences contribute to the observed effects. With the currently available data, we cannot assess their relative importance, and it is possible that DNA shape plays only a supporting role.

Even slight changes in affinity could influence the 1D scanning dynamics, subtly guided by the underlying nucleotide composition. Even though individual protein–DNA electrostatic interactions are strongly screened in the ionic environment of the nucleus — falling below 1% of their vacuum strength within 2.5–3.6 nm (approximately 7–11 bp) and below 0.1% by 3.8–5.4 nm (approximately 11– 16 bp) — sequence-encoded structural distortions extend their effective influence to much longer genomic distances: elevated GC content, directional dinucleotide biases, and subtle DNA shape changes generate broad patches spanning hundreds to thousands of base pairs around binding sites, reshaping groove geometry, DNA flexibility, electrostatic multipole moments, and the local electrostatic potential [Gordân et al., 2013]. In this modified landscape, TFs undergo facilitated diffusion effectively travel along low-free-energy “highways” and “funnels” that bias their sliding and hopping trajectories, extending their functional communication range well beyond the nominal Debye length [Bacher et al., 2004, Halford and Marko, 2004, Slutsky and Mirny, 2004]. We propose that the altered nucleotide composition observed around binding sites reflects precisely such a mechanism – a ‘statistical funnel’ guiding the TF toward a ‘beacon’ at the binding site.

While our findings are consistent with a model of a ‘funnel’ that points to a ‘beacon’ at the binding site, we view them more as hypothesis-generating rather than definitive; experimental validation will be necessary in future work. There are several significant features in the cellular nucleus that were omitted in this study, such as nucleosome post-translational modification etc.

## Conclusion

Our results suggest that the broader sequence environment around *in vivo* binding sites (±1500 bp) carries information relevant to TF binding, as reflected by systematic shifts in nucleotide composition. These patterns may be partly mediated by DNA shape, as indicated by predicted shape features. Integration with ATAC-seq indicates that the regions analyzed are largely open chromatin in the corresponding cell lines, supporting the relevance of sequence-based shape modeling. We hypothesize that such compositional/structural biases could influence 1D sliding, guiding TF search dynamics toward the binding site in a ‘funnel’-line manner. In addition, GC content appears to drive broad changes in predicted affinity for many HOCOMOCO TFs. This is consistent with a weak, low-specificity filter that could potentially be modulated by the concentration of the potential cofactor TF.

## Abbreviations

(TFs): transcription factors

## Data availability

Source codes, complete list of used ChIP-seq experiments and final description of excessive patches for each experiment is stored in Zenodo archive under DOI 10.5281/zenodo.17900173.

## Author contributions statement

K.F. and J.V. conceived the experiment, K.F. conducted the computational experiments and analysed the results with input from J.Š.. K.F., J.Š. wrote and reviewed the manuscript with input from J.V..

## Acknowledgements

The authors thank the anonymous reviewers for their valuable suggestions. Computational resources were provided by the e-INFRA CZ project (ID:90254), supported by the Ministry of Education, Youth and Sports of the Czech Republic. This work was supported by the ELIXIR CZ Research Infrastructure under Grant (ID LM2023055, MEYS CR); and Institute of Organic Chemistry and Biochemistry of the Czech Academy of Sciences under Grant (RVO: 61388963).

