## Supplementary materials for "Non-consensus flanking sequence of hundreds of base pairs around in vivo binding sites: statistical beacons for transcription factor scanning"

### Supplementary Note S1: Materials and methods

**Table S1.** List of ChIP-seq experiments used, together with the target transcription factor, biosample identification and the corresponding number of peaks. If more peaks was available, the chosen 10 000 peaks are sampled randomly.

| ENCODE file ID | ENCODE experiment ID | Target TF | Biosample name | Number of peaks |
|---|---|---|---|---|
| ENCFF966KGI | ENCSR000BHV | CTCF | A549 | 10000 |
| ENCFF536RGD | ENCSR000AKB | CTCF | GM12878 | 10000 |
| ENCFF017XLW | ENCSR000AKB | CTCF | GM12878 | 10000 |
| ENCFF692RPA | ENCSR000AMF | CTCF | H1 | 10000 |
| ENCFF252PLM | ENCSR000BIE | CTCF | HepG2 | 10000 |
| ENCFF123DAC | ENCSR000BIE | CTCF | HepG2 | 10000 |
| ENCFF769AUF | ENCSR000AKO | CTCF | K562 | 10000 |
| ENCFF962IZR | ENCSR000DMO | CTCF | MCF-7 | 10000 |
| ENCFF820BVW | ENCSR000DMO | CTCF | MCF-7 | 10000 |
| ENCFF503DSN | ENCSR992XTY | CTCF | WTC11 | 10000 |
| ENCFF855EOH | ENCSR861JUQ | FOXK2 | GM12878 | 4761 |
| ENCFF578XIP | ENCSR861JUQ | FOXK2 | GM12878 | 4949 |
| ENCFF008HBF | ENCSR171FUX | FOXK2 | HepG2 | 8460 |
| ENCFF997KQI | ENCSR171FUX | FOXK2 | HepG2 | 7652 |
| ENCFF022HGL | ENCSR302AWT | FOXK2 | K562 | 10000 |
| ENCFF154SRU | ENCSR302AWT | FOXK2 | K562 | 10000 |
| ENCFF583QAD | ENCSR465VLK | FOXK2 | MCF-7 | 937 |
| ENCFF356XXS | ENCSR465VLK | FOXK2 | MCF-7 | 1028 |
| ENCFF174QAR | ENCSR890DSP | IRF1 | HepG2 | 2802 |
| ENCFF019LOH | ENCSR854MCV | IRF1 | K562 | 10000 |
| ENCFF251ZSJ | ENCSR854MCV | IRF1 | K562 | 10000 |
| ENCFF910ZUK | ENCSR381IES | IRF1 | WTC11 | 4632 |
| ENCFF131JIX | ENCSR000BKB | MEF2A | GM12878 | 10000 |
| ENCFF775DWU | ENCSR000BKB | MEF2A | GM12878 | 10000 |
| ENCFF987BRO | ENCSR291MJH | MEF2A | HepG2 | 6412 |
| ENCFF288TSN | ENCSR000BNV | MEF2A | K562 | 5016 |
| ENCFF190IHG | ENCSR000BNV | MEF2A | K562 | 996 |
| ENCFF674RQX | ENCSR000DYC | MYC | A549 | 9031 |
| ENCFF356PWE | ENCSR000DYC | MYC | A549 | 8813 |
| ENCFF765CKK | ENCSR000DKU | MYC | GM12878 | 2239 |
| ENCFF372RQZ | ENCSR000EBY | MYC | H1 | 5205 |
| ENCFF700CXD | ENCSR000EBY | MYC | H1 | 4987 |
| ENCFF800JFG | ENCSR000DLR | MYC | HepG2 | 1616 |
| ENCFF043WTJ | ENCSR744JJU | MYC | K562 | 7275 |
| ENCFF149BIS | ENCSR000DMP | MYC | MCF-7 | 8306 |
| ENCFF897LBK | ENCSR061DGF | NANOG | GM23338 | 8269 |
| ENCFF463AFM | ENCSR061DGF | NANOG | GM23338 | 9475 |
| ENCFF655XZF | ENCSR000BMT | NANOG | H1 | 5713 |
| ENCFF722JFZ | ENCSR000BMT | NANOG | H1 | 5560 |
| ENCFF252JYE | ENCSR145BHD | NFRKB | HepG2 | 1590 |
| ENCFF748JYG | ENCSR145BHD | NFRKB | HepG2 | 1814 |
| ENCFF594KPG | ENCSR657EOF | NFRKB | K562 | 10000 |
| ENCFF048HYG | ENCSR657EOF | NFRKB | K562 | 10000 |
| ENCFF360OXK | ENCSR248IMH | NFRKB | MCF-7 | 757 |
| ENCFF768VQG | ENCSR248IMH | NFRKB | MCF-7 | 840 |
| ENCFF264QLP | ENCSR414TYY | RUNX1 | K562 | 3182 |
| ENCFF015ADG | ENCSR414TYY | RUNX1 | K562 | 3593 |

| | | | | |
|---|---|---|---|---|
| ENCFF881ARS | ENCSR000BGQ | SPI1 | GM12878 | 10000 |
| ENCFF198SPL | ENCSR000BGQ | SPI1 | GM12878 | 10000 |
| ENCFF185DGL | ENCSR000BGW | SPI1 | K562 | 10000 |
| ENCFF456BUE | ENCSR000BGW | SPI1 | K562 | 10000 |
| ENCFF488WQN | ENCSR112XUO | TP53 | A549 | 5702 |
| ENCFF701DTE | ENCSR980EGJ | TP53 | HepG2 | 7662 |
| ENCFF280RDQ | ENCSR760GZL | TP53 | WTC11 | 7866 |

**Table S2.** List of ATAC-seq experiments used, together with the target biosample identification and the corresponding number of peaks.

| ENCODE file ID | ENCODE experiment ID | Biosample name | Number of peaks |
|---|---|---|---|
| ENCFF653FJA | ENCSR139OYS | A549 | 211686 |
| ENCFF899OMR | ENCSR032RGS | A549 | 278564 |
| ENCFF665IBA | ENCSR074AHH | A549 | 206786 |
| ENCFF139XJH | ENCSR265ZXX | A549 | 186872 |
| ENCFF735UWS | ENCSR220ASC | A549 | 204227 |
| ENCFF232BHQ | ENCSR288YMH | A549 | 210062 |
| ENCFF748UZH | ENCSR637XSC | GM12878 | 277999 |
| ENCFF470YYO | ENCSR095QNB | GM12878 | 223873 |
| ENCFF567ZCX | ENCSR485TLP | GM23338 | 283236 |
| ENCFF439EIO | ENCSR291GJU | HepG2 | 279739 |
| ENCFF913MQB | ENCSR042AWH | HepG2 | 217283 |
| ENCFF282BIU | ENCSR962TDN | K562 | 95499 |
| ENCFF344PIT | ENCSR685YMG | K562 | 88200 |
| ENCFF988RER | ENCSR343TQX | K562 | 122812 |
| ENCFF243TXP | ENCSR644KPP | K562 | 115312 |
| ENCFF839NAB | ENCSR978HFU | K562 | 79800 |
| ENCFF200XYP | ENCSR091YVK | K562 | 142022 |
| ENCFF812SFF | ENCSR935JVI | K562 | 119745 |
| ENCFF558BLC | ENCSR483RKN | K562 | 203874 |
| ENCFF477IQJ | ENCSR966ZCL | K562 | 138982 |
| ENCFF586IMX | ENCSR179IAC | K562 | 77483 |
| ENCFF274UER | ENCSR934FTO | K562 | 108381 |
| ENCFF563WNA | ENCSR800RAH | K562 | 169113 |
| ENCFF084PBF | ENCSR509MYW | K562 | 119694 |
| ENCFF680FIO | ENCSR793YFK | K562 | 124858 |
| ENCFF703WCG | ENCSR165JXS | K562 | 150255 |
| ENCFF145AEP | ENCSR761CBY | K562 | 110274 |
| ENCFF158GOJ | ENCSR243BXK | K562 | 105641 |
| ENCFF550TGR | ENCSR822IYG | K562 | 114219 |
| ENCFF529ZME | ENCSR121GEL | K562 | 106922 |
| ENCFF092PVG | ENCSR017LGQ | K562 | 137656 |
| ENCFF153CSM | ENCSR503EZG | K562 | 57092 |
| ENCFF990VZL | ENCSR803QIE | K562 | 101766 |
| ENCFF614WKT | ENCSR442DWQ | K562 | 113322 |
| ENCFF517QIJ | ENCSR342GFK | K562 | 109055 |
| ENCFF919ZPT | ENCSR582NPJ | K562 | 142156 |
| ENCFF651JKF | ENCSR021GTX | K562 | 68526 |
| ENCFF359EAN | ENCSR524ZSN | K562 | 85001 |
| ENCFF481SJK | ENCSR465IDZ | K562 | 106861 |
| ENCFF087HGO | ENCSR446UJL | K562 | 101869 |
| ENCFF226XER | ENCSR874GAJ | K562 | 78919 |
| ENCFF824NVT | ENCSR390UVH | K562 | 71350 |
| ENCFF637YWY | ENCSR199HSR | K562 | 110733 |
| ENCFF333TAT | ENCSR868FGK | K562 | 269800 |
| ENCFF842UZU | ENCSR956DNB | K562 | 123697 |
| ENCFF356SBB | ENCSR068MIW | K562 | 124886 |
| ENCFF601WRI | ENCSR259HTB | K562 | 105071 |
| ENCFF050STM | ENCSR096HVF | K562 | 112469 |
| ENCFF392DVU | ENCSR680XOP | K562 | 109171 |
| ENCFF339UPL | ENCSR304UWR | K562 | 67688 |
| ENCFF968ZUO | ENCSR632VDU | K562 | 81082 |
| ENCFF412STN | ENCSR281LEY | K562 | 110802 |
| ENCFF215XQJ | ENCSR180IJO | K562 | 56164 |

| | | | |
|---|---|---|---|
| ENCFF246XXM | ENCSR974QYY | K562 | 96675 |
| ENCFF797KDM | ENCSR214QZO | K562 | 115418 |
| ENCFF816LDC | ENCSR217QAB | K562 | 71100 |
| ENCFF527ASE | ENCSR214SFA | K562 | 102484 |
| ENCFF110ZGQ | ENCSR888JRM | K562 | 101726 |
| ENCFF763PRE | ENCSR197GOQ | K562 | 104552 |
| ENCFF962YNW | ENCSR741QNS | K562 | 80702 |
| ENCFF429LOY | ENCSR676UFY | K562 | 53737 |
| ENCFF603APA | ENCSR920YDG | K562 | 110912 |
| ENCFF100HTV | ENCSR420NIU | K562 | 88732 |
| ENCFF113CYL | ENCSR303NUP | K562 | 85264 |
| ENCFF141VWA | ENCSR737HKX | K562 | 114684 |
| ENCFF741UWF | ENCSR163VQD | K562 | 85780 |
| ENCFF055NNT | ENCSR859USB | K562 | 99862 |
| ENCFF252OYM | ENCSR314EDO | K562 | 123284 |
| ENCFF291IVT | ENCSR522ALT | K562 | 141367 |
| ENCFF415FXA | ENCSR296YAS | K562 | 84048 |
| ENCFF984BJB | ENCSR508LLL | K562 | 61571 |
| ENCFF239GPE | ENCSR745KUZ | K562 | 120757 |
| ENCFF520WSX | ENCSR005BVE | K562 | 125390 |
| ENCFF449NMH | ENCSR366CEQ | K562 | 128960 |
| ENCFF891XBE | ENCSR548GGF | K562 | 87188 |
| ENCFF821OEF | ENCSR422SUG | MCF-7 | 248668 |
| ENCFF524JAA | ENCSR089KIJ | WTC11 | 197596 |
| ENCFF321VDH | ENCSR541KFY | WTC11 | 264234 |

### Supplementary Note S2: Results

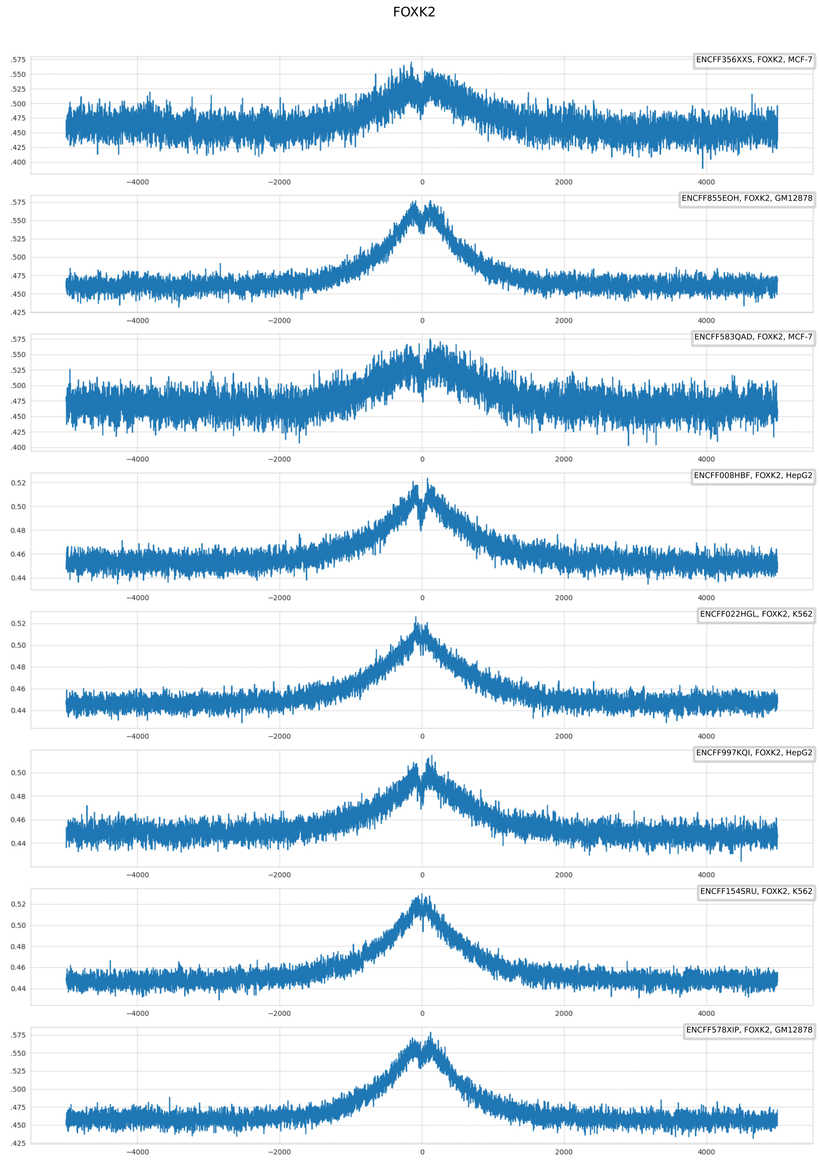


**Fig. S1.** Relation between GC content and distance from the putative FOXK2 binding site. On the y-axis, proportion of G or C base on given position can be found, while on the x-axis, number of base pairs from the putative binding site (ChIP-seq peak center).

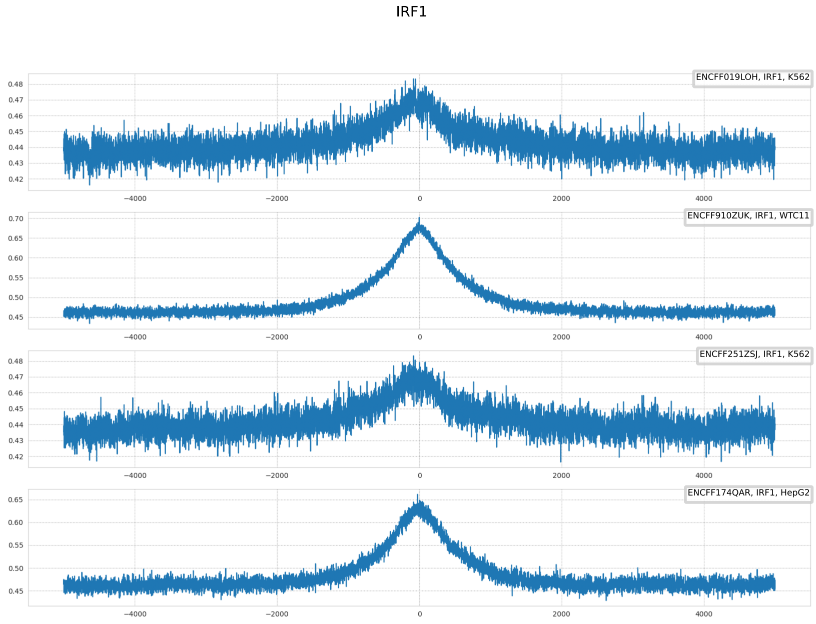


**Fig. S2.** Relation between GC content and distance from the putative IRF1 binding site. On the y-axis, proportion of G or C base on given position can be found, while on the x-axis, number of base pairs from the putative binding site (ChIP-seq peak center).

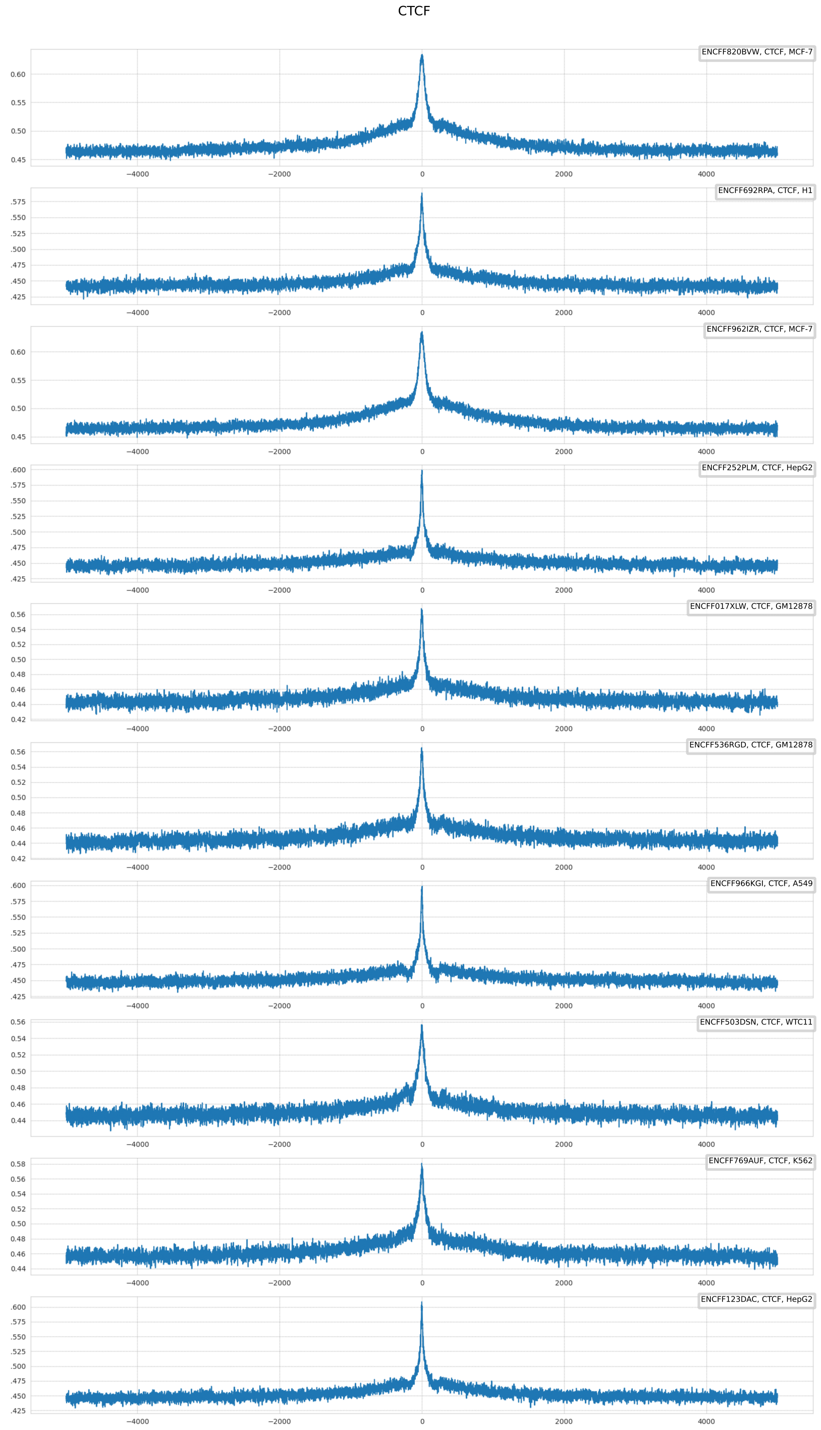


**Fig. S3.** Relation between GC content and distance from the putative CTCF binding site. On the y-axis, proportion of G or C base on given position can be found, while on the x-axis, number of base pairs from the putative binding site (ChIP-seq or Cut&Tag peak center).

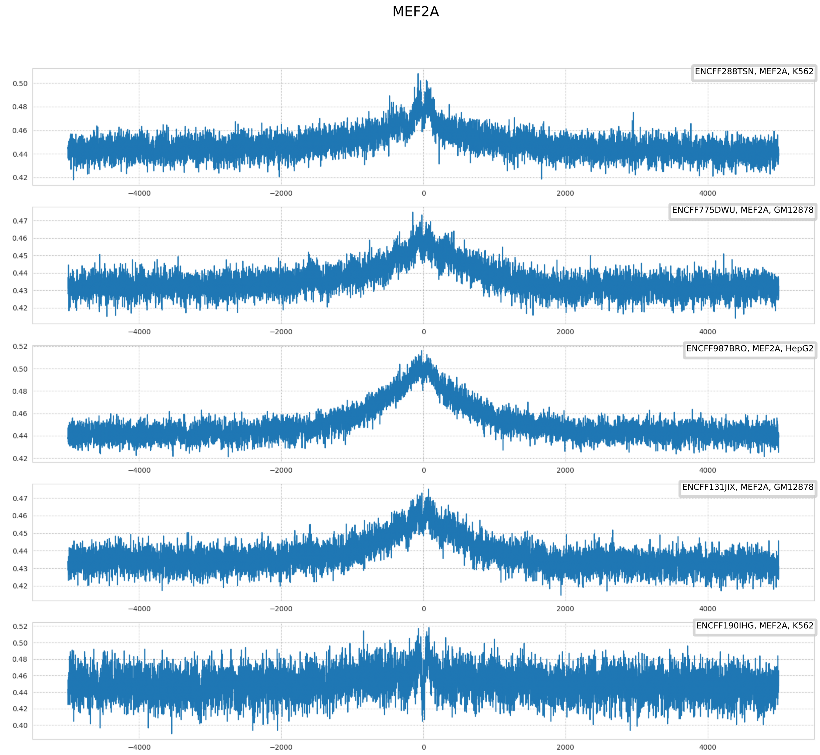


**Fig. S4.** Relation between GC content and distance from the putative MEF2A binding site. On the y-axis, proportion of G or C base on given position can be found, while on the x-axis, number of base pairs from the putative binding site (ChIP-seq peak center).

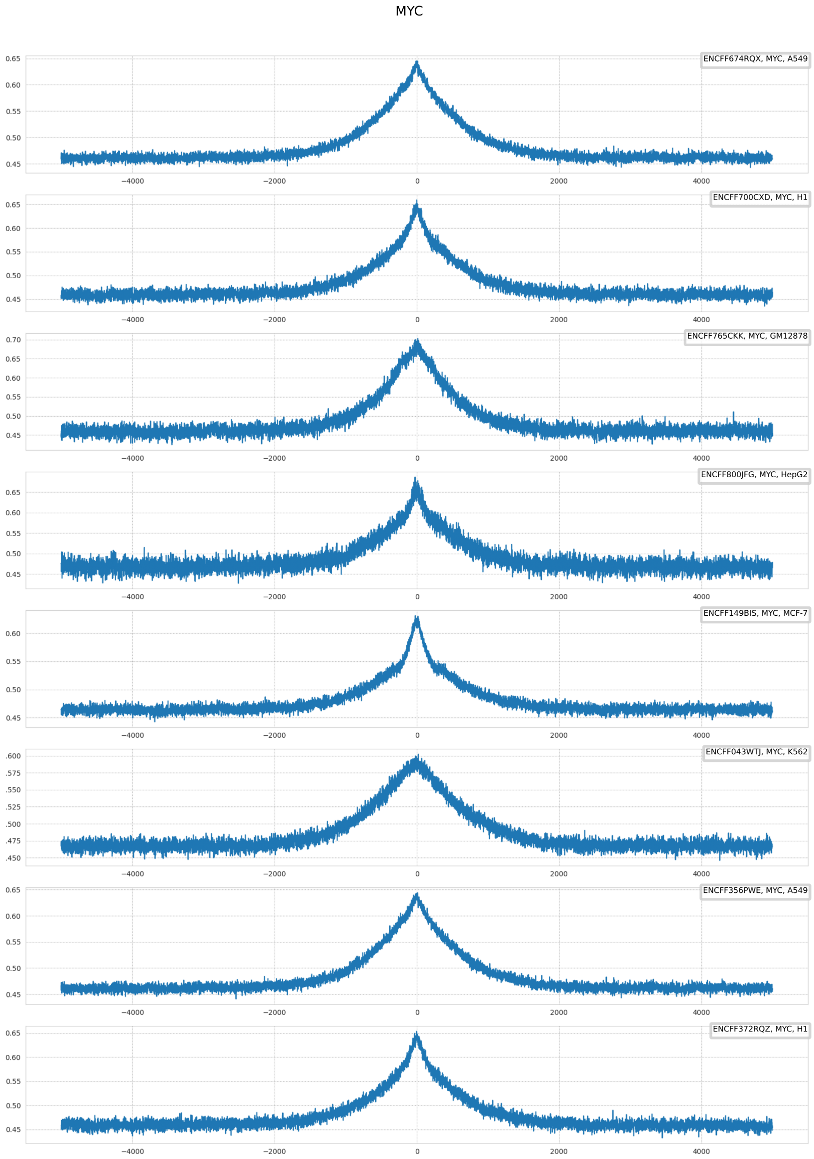


**Fig. S5.** Relation between GC content and distance from the putative MYC binding site. On the y-axis, proportion of G or C base on given position can be found, while on the x-axis, number of base pairs from the putative binding site (ChIP-seq peak center).

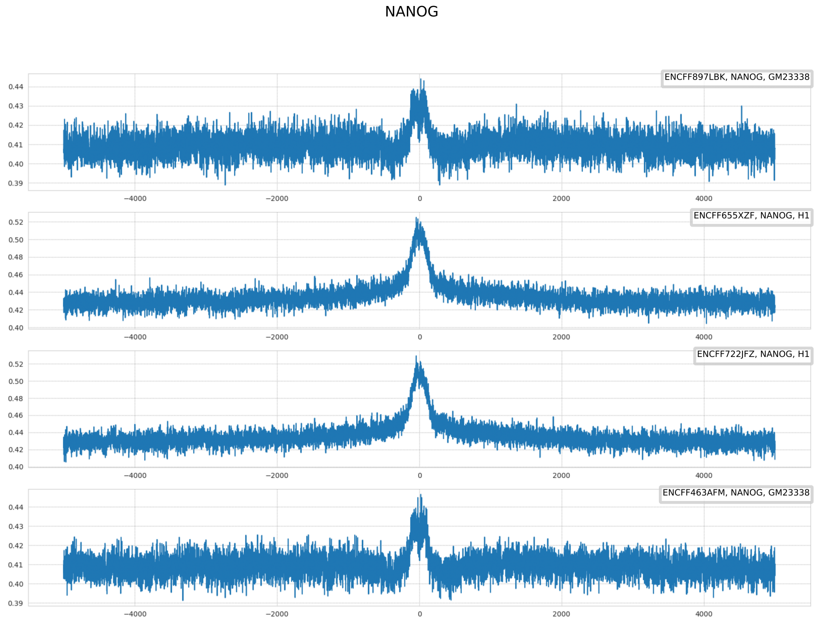


**Fig. S6.** Relation between GC content and distance from the putative NANOG binding site. On the y-axis, proportion of G or C base on given position can be found, while on the x-axis, number of base pairs from the putative binding site (ChIP-seq peak center).

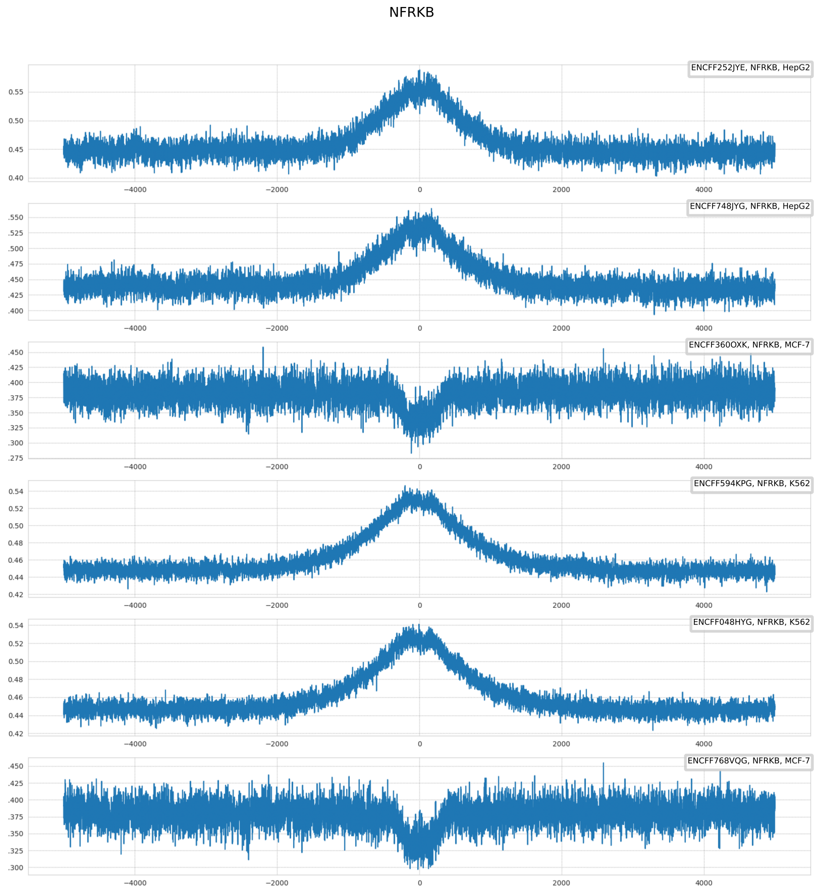


**Fig. S7.** Relation between GC content and distance from the putative NFRKB binding site. On the y-axis, proportion of G or C base on given position can be found, while on the x-axis, number of base pairs from the putative binding site (ChIP-seq peak center).

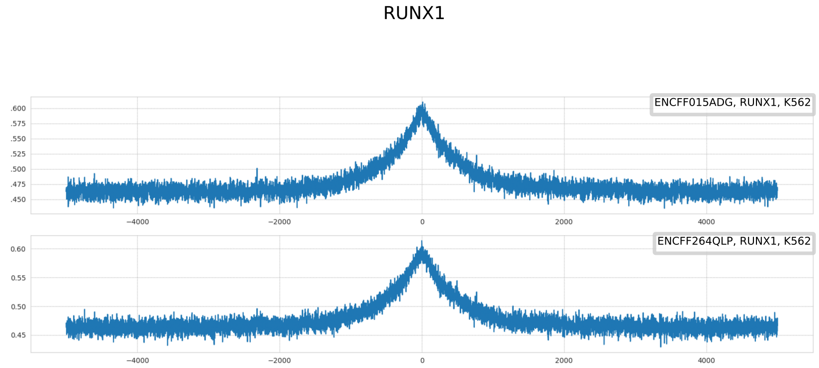


**Fig. S8.** Relation between GC content and distance from the putative RUNX1 binding site. On the y-axis, proportion of G or C base on given position can be found, while on the x-axis, number of base pairs from the putative binding site (ChIP-seq peak center).

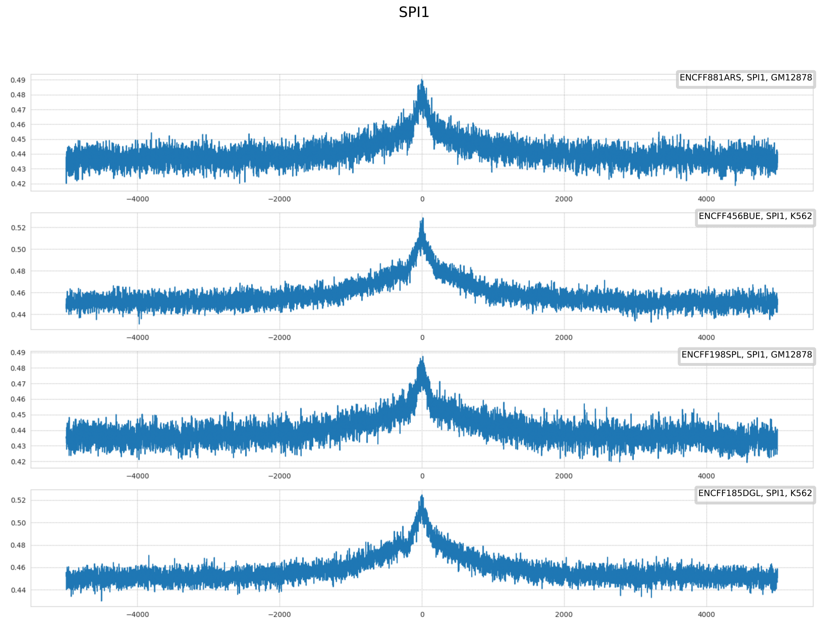


**Fig. S9.** Relation between GC content and distance from the putative SPI1 binding site. On the y-axis, proportion of G or C base on given position can be found, while on the x-axis, number of base pairs from the putative binding site (ChIP-seq peak center).

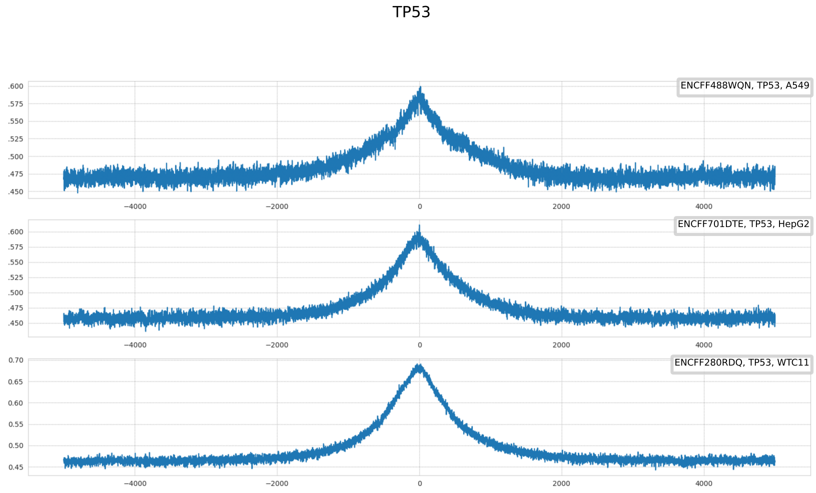


**Fig. S10.** Relation between GC content and distance from the putative TP53 binding site. On the y-axis, proportion of G or C base on given position can be found, while on the x-axis, number of base pairs from the putative binding site (ChIP-seq peak center).

**Table S3.** Coordinates (base pair offset w. r. to peak center) of the excessive patches in GC content identified by cluster analysis (see Materials and methods, Statistical analysis).

| Excessive patch start | Excessive patch end | Cluster analysis p-value | Excess direction | Source | Target TF | Biosample name |
|---|---|---|---|---|---|---|
| 2250 | 2400 | $3.3 \cdot 10^{-2}$ | increase | ENCFF966KGI | CTCF | A549 |
| -1900 | 1800 | $1.0 \cdot 10^{-3}$ | increase | ENCFF966KGI | CTCF | A549 |
| 1875 | 2200 | $2.0 \cdot 10^{-3}$ | increase | ENCFF966KGI | CTCF | A549 |
| -1775 | 1450 | $1.0 \cdot 10^{-3}$ | increase | ENCFF017XLW | CTCF | GM12878 |
| 1850 | 2050 | $2.9 \cdot 10^{-2}$ | increase | ENCFF017XLW | CTCF | GM12878 |
| -3425 | -3275 | $4.9 \cdot 10^{-2}$ | increase | ENCFF536RGD | CTCF | GM12878 |
| -2725 | -2450 | $4.0 \cdot 10^{-3}$ | increase | ENCFF536RGD | CTCF | GM12878 |
| -2300 | -1975 | $1.0 \cdot 10^{-3}$ | increase | ENCFF536RGD | CTCF | GM12878 |
| -1900 | 1750 | $1.0 \cdot 10^{-3}$ | increase | ENCFF536RGD | CTCF | GM12878 |
| 2150 | 2550 | $1.0 \cdot 10^{-3}$ | increase | ENCFF536RGD | CTCF | GM12878 |
| 2650 | 2900 | $3.0 \cdot 10^{-3}$ | increase | ENCFF536RGD | CTCF | GM12878 |
| -1775 | 2050 | $1.0 \cdot 10^{-3}$ | increase | ENCFF692RPA | CTCF | H1 |
| -3975 | -3775 | $2.1 \cdot 10^{-2}$ | increase | ENCFF692RPA | CTCF | H1 |
| 2075 | 2375 | $7.0 \cdot 10^{-3}$ | increase | ENCFF123DAC | CTCF | HepG2 |
| -2400 | -2150 | $1.0 \cdot 10^{-2}$ | increase | ENCFF123DAC | CTCF | HepG2 |
| -1650 | 1800 | $1.0 \cdot 10^{-3}$ | increase | ENCFF123DAC | CTCF | HepG2 |
| -2050 | -1700 | $2.0 \cdot 10^{-3}$ | increase | ENCFF123DAC | CTCF | HepG2 |
| 3500 | 3700 | $2.2 \cdot 10^{-2}$ | increase | ENCFF252PLM | CTCF | HepG2 |
| -1625 | 2400 | $1.0 \cdot 10^{-3}$ | increase | ENCFF252PLM | CTCF | HepG2 |
| -2600 | 2800 | $1.0 \cdot 10^{-3}$ | increase | ENCFF769AUF | CTCF | K562 |
| -2800 | -2650 | $5.0 \cdot 10^{-2}$ | increase | ENCFF769AUF | CTCF | K562 |
| -300 | 250 | $1.0 \cdot 10^{-3}$ | increase | GSM3560258 | CTCF | K562 |
| -2525 | 2325 | $1.0 \cdot 10^{-3}$ | increase | ENCFF820BVW | CTCF | MCF-7 |
| -2850 | 2425 | $1.0 \cdot 10^{-3}$ | increase | ENCFF962IZR | CTCF | MCF-7 |
| -2000 | -1575 | $1.0 \cdot 10^{-3}$ | increase | ENCFF503DSN | CTCF | WTC11 |
| -2725 | -2450 | $6.0 \cdot 10^{-3}$ | increase | ENCFF503DSN | CTCF | WTC11 |
| -3900 | -3625 | $6.0 \cdot 10^{-3}$ | increase | ENCFF503DSN | CTCF | WTC11 |
| -2400 | -2075 | $2.0 \cdot 10^{-3}$ | increase | ENCFF503DSN | CTCF | WTC11 |
| -1475 | 1800 | $1.0 \cdot 10^{-3}$ | increase | ENCFF503DSN | CTCF | WTC11 |
| 2250 | 2425 | $3.4 \cdot 10^{-2}$ | increase | ENCFF503DSN | CTCF | WTC11 |
| -2000 | 525 | $1.0 \cdot 10^{-3}$ | increase | ENCFF578XIP | FOXK2 | GM12878 |
| -1425 | 1525 | $1.0 \cdot 10^{-3}$ | increase | ENCFF855EOH | FOXK2 | GM12878 |
| -1450 | 1900 | $1.0 \cdot 10^{-3}$ | increase | ENCFF008HBF | FOXK2 | HepG2 |
| -1400 | 275 | $1.0 \cdot 10^{-3}$ | increase | ENCFF997KQI | FOXK2 | HepG2 |
| -1775 | 1725 | $1.0 \cdot 10^{-3}$ | increase | ENCFF022HGL | FOXK2 | K562 |
| -1975 | -1700 | $1.7 \cdot 10^{-2}$ | increase | ENCFF154SRU | FOXK2 | K562 |
| -1650 | 2150 | $1.0 \cdot 10^{-3}$ | increase | ENCFF154SRU | FOXK2 | K562 |
| -1275 | 200 | $1.0 \cdot 10^{-3}$ | increase | ENCFF356XXS | FOXK2 | MCF-7 |
| -1325 | 1425 | $1.0 \cdot 10^{-3}$ | increase | ENCFF583QAD | FOXK2 | MCF-7 |
| -1650 | 1575 | $1.0 \cdot 10^{-3}$ | increase | ENCFF174QAR | IRF1 | HepG2 |
| -2625 | -2450 | $1.4 \cdot 10^{-2}$ | increase | ENCFF019LOH | IRF1 | K562 |
| -2300 | -2150 | $3.1 \cdot 10^{-2}$ | increase | ENCFF019LOH | IRF1 | K562 |
| -2100 | -1675 | $1.0 \cdot 10^{-3}$ | increase | ENCFF019LOH | IRF1 | K562 |
| -1625 | 1725 | $1.0 \cdot 10^{-3}$ | increase | ENCFF019LOH | IRF1 | K562 |
| 1800 | 2550 | $1.0 \cdot 10^{-3}$ | increase | ENCFF019LOH | IRF1 | K562 |
| 1775 | 2000 | $1.6 \cdot 10^{-2}$ | increase | ENCFF251ZSJ | IRF1 | K562 |
| -2575 | -2225 | $3.0 \cdot 10^{-3}$ | increase | ENCFF251ZSJ | IRF1 | K562 |
| 2250 | 2550 | $6.0 \cdot 10^{-3}$ | increase | ENCFF251ZSJ | IRF1 | K562 |
| -1175 | 1650 | $1.0 \cdot 10^{-3}$ | increase | ENCFF251ZSJ | IRF1 | K562 |
| -1850 | 1950 | $1.0 \cdot 10^{-3}$ | increase | ENCFF910ZUK | IRF1 | WTC11 |
| 25 | 475 | $1.0 \cdot 10^{-3}$ | increase | ENCFF131JIX | MEF2A | GM12878 |

| | | | | | | |
|---|---|---|---|---|---|---|
| -650 | -25 | $1.0 \cdot 10^{-3}$ | increase | ENCFF131JIX | MEF2A | GM12878 |
| -1100 | -925 | $1.7 \cdot 10^{-2}$ | increase | ENCFF775DWU | MEF2A | GM12878 |
| -1625 | -1475 | $3.2 \cdot 10^{-2}$ | increase | ENCFF775DWU | MEF2A | GM12878 |
| -825 | 1000 | $1.0 \cdot 10^{-3}$ | increase | ENCFF775DWU | MEF2A | GM12878 |
| -1375 | 1350 | $1.0 \cdot 10^{-3}$ | increase | ENCFF987BRO | MEF2A | HepG2 |
| -900 | -250 | $1.0 \cdot 10^{-3}$ | increase | ENCFF190IHG | MEF2A | K562 |
| 425 | 575 | $4.0 \cdot 10^{-2}$ | increase | ENCFF190IHG | MEF2A | K562 |
| 25 | 225 | $6.0 \cdot 10^{-3}$ | increase | ENCFF190IHG | MEF2A | K562 |
| -175 | -25 | $9.0 \cdot 10^{-3}$ | increase | ENCFF190IHG | MEF2A | K562 |
| -1300 | 700 | $1.0 \cdot 10^{-3}$ | increase | ENCFF288TSN | MEF2A | K562 |
| 1325 | 1500 | $1.3 \cdot 10^{-2}$ | increase | ENCFF288TSN | MEF2A | K562 |
| 750 | 1100 | $4.0 \cdot 10^{-3}$ | increase | ENCFF288TSN | MEF2A | K562 |
| -2175 | -1875 | $2.3 \cdot 10^{-2}$ | increase | ENCFF356PWE | MYC | A549 |
| -1825 | 1850 | $1.0 \cdot 10^{-3}$ | increase | ENCFF356PWE | MYC | A549 |
| -2200 | 1800 | $1.0 \cdot 10^{-3}$ | increase | ENCFF674RQX | MYC | A549 |
| -1475 | 1500 | $1.0 \cdot 10^{-3}$ | increase | ENCFF765CKK | MYC | GM12878 |
| -1575 | 375 | $1.0 \cdot 10^{-3}$ | increase | ENCFF372RQZ | MYC | H1 |
| -1625 | 1825 | $1.0 \cdot 10^{-3}$ | increase | ENCFF700CXD | MYC | H1 |
| -1900 | -1700 | $3.8 \cdot 10^{-2}$ | increase | ENCFF700CXD | MYC | H1 |
| -1675 | 1700 | $1.0 \cdot 10^{-3}$ | increase | ENCFF800JFG | MYC | HepG2 |
| 2000 | 2275 | $1.7 \cdot 10^{-2}$ | increase | ENCFF800JFG | MYC | HepG2 |
| -1750 | 1675 | $1.0 \cdot 10^{-3}$ | increase | ENCFF043WTJ | MYC | K562 |
| -1700 | 1825 | $1.0 \cdot 10^{-3}$ | increase | ENCFF149BIS | MYC | MCF-7 |
| 2000 | 2225 | $1.8 \cdot 10^{-2}$ | increase | ENCFF149BIS | MYC | MCF-7 |
| 175 | 700 | $1.0 \cdot 10^{-3}$ | decrease | ENCFF463AFM | NANOG | GM23338 |
| -600 | -275 | $2.0 \cdot 10^{-3}$ | decrease | ENCFF463AFM | NANOG | GM23338 |
| -125 | 125 | $1.0 \cdot 10^{-3}$ | increase | ENCFF463AFM | NANOG | GM23338 |
| 175 | 700 | $1.0 \cdot 10^{-3}$ | decrease | ENCFF897LBK | NANOG | GM23338 |
| -125 | 125 | $1.0 \cdot 10^{-3}$ | increase | ENCFF897LBK | NANOG | GM23338 |
| -475 | -275 | $6.0 \cdot 10^{-3}$ | decrease | ENCFF897LBK | NANOG | GM23338 |
| 650 | 1450 | $1.0 \cdot 10^{-3}$ | increase | ENCFF655XZF | NANOG | H1 |
| -475 | 500 | $1.0 \cdot 10^{-3}$ | increase | ENCFF655XZF | NANOG | H1 |
| -675 | -525 | $2.3 \cdot 10^{-2}$ | increase | ENCFF655XZF | NANOG | H1 |
| -900 | -725 | $2.3 \cdot 10^{-2}$ | increase | ENCFF655XZF | NANOG | H1 |
| -475 | 150 | $1.0 \cdot 10^{-3}$ | increase | ENCFF722JFZ | NANOG | H1 |
| -675 | -525 | $3.0 \cdot 10^{-2}$ | increase | ENCFF722JFZ | NANOG | H1 |
| -1025 | -750 | $5.0 \cdot 10^{-3}$ | increase | ENCFF722JFZ | NANOG | H1 |
| -1575 | -1300 | $9.0 \cdot 10^{-3}$ | increase | ENCFF722JFZ | NANOG | H1 |
| -1225 | 1375 | $1.0 \cdot 10^{-3}$ | increase | ENCFF252JYE | NFRKB | HepG2 |
| -1200 | 500 | $1.0 \cdot 10^{-3}$ | increase | ENCFF748JYG | NFRKB | HepG2 |
| -3125 | 1775 | $1.0 \cdot 10^{-3}$ | increase | ENCFF048HYG | NFRKB | K562 |
| 2175 | 2900 | $1.0 \cdot 10^{-3}$ | increase | ENCFF048HYG | NFRKB | K562 |
| 1850 | 2125 | $1.0 \cdot 10^{-2}$ | increase | ENCFF048HYG | NFRKB | K562 |
| -2900 | -2100 | $1.0 \cdot 10^{-3}$ | increase | ENCFF594KPG | NFRKB | K562 |
| -2050 | 2850 | $1.0 \cdot 10^{-3}$ | increase | ENCFF594KPG | NFRKB | K562 |
| -300 | 325 | $1.0 \cdot 10^{-3}$ | decrease | ENCFF360OXK | NFRKB | MCF-7 |
| -325 | 325 | $1.0 \cdot 10^{-3}$ | decrease | ENCFF768VQG | NFRKB | MCF-7 |
| 1650 | 1875 | $3.3 \cdot 10^{-2}$ | increase | ENCFF015ADG | RUNX1 | K562 |
| -1325 | 1400 | $1.0 \cdot 10^{-3}$ | increase | ENCFF015ADG | RUNX1 | K562 |
| -1325 | 1225 | $1.0 \cdot 10^{-3}$ | increase | ENCFF264QLP | RUNX1 | K562 |
| -1650 | -1475 | $3.3 \cdot 10^{-2}$ | increase | ENCFF198SPL | SPI1 | GM12878 |
| -1425 | 1425 | $1.0 \cdot 10^{-3}$ | increase | ENCFF198SPL | SPI1 | GM12878 |
| -1150 | 1375 | $1.0 \cdot 10^{-3}$ | increase | ENCFF881ARS | SPI1 | GM12878 |
| -2325 | -2150 | $4.2 \cdot 10^{-2}$ | increase | ENCFF185DGL | SPI1 | K562 |
| -1925 | 1575 | $1.0 \cdot 10^{-3}$ | increase | ENCFF185DGL | SPI1 | K562 |

| | | | | | | |
|---|---|---|---|---|---|---|
| 1625 | 1775 | $2.6 \cdot 10^{-2}$ | increase | ENCFF185DGL | SPI1 | K562 |
| -1575 | 1825 | $1.0 \cdot 10^{-3}$ | increase | ENCFF456BUE | SPI1 | K562 |
| -1925 | -1650 | $1.0 \cdot 10^{-2}$ | increase | ENCFF456BUE | SPI1 | K562 |
| -1725 | 1600 | $1.0 \cdot 10^{-3}$ | increase | ENCFF488WQN | TP53 | A549 |
| -1650 | 1725 | $1.0 \cdot 10^{-3}$ | increase | ENCFF701DTE | TP53 | HepG2 |
| 2450 | 2750 | $3.9 \cdot 10^{-2}$ | increase | ENCFF280RDQ | TP53 | WTC11 |
| -1900 | 2175 | $1.0 \cdot 10^{-3}$ | increase | ENCFF280RDQ | TP53 | WTC11 |

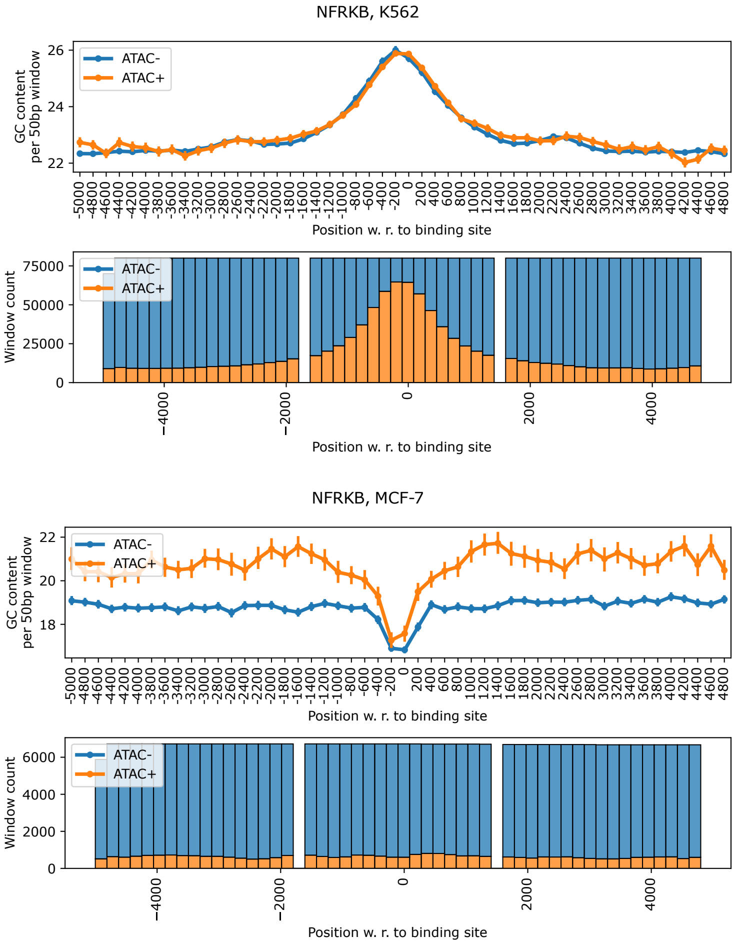


**Fig. S11.** Overview of ATAC-seq and ChIP-seq combined information on NFRKB in K562 (top pair of plots) and in MCF-7 (bottom pair of plots). In the top plot, the average GC content (with 95% confidence interval shown as error bars) by position with respect to the binding site, distinguished for k-mers overlapping with an ATAC-seq peak (ATAC+) and k-mers not overlapping with an ATAC-seq peak (ATAC-). In the bottom plot, the distribution of ATAC+ aand ATAC- k-mers per position.

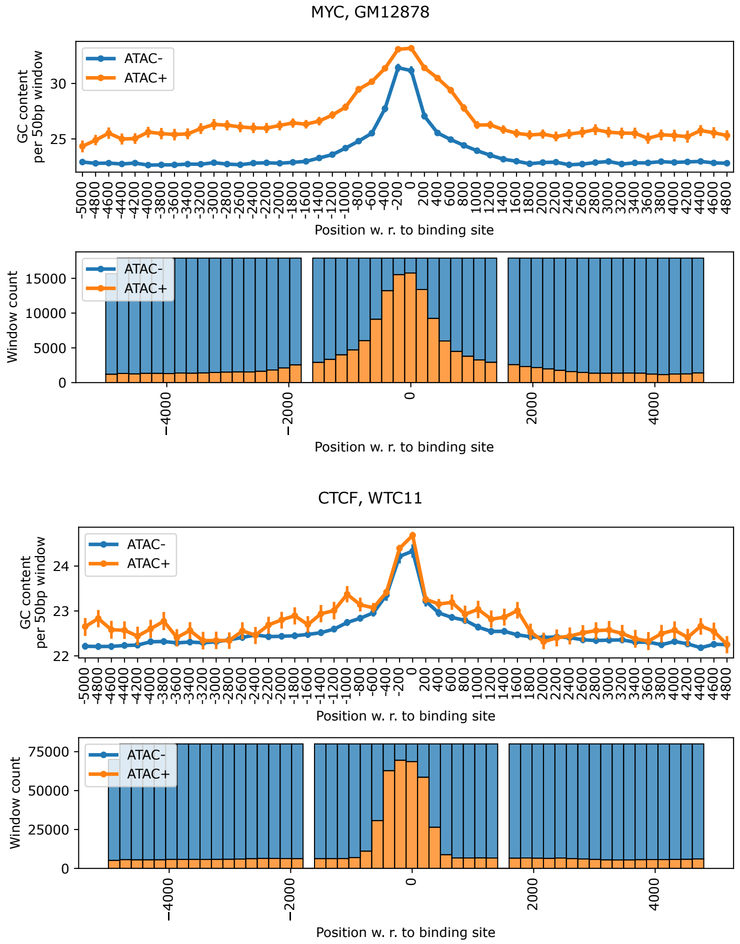


**Fig. S12.** Overview of ATAC-seq and ChIP-seq combined information on MYC in GM12878 (top pair of plots) and on CTCF in WTC11 (bottom pair of plots). In the top plot, the average GC content (with 95% confidence interval shown as error bars) by position with respect to the binding site, distinguished for k-mers overlapping with an ATAC-seq peak (ATAC+) and k-mers not overlapping with an ATAC-seq peak (ATAC-). In the bottom plot, the distribution of ATAC+ aand ATAC- k-mers per position.

**Table S4.** Average GC content for k-mers overlapping with an ATAC-seq peak (ATAC+) and k-mers not overlapping with an ATAC-seq peak (ATAC-) per chosen positions (w. r. to the binding site). In every cell, first value is the average GC content in ATAC+ windows, second one described the average GC content in ATAC- windows. Statistical significance (Mann-Whitney U test with Holm correction for multiple testing for ATAC+ and ATAC- windows, "greater" alternative hypothesis) encoded as stars: $^{****}p < 10^{-4}$, $^{***}p < 10^{-3}$, $^{**}p < 10^{-2}$, $^{*}p < 0.05$; ns omitted.

| ENCODE file ID | Target TF | Cell line | -3000 | -1000 | 0 | 1000 | 3000 |
|---|---|---|---|---|---|---|---|
| ENCFF966KGI | CTCF | A549 | 25.9 (s. d. 6.5) / 22.0 (s. d. 6.2)**** | 26.3 (s. d. 7.1) / 22.2 (s. d. 6.3)**** | 25.0 (s. d. 6.9) / 21.6 (s. d. 6.9)**** | 26.2 (s. d. 6.9) / 22.2 (s. d. 6.3)**** | 26.2 (s. d. 6.8) / 22.0 (s. d. 6.3)**** |
| ENCFF017XLW | CTCF | GM12878 | 23.5 (s. d. 6.7) / 22.1 (s. d. 6.3)**** | 23.9 (s. d. 7.1) / 22.6 (s. d. 6.5)**** | 24.6 (s. d. 6.8) / 24.3 (s. d. 6.7)**** | 23.8 (s. d. 6.9) / 22.4 (s. d. 6.5)**** | 23.3 (s. d. 6.9) / 22.1 (s. d. 6.3)**** |
| ENCFF536RGD | CTCF | GM12878 | 24.5 (s. d. 6.8) / 22.0 (s. d. 6.3)**** | 25.7 (s. d. 7.5) / 22.3 (s. d. 6.4)**** | 24.6 (s. d. 6.8) / 23.7 (s. d. 6.7)**** | 25.2 (s. d. 7.1) / 22.3 (s. d. 6.3)**** | 24.9 (s. d. 7.2) / 22.0 (s. d. 6.3)**** |
| ENCFF123DAC | CTCF | HepG2 | 23.8 (s. d. 6.8) / 22.2 (s. d. 6.4)**** | 24.4 (s. d. 7.0) / 22.6 (s. d. 6.5)**** | 25.2 (s. d. 6.7) / 24.1 (s. d. 6.8)**** | 24.1 (s. d. 6.8) / 22.6 (s. d. 6.5)**** | 23.8 (s. d. 6.8) / 22.3 (s. d. 6.4)**** |
| ENCFF252PLM | CTCF | HepG2 | 23.1 (s. d. 6.6) / 22.3 (s. d. 6.4)**** | 23.7 (s. d. 6.6) / 22.8 (s. d. 6.6)**** | 24.9 (s. d. 6.7) / 24.4 (s. d. 6.7)**** | 23.7 (s. d. 6.9) / 22.6 (s. d. 6.5)**** | 23.2 (s. d. 6.7) / 22.3 (s. d. 6.4)**** |
| ENCFF769AUF | CTCF | K562 | 22.8 (s. d. 6.4) / 22.8 (s. d. 6.4) | 23.6 (s. d. 6.5) / 23.3 (s. d. 6.6)*** | 25.2 (s. d. 6.5) / 24.6 (s. d. 6.5)**** | 23.3 (s. d. 6.4) / 23.2 (s. d. 6.5) | 23.1 (s. d. 6.6) / 22.8 (s. d. 6.4)** |
| ENCFF820BVW | CTCF | MCF-7 | 23.6 (s. d. 6.7) / 23.3 (s. d. 6.4)* | 24.8 (s. d. 7.0) / 24.2 (s. d. 6.7)**** | 27.7 (s. d. 6.4) / 27.6 (s. d. 6.4) | 24.8 (s. d. 6.6) / 24.0 (s. d. 6.6)**** | 24.0 (s. d. 6.6) / 23.2 (s. d. 6.4)**** |
| ENCFF962IZR | CTCF | MCF-7 | 23.3 (s. d. 6.6) / 23.4 (s. d. 6.4) | 24.2 (s. d. 6.8) / 24.3 (s. d. 6.7) | 27.7 (s. d. 6.4) / 27.6 (s. d. 6.4) | 24.0 (s. d. 6.5) / 24.1 (s. d. 6.6) | 23.4 (s. d. 6.5) / 23.3 (s. d. 6.4) |
| ENCFF503DSN | CTCF | WTC11 | 22.3 (s. d. 6.3) / 22.3 (s. d. 6.4) | 23.4 (s. d. 6.7) / 22.7 (s. d. 6.6)**** | 24.7 (s. d. 6.9) / 24.3 (s. d. 6.7)*** | 23.0 (s. d. 6.7) / 22.6 (s. d. 6.5)** | 22.6 (s. d. 6.6) / 22.3 (s. d. 6.4) |
| ENCFF578XIP | FOXK2 | GM12878 | 23.1 (s. d. 6.5) / 23.0 (s. d. 6.5) | 24.6 (s. d. 6.8) / 24.2 (s. d. 6.7)**** | 27.4 (s. d. 6.6) / 26.8 (s. d. 6.5)**** | 23.8 (s. d. 6.7) / 23.8 (s. d. 6.7) | 23.0 (s. d. 6.4) / 23.0 (s. d. 6.5) |
| ENCFF855EOH | FOXK2 | GM12878 | 23.3 (s. d. 6.2) / 23.1 (s. d. 6.3) | 24.4 (s. d. 6.4) / 24.3 (s. d. 6.7) | 27.9 (s. d. 6.4) / 27.0 (s. d. 6.2)**** | 24.1 (s. d. 6.9) / 23.9 (s. d. 6.5) | 23.2 (s. d. 6.3) / 23.1 (s. d. 6.3) |
| ENCFF008HBF | FOXK2 | HepG2 | 22.6 (s. d. 6.4) / 22.7 (s. d. 6.3) | 23.6 (s. d. 6.6) / 23.4 (s. d. 6.5) | 25.2 (s. d. 6.2) / 24.9 (s. d. 6.1)*** | 23.4 (s. d. 6.4) / 23.2 (s. d. 6.4)* | 22.6 (s. d. 6.3) / 22.7 (s. d. 6.3) |
| ENCFF997KQI | FOXK2 | HepG2 | 22.5 (s. d. 6.2) / 22.5 (s. d. 6.4) | 23.1 (s. d. 6.6) / 23.2 (s. d. 6.5) | 24.9 (s. d. 6.1) / 23.9 (s. d. 5.7)**** | 23.3 (s. d. 6.4) / 22.9 (s. d. 6.4)**** | 22.5 (s. d. 6.6) / 22.4 (s. d. 6.4) |

| | | | | | | | |
|---|---|---|---|---|---|---|---|
| ENCFF022HGL | FOXK2 | K562 | 22.5 (s. d. 6.4) / 22.3 (s. d. 6.4)* | 23.4 (s. d. 6.8) / 23.2 (s. d. 6.8) | 25.2 (s. d. 6.4) / 25.0 (s. d. 6.5) | 23.0 (s. d. 6.7) / 23.0 (s. d. 6.6) | 22.6 (s. d. 6.4) / 22.4 (s. d. 6.3) |
| ENCFF154SRU | FOXK2 | K562 | 22.3 (s. d. 6.3) / 22.3 (s. d. 6.4) | 23.4 (s. d. 6.8) / 23.3 (s. d. 6.7) | 25.6 (s. d. 6.4) / 25.5 (s. d. 6.4) | 23.1 (s. d. 6.8) / 23.0 (s. d. 6.6) | 22.6 (s. d. 6.4) / 22.4 (s. d. 6.3)** |
| ENCFF356XXS | FOXK2 | MCF-7 | 25.5 (s. d. 6.3) / 22.7 (s. d. 6.6)**** | 26.3 (s. d. 6.3) / 23.5 (s. d. 6.6)**** | 29.2 (s. d. 6.6) / 23.6 (s. d. 5.2)**** | 26.1 (s. d. 6.5) / 23.5 (s. d. 6.4)**** | 26.5 (s. d. 6.5) / 22.7 (s. d. 6.6)**** |
| ENCFF583QAD | FOXK2 | MCF-7 | 24.7 (s. d. 6.8) / 23.4 (s. d. 6.3)*** | 23.8 (s. d. 6.3) / 24.4 (s. d. 6.4) | 28.0 (s. d. 6.7) / 24.8 (s. d. 5.6)**** | 23.8 (s. d. 6.0) / 24.3 (s. d. 6.4) | 23.1 (s. d. 6.1) / 23.2 (s. d. 6.2) |
| ENCFF174QAR | IRF1 | HepG2 | 27.5 (s. d. 6.8) / 22.7 (s. d. 6.3)**** | 28.7 (s. d. 7.1) / 23.8 (s. d. 6.6)**** | 31.3 (s. d. 6.0) / 28.0 (s. d. 6.3)**** | 27.8 (s. d. 7.0) / 23.5 (s. d. 6.3)**** | 27.6 (s. d. 6.7) / 22.8 (s. d. 6.2)**** |
| ENCFF019LOH | IRF1 | K562 | 21.9 (s. d. 6.3) / 21.9 (s. d. 6.2) | 22.4 (s. d. 6.6) / 22.3 (s. d. 6.5) | 23.0 (s. d. 6.2) / 23.2 (s. d. 6.1) | 22.2 (s. d. 6.3) / 22.3 (s. d. 6.4) | 21.9 (s. d. 6.3) / 22.0 (s. d. 6.2) |
| ENCFF251ZSJ | IRF1 | K562 | 21.7 (s. d. 6.2) / 21.9 (s. d. 6.2) | 22.4 (s. d. 6.6) / 22.4 (s. d. 6.5) | 23.1 (s. d. 6.2) / 23.1 (s. d. 6.2) | 22.3 (s. d. 6.4) / 22.2 (s. d. 6.4) | 22.0 (s. d. 6.3) / 22.0 (s. d. 6.2) |
| ENCFF910ZUK | IRF1 | WTC11 | 28.2 (s. d. 6.5) / 22.8 (s. d. 6.3)**** | 30.9 (s. d. 6.9) / 24.4 (s. d. 6.7)**** | 33.4 (s. d. 5.3) / 31.6 (s. d. 5.8)**** | 29.9 (s. d. 7.1) / 24.1 (s. d. 6.5)**** | 28.0 (s. d. 6.9) / 22.8 (s. d. 6.3)**** |
| ENCFF131JIX | MEF2A | GM12878 | 23.2 (s. d. 6.7) / 21.5 (s. d. 6.1)**** | 23.3 (s. d. 6.6) / 21.7 (s. d. 6.2)**** | 23.0 (s. d. 6.8) / 22.4 (s. d. 7.0)**** | 23.0 (s. d. 6.7) / 21.7 (s. d. 6.2)**** | 22.9 (s. d. 6.4) / 21.5 (s. d. 6.2)**** |
| ENCFF775DWU | MEF2A | GM12878 | 23.3 (s. d. 6.6) / 21.3 (s. d. 6.0)**** | 23.2 (s. d. 6.6) / 21.6 (s. d. 6.2)**** | 23.0 (s. d. 6.7) / 21.6 (s. d. 6.7)**** | 23.0 (s. d. 6.7) / 21.6 (s. d. 6.1)**** | 23.1 (s. d. 6.5) / 21.3 (s. d. 6.0)**** |
| ENCFF987BRO | MEF2A | HepG2 | 22.4 (s. d. 6.5) / 22.1 (s. d. 6.3) | 22.9 (s. d. 6.6) / 22.9 (s. d. 6.7) | 24.8 (s. d. 7.1) / 24.5 (s. d. 7.3) | 22.6 (s. d. 6.4) / 22.6 (s. d. 6.5) | 22.3 (s. d. 6.5) / 22.2 (s. d. 6.2) |
| ENCFF190IHG | MEF2A | K562 | 24.4 (s. d. 6.9) / 22.1 (s. d. 6.2)**** | 25.1 (s. d. 6.8) / 22.1 (s. d. 6.4)**** | 23.5 (s. d. 6.2) / 21.2 (s. d. 6.1)**** | 24.3 (s. d. 6.7) / 22.1 (s. d. 6.2)**** | 24.6 (s. d. 6.3) / 21.9 (s. d. 6.2)**** |
| ENCFF288TSN | MEF2A | K562 | 24.6 (s. d. 6.7) / 21.8 (s. d. 6.2)**** | 25.1 (s. d. 6.9) / 21.9 (s. d. 6.3)**** | 24.3 (s. d. 6.4) / 20.7 (s. d. 5.7)**** | 24.8 (s. d. 6.9) / 21.8 (s. d. 6.3)**** | 24.7 (s. d. 6.6) / 21.7 (s. d. 6.2)**** |
| ENCFF356PWE | MYC | A549 | 23.2 (s. d. 6.3) / 23.1 (s. d. 6.3) | 25.1 (s. d. 7.0) / 25.1 (s. d. 7.0) | 30.6 (s. d. 6.8) / 30.7 (s. d. 6.9) | 24.5 (s. d. 6.8) / 24.5 (s. d. 6.9) | 23.2 (s. d. 6.4) / 23.1 (s. d. 6.4) |

| | | | | | | | |
|---|---|---|---|---|---|---|---|
| ENCFF674RQX | MYC | A549 | 25.0 (s. d. 6.7) / 23.0 (s. d. 6.2)**** | 26.5 (s. d. 7.5) / 24.5 (s. d. 6.8)**** | 30.8 (s. d. 6.8) / 29.2 (s. d. 7.1)**** | 26.0 (s. d. 7.2) / 24.0 (s. d. 6.7)**** | 25.1 (s. d. 7.0) / 22.9 (s. d. 6.3)**** |
| ENCFF765CKK | MYC | GM12878 | 26.3 (s. d. 7.2) / 22.9 (s. d. 6.2)**** | 27.9 (s. d. 7.4) / 24.2 (s. d. 6.8)**** | 33.2 (s. d. 5.3) / 31.2 (s. d. 6.3)**** | 26.2 (s. d. 7.4) / 23.9 (s. d. 6.6)**** | 25.6 (s. d. 6.9) / 23.0 (s. d. 6.3)**** |
| ENCFF800JFG | MYC | HepG2 | 26.8 (s. d. 6.5) / 23.1 (s. d. 6.3)**** | 28.7 (s. d. 7.1) / 24.0 (s. d. 6.4)**** | 31.2 (s. d. 6.2) / 29.8 (s. d. 6.8)**** | 28.5 (s. d. 6.9) / 23.6 (s. d. 6.3)**** | 27.1 (s. d. 6.5) / 23.0 (s. d. 6.3)**** |
| ENCFF043WTJ | MYC | K562 | 23.6 (s. d. 6.5) / 23.4 (s. d. 6.4) | 25.3 (s. d. 7.1) / 25.1 (s. d. 7.2) | 29.1 (s. d. 6.2) / 28.5 (s. d. 6.3)**** | 24.7 (s. d. 6.9) / 24.7 (s. d. 6.9) | 23.5 (s. d. 6.6) / 23.4 (s. d. 6.4) |
| ENCFF149BIS | MYC | MCF-7 | 26.3 (s. d. 6.4) / 23.0 (s. d. 6.3)**** | 28.9 (s. d. 7.2) / 24.0 (s. d. 6.5)**** | 30.2 (s. d. 5.9) / 27.4 (s. d. 6.3)**** | 28.0 (s. d. 7.1) / 23.8 (s. d. 6.4)**** | 26.5 (s. d. 6.6) / 23.0 (s. d. 6.3)**** |
| ENCFF463AFM | NANOG | GM23338 | 23.9 (s. d. 6.8) / 20.1 (s. d. 5.7)**** | 22.9 (s. d. 6.5) / 19.9 (s. d. 5.6)**** | 21.7 (s. d. 5.2) / 19.6 (s. d. 5.0)**** | 22.7 (s. d. 6.6) / 20.1 (s. d. 5.8)**** | 23.9 (s. d. 6.9) / 20.2 (s. d. 5.7)**** |
| ENCFF897LBK | NANOG | GM23338 | 24.0 (s. d. 7.0) / 20.1 (s. d. 5.7)**** | 23.0 (s. d. 6.6) / 20.0 (s. d. 5.7)**** | 21.7 (s. d. 5.3) / 19.3 (s. d. 4.9)**** | 22.7 (s. d. 6.6) / 20.1 (s. d. 5.8)**** | 23.8 (s. d. 6.9) / 20.2 (s. d. 5.7)**** |
| ENCFF252JYE | NFRKB | HepG2 | 22.8 (s. d. 6.1) / 22.4 (s. d. 6.3) | 23.3 (s. d. 6.4) / 23.6 (s. d. 6.5) | 28.2 (s. d. 6.3) / 26.5 (s. d. 6.5)**** | 23.4 (s. d. 6.1) / 23.1 (s. d. 6.5) | 22.5 (s. d. 6.2) / 22.1 (s. d. 6.4) |
| ENCFF748JYG | NFRKB | HepG2 | 22.3 (s. d. 6.7) / 22.0 (s. d. 6.4) | 23.7 (s. d. 6.7) / 22.9 (s. d. 6.4)**** | 27.7 (s. d. 6.4) / 24.8 (s. d. 6.2)**** | 23.0 (s. d. 6.3) / 22.7 (s. d. 6.6) | 22.4 (s. d. 6.1) / 21.8 (s. d. 6.6)* |
| ENCFF048HYG | NFRKB | K562 | 22.8 (s. d. 6.4) / 22.6 (s. d. 6.4)** | 23.6 (s. d. 7.0) / 23.6 (s. d. 7.0) | 25.6 (s. d. 6.6) / 25.7 (s. d. 6.7) | 23.3 (s. d. 6.7) / 23.1 (s. d. 6.8)**** | 22.7 (s. d. 6.7) / 22.4 (s. d. 6.4)**** |
| ENCFF594KPG | NFRKB | K562 | 22.5 (s. d. 6.5) / 22.6 (s. d. 6.5) | 23.7 (s. d. 7.0) / 23.7 (s. d. 7.0) | 25.9 (s. d. 6.7) / 25.7 (s. d. 6.5) | 23.4 (s. d. 6.7) / 23.3 (s. d. 6.8) | 22.6 (s. d. 6.6) / 22.4 (s. d. 6.4) |
| ENCFF360OXK | NFRKB | MCF-7 | 21.1 (s. d. 4.7) / 18.9 (s. d. 5.4)**** | 20.6 (s. d. 5.6) / 18.9 (s. d. 5.3)**** | 17.5 (s. d. 3.8) / 16.9 (s. d. 3.8)*** | 21.2 (s. d. 5.4) / 18.9 (s. d. 5.3)**** | 20.6 (s. d. 5.2) / 18.9 (s. d. 5.3)**** |
| ENCFF768VQG | NFRKB | MCF-7 | 21.0 (s. d. 4.9) / 18.7 (s. d. 5.7)**** | 20.4 (s. d. 5.3) / 18.9 (s. d. 5.6)**** | 17.6 (s. d. 3.9) / 16.8 (s. d. 4.1)**** | 21.3 (s. d. 5.9) / 18.7 (s. d. 5.5)**** | 21.0 (s. d. 5.5) / 18.8 (s. d. 5.4)**** |
| ENCFF015ADG | RUNX1 | K562 | 25.4 (s. d. 6.3) / 22.8 (s. d. 6.3)**** | 27.1 (s. d. 7.0) / 23.3 (s. d. 6.4)**** | 28.9 (s. d. 7.4) / 26.7 (s. d. 7.2)**** | 26.5 (s. d. 7.0) / 23.1 (s. d. 6.4)**** | 26.1 (s. d. 6.7) / 22.8 (s. d. 6.3)**** |

| | | | | | | | |
|---|---|---|---|---|---|---|---|
| ENCFF264QLP | RUNX1 | K562 | 25.3 (s. d. 6.2) / 22.9 (s. d. 6.3)**** | 27.1 (s. d. 7.0) / 23.3 (s. d. 6.4)**** | 28.8 (s. d. 7.3) / 26.6 (s. d. 7.5)**** | 26.3 (s. d. 6.9) / 23.2 (s. d. 6.4)**** | 26.2 (s. d. 6.6) / 22.8 (s. d. 6.3)**** |
| ENCFF198SPL | SPI1 | GM12878 | 22.1 (s. d. 6.3) / 21.8 (s. d. 6.2)** | 22.8 (s. d. 6.5) / 22.1 (s. d. 6.4)**** | 23.3 (s. d. 6.0) / 23.1 (s. d. 5.9) | 22.7 (s. d. 6.6) / 22.0 (s. d. 6.3)**** | 22.0 (s. d. 6.4) / 21.8 (s. d. 6.1)** |
| ENCFF881ARS | SPI1 | GM12878 | 22.0 (s. d. 6.4) / 21.8 (s. d. 6.2) | 22.3 (s. d. 6.5) / 22.2 (s. d. 6.4) | 23.4 (s. d. 6.1) / 23.3 (s. d. 5.9) | 22.4 (s. d. 6.4) / 22.1 (s. d. 6.3)**** | 22.3 (s. d. 6.4) / 21.7 (s. d. 6.2)**** |
| ENCFF185DGL | SPI1 | K562 | 25.1 (s. d. 6.4) / 22.1 (s. d. 6.2)**** | 26.1 (s. d. 6.9) / 22.3 (s. d. 6.2)**** | 25.6 (s. d. 6.2) / 22.7 (s. d. 5.4)**** | 26.1 (s. d. 6.9) / 22.2 (s. d. 6.2)**** | 25.0 (s. d. 6.4) / 22.2 (s. d. 6.2)**** |
| ENCFF456BUE | SPI1 | K562 | 24.8 (s. d. 6.4) / 22.2 (s. d. 6.2)**** | 26.1 (s. d. 6.9) / 22.4 (s. d. 6.2)**** | 25.6 (s. d. 6.2) / 22.6 (s. d. 5.6)**** | 25.9 (s. d. 6.9) / 22.3 (s. d. 6.2)**** | 24.7 (s. d. 6.3) / 22.2 (s. d. 6.2)**** |
| ENCFF488WQN | TP53 | A549 | 23.6 (s. d. 6.4) / 23.6 (s. d. 6.5) | 25.0 (s. d. 7.0) / 25.0 (s. d. 7.1) | 28.5 (s. d. 6.8) / 28.6 (s. d. 6.9) | 24.6 (s. d. 7.1) / 24.6 (s. d. 7.0) | 23.6 (s. d. 6.5) / 23.4 (s. d. 6.5) |
| ENCFF701DTE | TP53 | HepG2 | 23.0 (s. d. 6.4) / 23.0 (s. d. 6.5) | 24.3 (s. d. 7.0) / 24.4 (s. d. 6.9) | 28.9 (s. d. 6.6) / 28.5 (s. d. 6.8)**** | 23.8 (s. d. 6.7) / 24.0 (s. d. 6.8) | 23.1 (s. d. 6.4) / 23.0 (s. d. 6.4) |
| ENCFF280RDQ | TP53 | WTC11 | 23.0 (s. d. 6.7) / 23.3 (s. d. 6.5) | 25.7 (s. d. 7.1) / 25.5 (s. d. 7.1) | 33.2 (s. d. 5.9) / 33.1 (s. d. 6.0) | 25.2 (s. d. 7.0) / 25.0 (s. d. 6.9) | 23.2 (s. d. 6.3) / 23.3 (s. d. 6.5) |

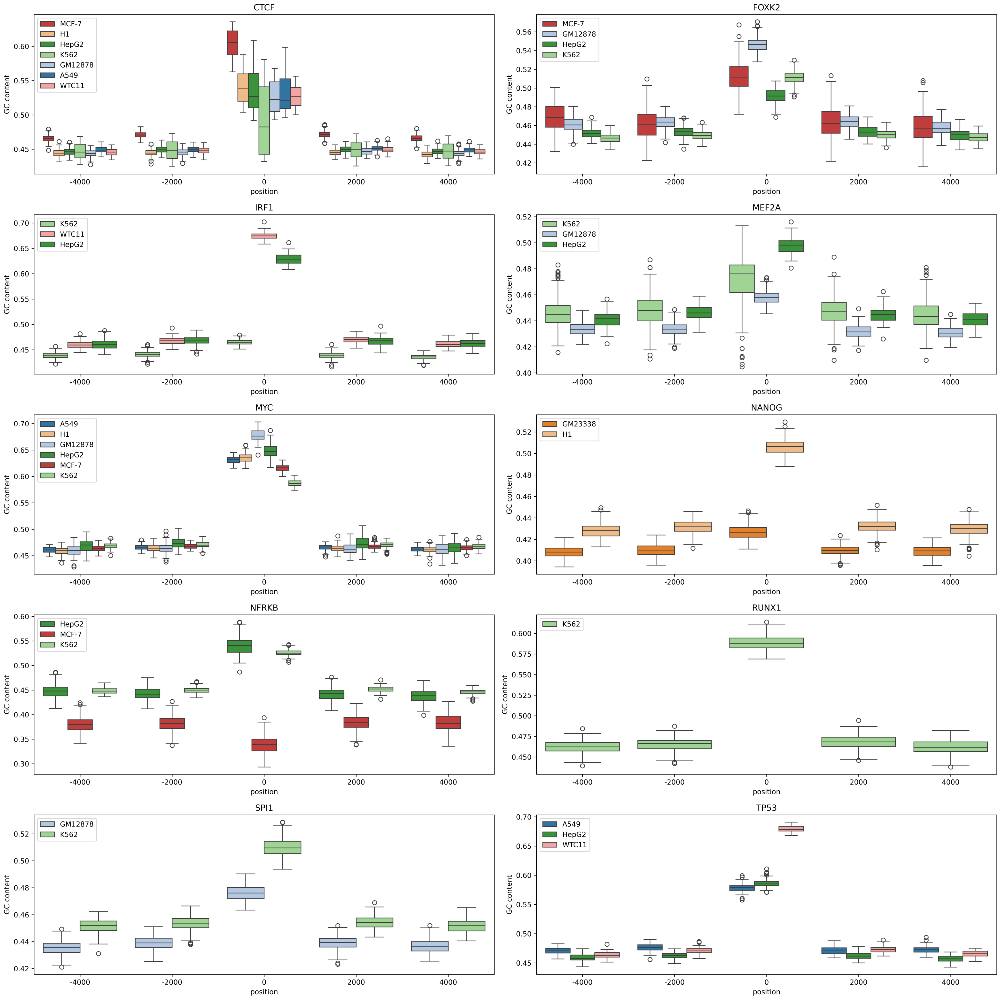


**Fig. S13.** GC content in the vicinity of the position (± 50bp) on the x-axis per TF and cell line. The GC content is calculated per offset from the putative binding site (experimental peak center). Positions were chosen arbitrarily.

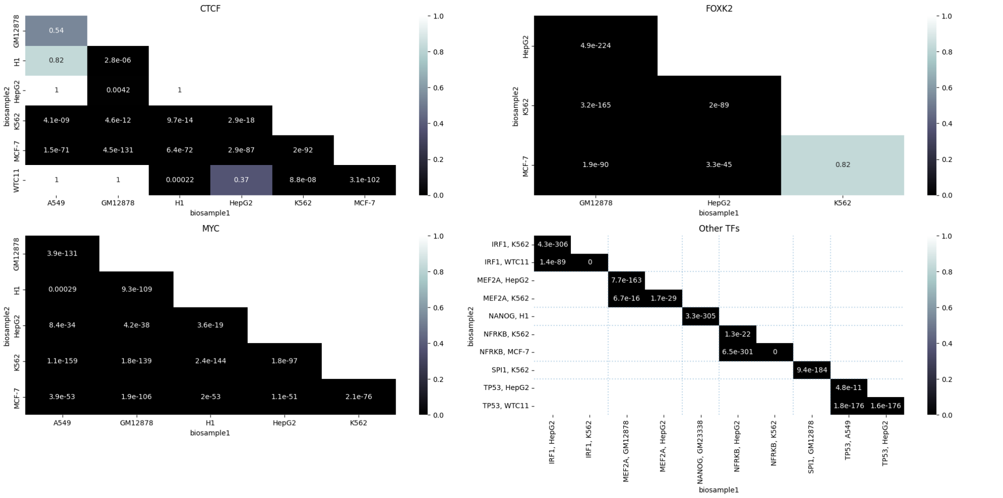


**Fig. S14.** Results of the statistical testing for GC content difference between cell lines in the vicinity of the binding site (±50 bp, as shown in S13 at position 0). The values represent p-values from Mann–Whitney U tests comparing each pair of cell lines, adjusted for multiple testing using the Holm correction.

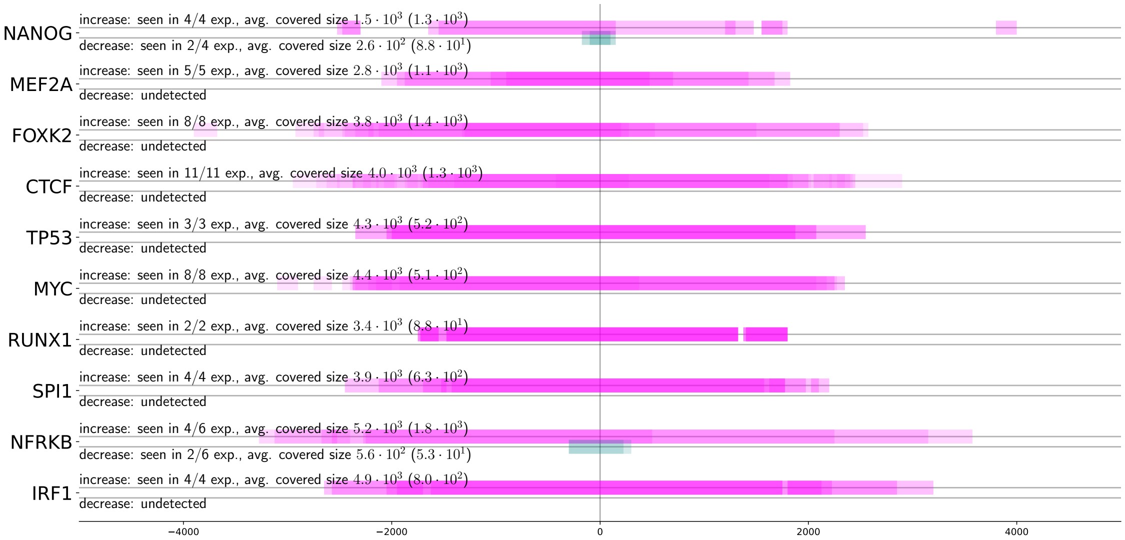


**Fig. S15.** Areas with significant increase or decrease in frequency of CG dinucleotide w. r. to the putative binding site position (middle of the experimental peak) for the tested TFs, using the 50 % most relevant experimental peaks. Decrease is depicted in teal, increase in magenta. Color intensity is proportional to the number of experiments corresponding to the TF – if the color if fully opaque, the excess (increase or decrease) was seen in all experiments for the given TF. The average size of covered area is calculcated as the mean area covered by excessive patches in the given excess direction. If there are no excessive patches found for a direction and experiment, it is not included in the average.

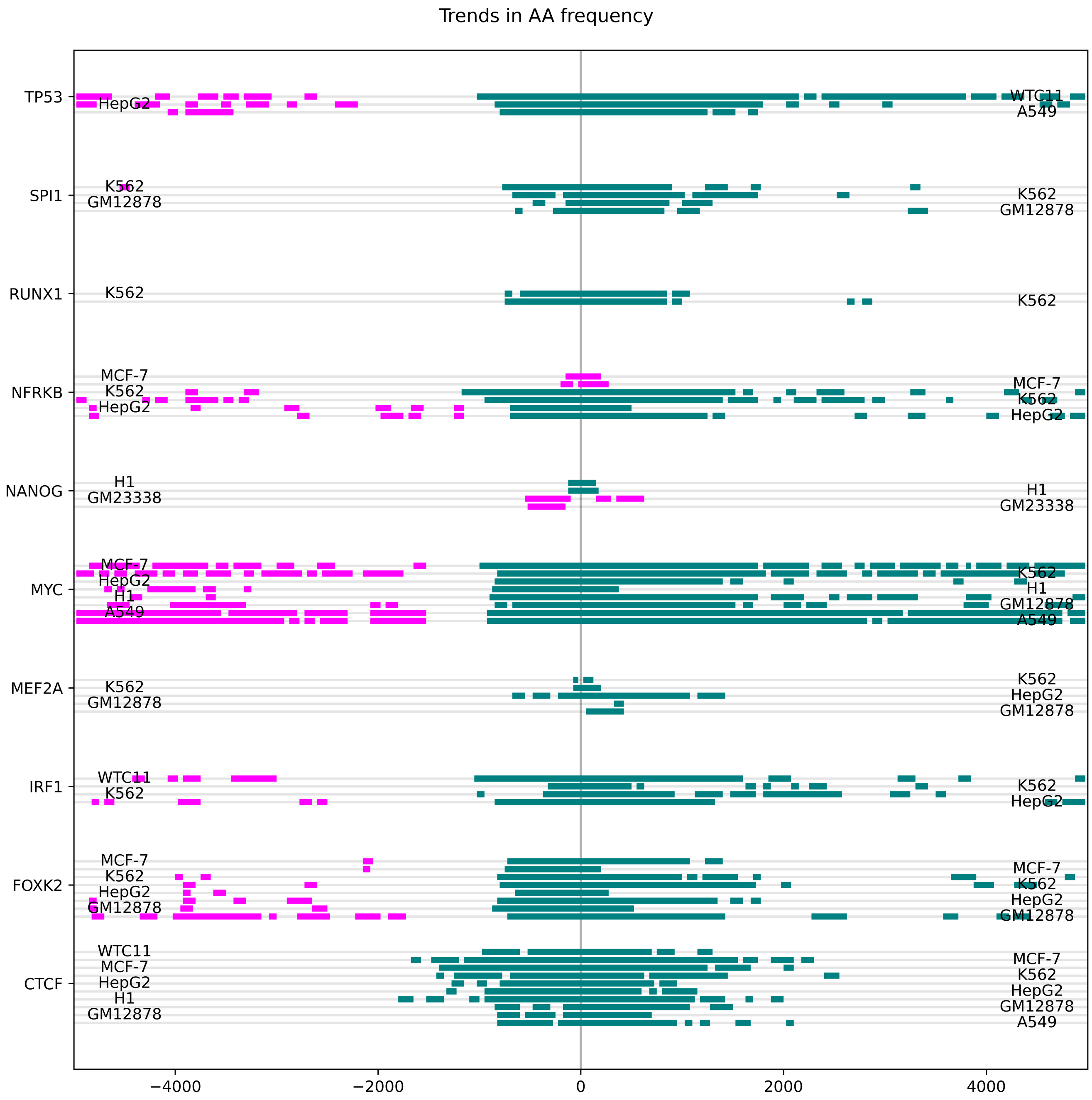


**Fig. S16.** Areas with significant increase or decrease (excessive patches) in frequency of the AA dinucleotide w. r. to the putative binding site position (middle of the experimental peak) for the tested TFs, using the 50 % most relevant experimental peaks. Decrease is depicted in teal, increase in magenta. The excessive patches are estimated using the permutation test.

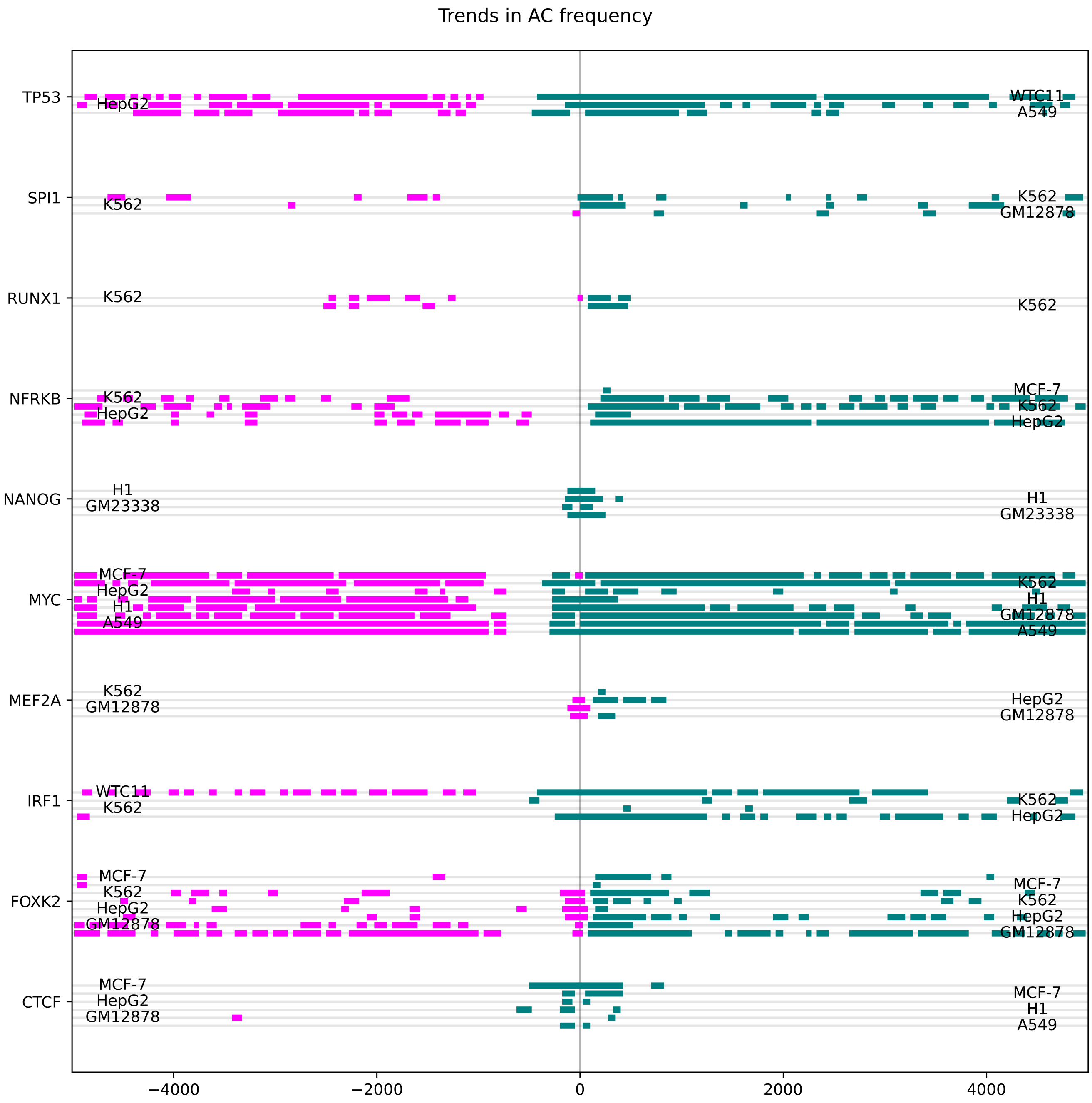


**Fig. S17.** Areas with significant increase or decrease (excessive patches) in frequency of the AC dinucleotide w. r. to the putative binding site position (middle of the experimental peak) for the tested TFs, using the 50 % most relevant experimental peaks. Decrease is depicted in teal, increase in magenta. The excessive patches are estimated using the permutation test.

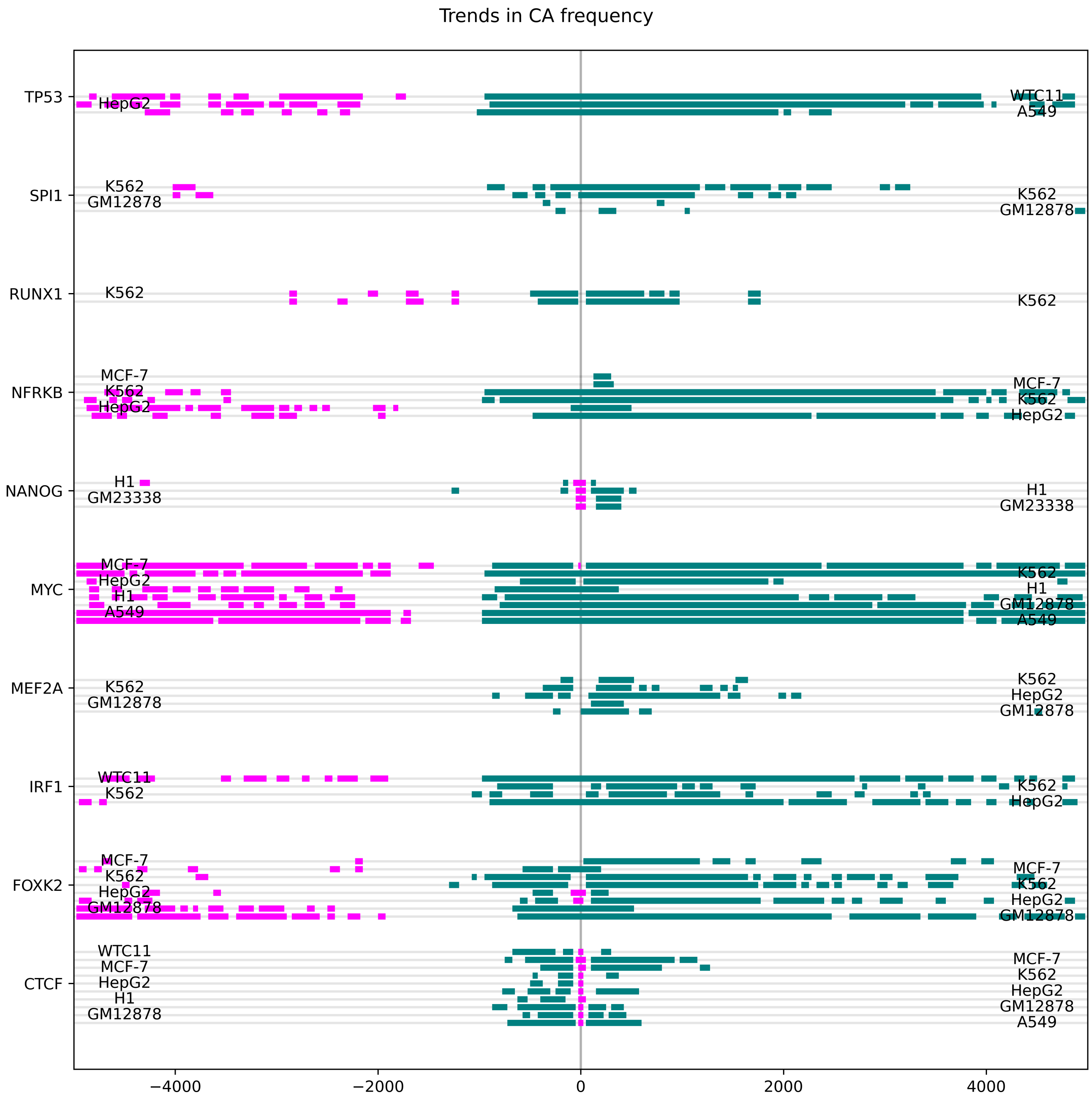


**Fig. S18.** Areas with significant increase or decrease (excessive patches) in frequency of the CA dinucleotide w. r. to the putative binding site position (middle of the experimental peak) for the tested TFs, using the 50 % most relevant experimental peaks. Decrease is depicted in teal, increase in magenta. The excessive patches are estimated using the permutation test.

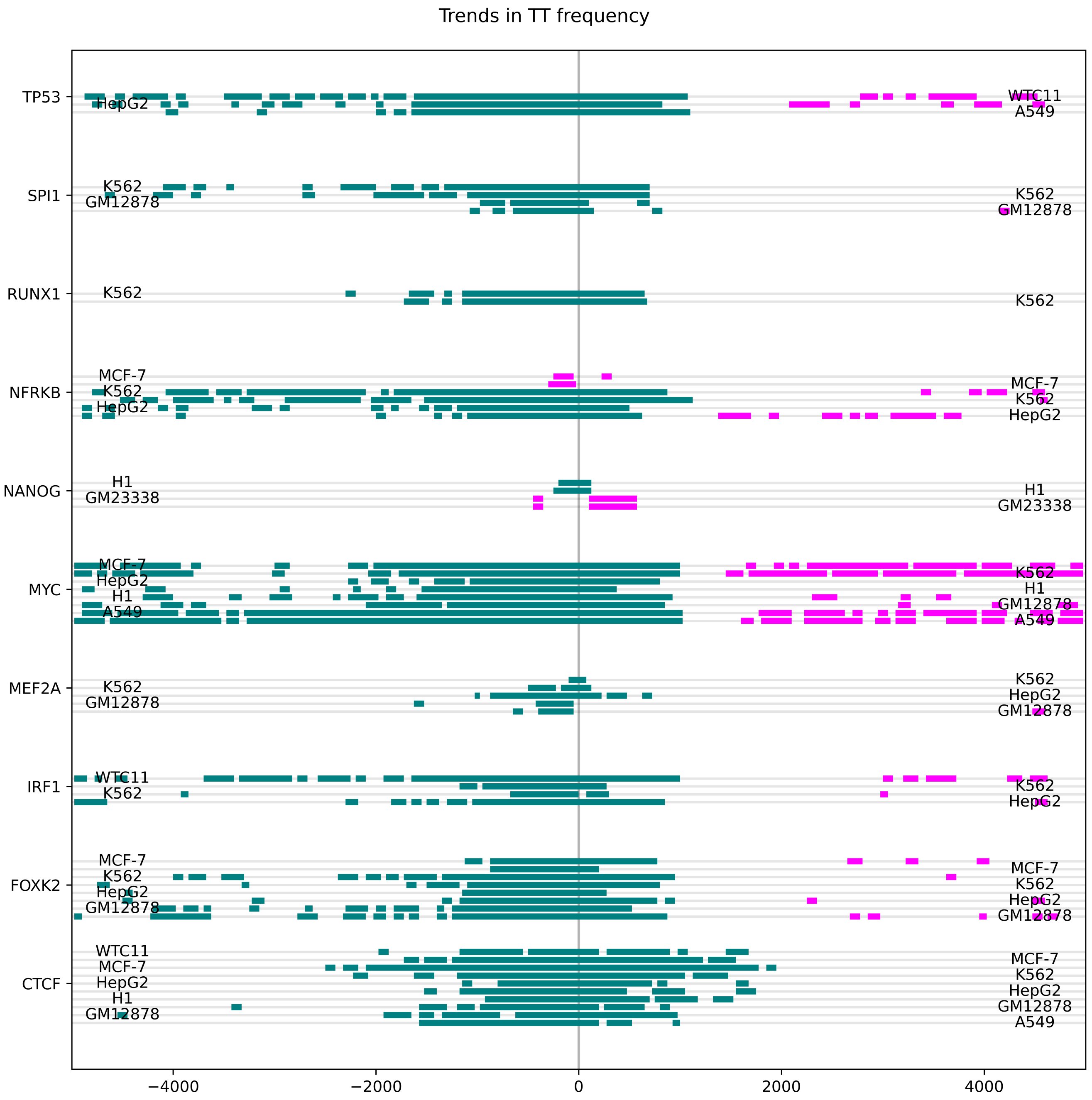


**Fig. S19.** Areas with significant increase or decrease (excessive patches) in frequency of the TT dinucleotide w. r. to the putative binding site position (middle of the experimental peak) for the tested TFs, using the 50 % most relevant experimental peaks. Decrease is depicted in teal, increase in magenta. The excessive patches are estimated using the permutation test.

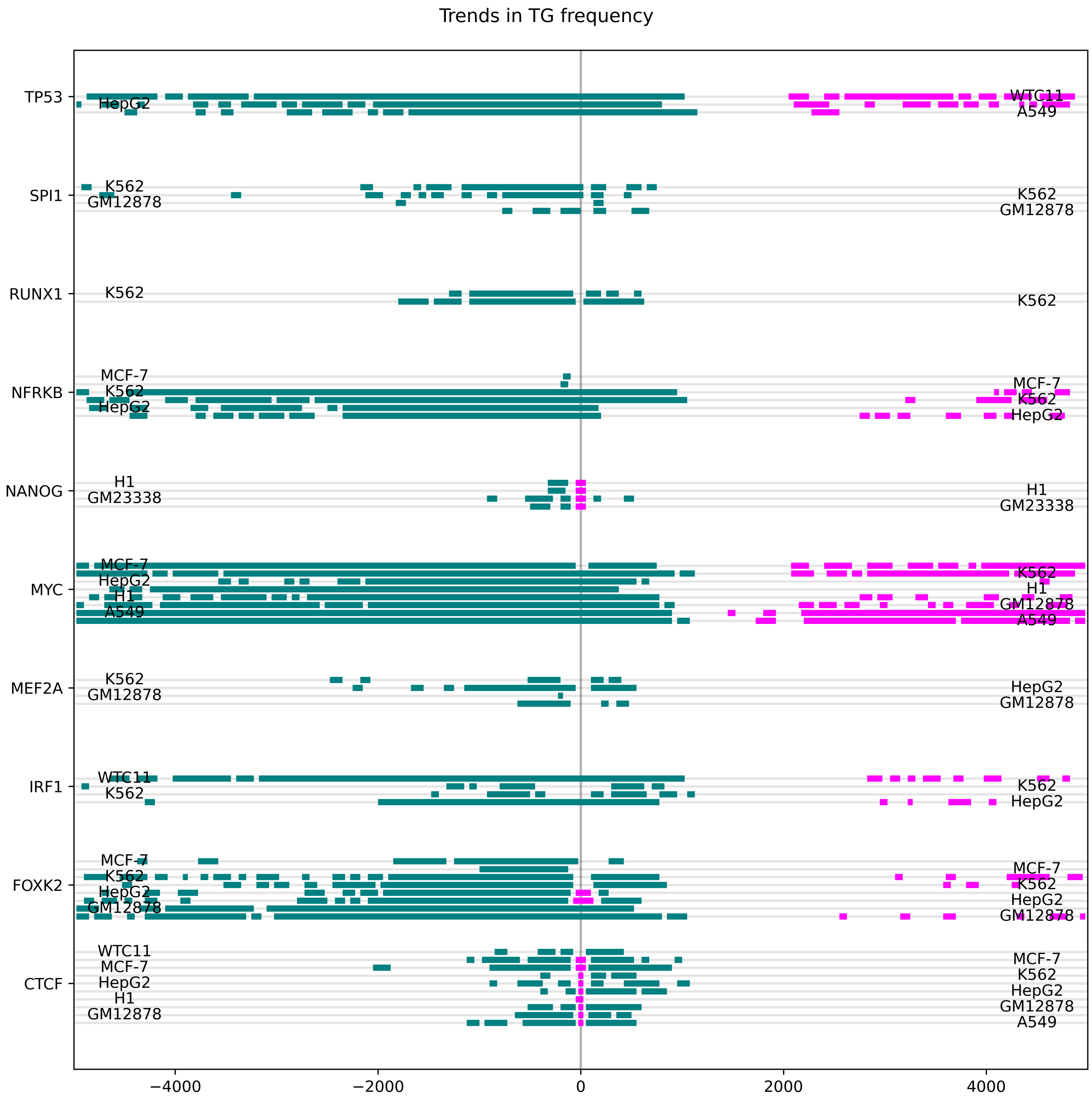


**Fig. S20.** Areas with significant increase or decrease (excessive patches) in frequency of the TG dinucleotide w. r. to the putative binding site position (middle of the experimental peak) for the tested TFs, using the 50 % most relevant experimental peaks. Decrease is depicted in teal, increase in magenta. The excessive patches are estimated using the permutation test.

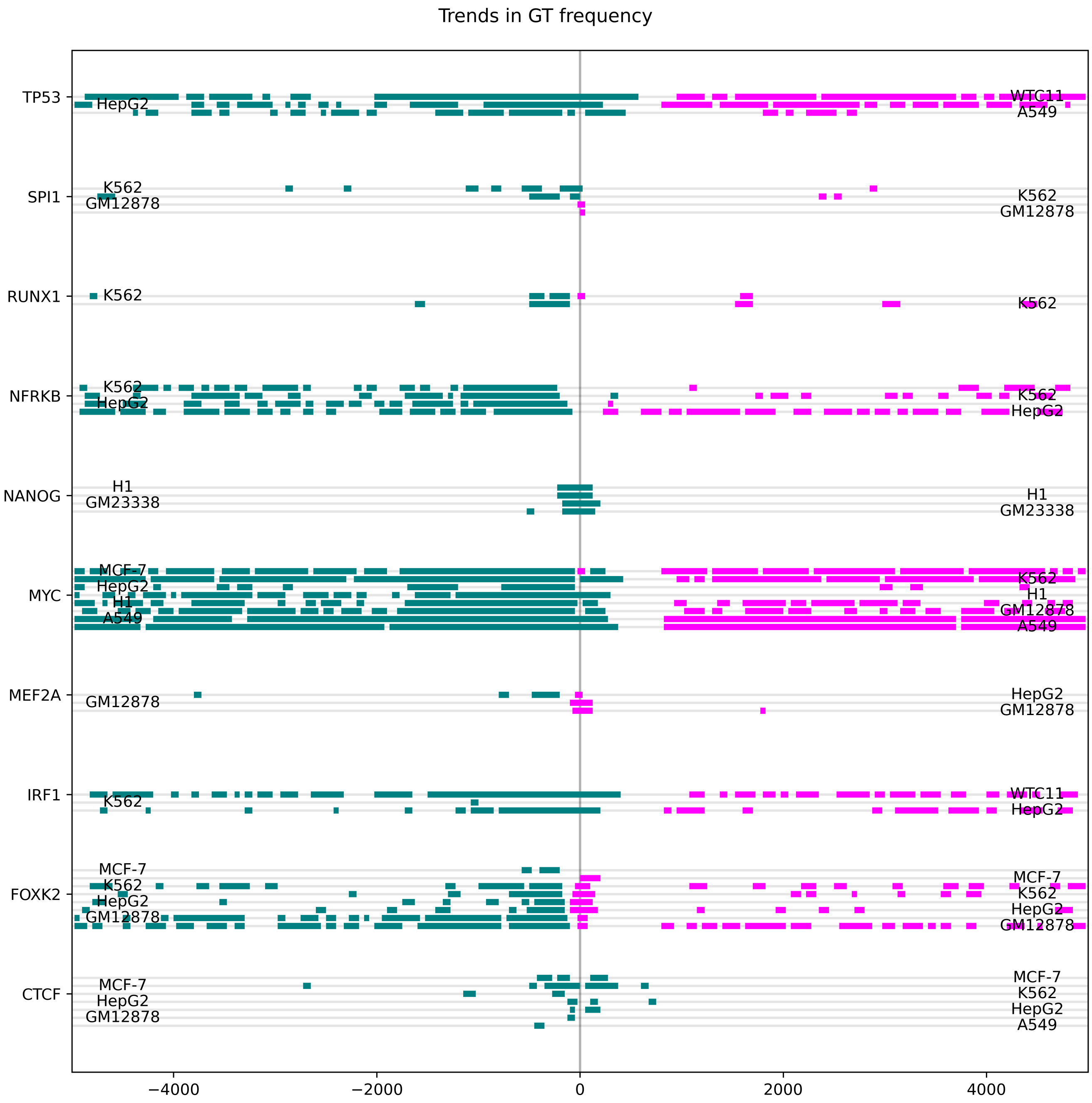


**Fig. S21.** Areas with significant increase or decrease (excessive patches) in frequency of the GT dinucleotide w. r. to the putative binding site position (middle of the experimental peak) for the tested TFs, using the 50 % most relevant experimental peaks. Decrease is depicted in teal, increase in magenta. The excessive patches are estimated using the permutation test.

| Offset | -3000 | -1500 | 0 | 1500 | 3000 |
|---|---|---|---|---|---|
| AA | 1.1 ± 2.5 | 1.0 ± 2.3* | -0.0 ± 1.6**** | 0.7 ± 2.2**** | 0.7 ± 2.2 |
| ANA | 1.1 ± 2.5 | 1.0 ± 2.3*** | -0.0 ± 1.6**** | 0.7 ± 2.2**** | 0.7 ± 2.2 |
| ANNA | 0.9 ± 2.2 | 0.8 ± 2.1*** | -0.2 ± 1.4**** | 0.5 ± 2.0**** | 0.5 ± 2.0 |
| ANNNA | 0.9 ± 2.3 | 0.7 ± 2.1*** | -0.2 ± 1.5**** | 0.5 ± 2.0**** | 0.5 ± 2.0 |
| ANNNNA | 0.8 ± 2.1 | 0.7 ± 2.0*** | -0.3 ± 1.4**** | 0.4 ± 1.9**** | 0.4 ± 1.9 |
| ANNNNNA | 0.8 ± 2.2 | 0.7 ± 2.0**** | -0.2 ± 1.4**** | 0.4 ± 1.9**** | 0.4 ± 1.9 |
| AC | 0.1 ± 1.1 | 0.1 ± 1.1 | -0.1 ± 1.1**** | -0.2 ± 1.0**** | -0.2 ± 1.0 |
| ANC | 0.2 ± 1.1 | 0.2 ± 1.1 | 0.1 ± 1.0**** | -0.1 ± 1.0**** | -0.1 ± 1.0 |
| ANNC | 0.3 ± 1.1 | 0.3 ± 1.1 | 0.1 ± 1.1**** | 0.0 ± 1.0**** | 0.0 ± 1.0 |
| ANNNC | 0.2 ± 1.1 | 0.2 ± 1.1 | 0.1 ± 1.0**** | -0.0 ± 1.0**** | -0.0 ± 1.0 |
| ANNNNC | 0.1 ± 1.1 | 0.2 ± 1.0 | 0.1 ± 1.0**** | -0.1 ± 1.0**** | -0.1 ± 1.0 |
| ANNNNNC | -0.0 ± 1.0 | 0.0 ± 1.0** | -0.1 ± 1.0**** | -0.2 ± 0.9**** | -0.2 ± 1.0 |
| CA | 0.8 ± 1.2 | 0.8 ± 1.2*** | 0.5 ± 1.2**** | 0.5 ± 1.1 | 0.5 ± 1.1** |
| CNA | 0.1 ± 1.1 | 0.1 ± 1.0 | -0.0 ± 1.0**** | -0.1 ± 1.0**** | -0.1 ± 1.0 |
| CNNA | 0.2 ± 1.1 | 0.2 ± 1.1 | 0.1 ± 1.1**** | -0.1 ± 1.0**** | -0.1 ± 1.0 |
| CNNNA | 0.2 ± 1.0 | 0.2 ± 1.1 | 0.1 ± 1.0**** | -0.1 ± 1.0**** | -0.1 ± 1.0 |
| CNNNNA | 0.1 ± 1.0 | 0.1 ± 1.0 | 0.0 ± 1.0**** | -0.1 ± 1.0**** | -0.1 ± 1.0 |
| CNNNNNA | 0.0 ± 1.0 | 0.1 ± 1.0 | -0.1 ± 1.0**** | -0.2 ± 0.9**** | -0.2 ± 0.9 |
| TT | 0.9 ± 2.3 | 0.8 ± 2.2* | -0.1 ± 1.5**** | 0.9 ± 2.3**** | 0.9 ± 2.4* |
| TNT | 0.9 ± 2.3 | 0.8 ± 2.2** | -0.1 ± 1.5**** | 0.9 ± 2.3**** | 1.0 ± 2.4* |
| TNNT | 0.7 ± 2.1 | 0.6 ± 2.0*** | -0.2 ± 1.4**** | 0.6 ± 2.1**** | 0.7 ± 2.2** |
| TNNNT | 0.7 ± 2.1 | 0.6 ± 2.0** | -0.2 ± 1.4**** | 0.7 ± 2.1**** | 0.7 ± 2.3* |
| TNNNNT | 0.6 ± 2.0 | 0.5 ± 1.9** | -0.3 ± 1.3**** | 0.6 ± 2.0**** | 0.6 ± 2.1* |
| TNNNNNT | 0.6 ± 2.0 | 0.5 ± 1.9** | -0.2 ± 1.4**** | 0.6 ± 2.0**** | 0.6 ± 2.1 |
| GT | -0.1 ± 1.0 | -0.1 ± 1.0 | -0.1 ± 1.0 | -0.1 ± 1.0 | -0.1 ± 1.0 |
| GNT | 0.1 ± 1.0 | 0.0 ± 1.0 | 0.1 ± 1.0 | 0.0 ± 1.0 | 0.0 ± 1.0 |
| GNNT | 0.1 ± 1.1 | 0.1 ± 1.0 | 0.2 ± 1.1 | 0.2 ± 1.1 | 0.1 ± 1.0** |
| GNNNT | 0.1 ± 1.0 | 0.1 ± 1.0 | 0.1 ± 1.0 | 0.1 ± 1.0 | 0.1 ± 1.0 |
| GNNNNT | 0.0 ± 1.0 | 0.0 ± 1.0 | 0.1 ± 1.0* | 0.0 ± 1.0** | -0.0 ± 1.0 |
| GNNNNNT | -0.1 ± 1.0 | -0.1 ± 1.0 | -0.1 ± 1.0* | -0.1 ± 1.0 | -0.1 ± 1.0 |
| TG | 0.7 ± 1.2 | 0.6 ± 1.1*** | 0.5 ± 1.2**** | 0.6 ± 1.2**** | 0.7 ± 1.2* |
| TNG | 0.0 ± 1.0 | -0.0 ± 1.0 | -0.0 ± 1.0 | -0.0 ± 1.0 | 0.0 ± 1.0 |
| TNNG | 0.1 ± 1.0 | 0.1 ± 1.0 | 0.1 ± 1.1 | 0.1 ± 1.0 | 0.1 ± 1.1 |
| TNNNG | 0.0 ± 1.0 | 0.1 ± 1.0 | 0.0 ± 1.0 | 0.0 ± 1.0 | 0.0 ± 1.0 |
| TNNNNG | -0.0 ± 1.0 | -0.0 ± 1.0 | 0.0 ± 1.0 | -0.0 ± 1.0* | -0.0 ± 1.0 |
| TNNNNNG | -0.1 ± 1.0 | -0.1 ± 1.0 | -0.1 ± 1.0 | -0.1 ± 1.0 | -0.1 ± 1.0 |

**Table S5.** Average dinucleotide count z-score in a 50 bp k-mer centered at the specified offset for experiment ENCFF043WTJ, a ChIP-seq experiment targetting MYC in K562. Stars indicate whether the average count is significantly different from the preceding offset in this table (Welch t-test, 0.0001, indicated by "****", 0.001 by "***", 0.01 by "**" and 0.05 by"*"). Addendum to Figure 3.2. As background distribution for z-score normalization was acquired by scrambling available k-mer sequences at a given offset and identifying dinucleotide frequencies in these scrambled sequences. The process mirrors calculating scramble-normalized TF affinity (see Methods) to keep base pair frequencies but not their order.

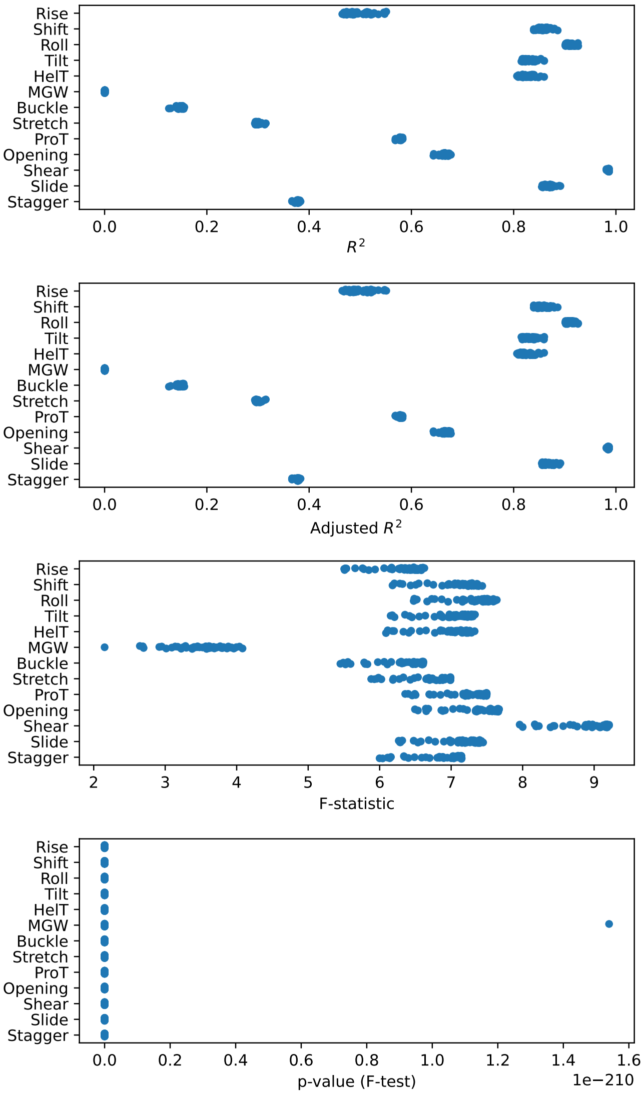


**Fig. S22.** Coefficients of determination and model statistics for all trained linear models by shape feature. Panels show (from top to bottom): the coefficient of determination ($R^2$), adjusted coefficient of determination (adjusted $R^2$), F-statistic values, and the corresponding p-values of the F-statistic for each model.

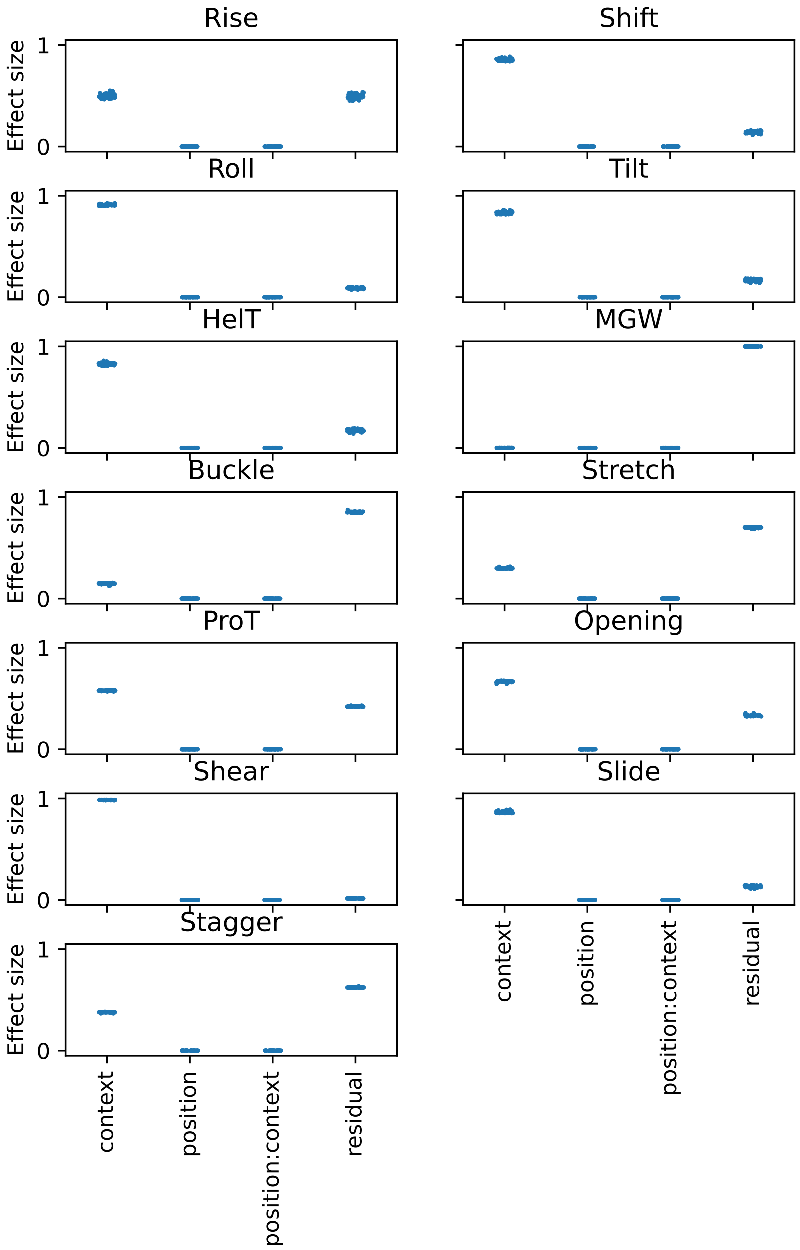


**Fig. S23.** Effect size of different variables. One point corresponds to one experiment. Each shape feature was modeled as a function of sequence context (defined as either a single base pair or a pair of neighboring base pairs), position relative to the binding site, and their interaction (position:context on the plot). ANOVA tables were generated using the *anova_lm* function from the *statsmodels* package, and effect size ($\eta^2$) was calculated as the ratio of the variable sum of squares to the total sum of squares.

| | | -3000 bp | -1500 bp | at binding site | 1500 bp | 3000 bp |
|---|---|---|---|---|---|---|
| CTCF | A | -9.51 ± 2.38 | -9.51 ± 2.37 | -8.44 ± 2.07 | -9.49 ± 2.38 | -9.52 ± 2.39 |
| | C | -4.11 ± 2.21 | -4.07 ± 2.21 | -3.72 ± 2.02 | -4.09 ± 2.21 | -4.11 ± 2.22 |
| | G | -4.1 ± 2.22 | -4.07 ± 2.19 | -3.7 ± 2.0 | -4.09 ± 2.21 | -4.08 ± 2.22 |
| | T | -9.51 ± 2.38 | -9.49 ± 2.38 | -8.46 ± 2.08 | -9.49 ± 2.37 | -9.52 ± 2.38 |
| FOXK2 | A | -9.54 ± 2.43 | -9.52 ± 2.42 | -9.18 ± 2.21 | -9.47 ± 2.38 | -9.51 ± 2.41 |
| | C | -4.1 ± 2.21 | -4.11 ± 2.23 | -4.22 ± 2.28 | -4.1 ± 2.2 | -4.12 ± 2.2 |
| | G | -4.1 ± 2.16 | -4.05 ± 2.18 | -4.26 ± 2.26 | -4.09 ± 2.2 | -4.13 ± 2.21 |
| | T | -9.53 ± 2.38 | -9.51 ± 2.39 | -9.22 ± 2.23 | -9.56 ± 2.41 | -9.56 ± 2.44 |
| IRF1 | A | -9.59 ± 2.41 | -9.55 ± 2.37 | -9.3 ± 2.32 | -9.53 ± 2.38 | -9.47 ± 2.37 |
| | C | -4.16 ± 2.2 | -4.03 ± 2.18 | -3.94 ± 2.09 | -4.09 ± 2.21 | -4.09 ± 2.16 |
| | G | -4.05 ± 2.2 | -4.04 ± 2.17 | -3.93 ± 2.07 | -4.16 ± 2.2 | -4.16 ± 2.26 |
| | T | -9.58 ± 2.37 | -9.56 ± 2.41 | -9.19 ± 2.3 | -9.59 ± 2.41 | -9.58 ± 2.44 |
| MEF2A | A | -9.57 ± 2.38 | -9.57 ± 2.41 | -9.5 ± 2.33 | -9.61 ± 2.42 | -9.63 ± 2.4 |
| | C | -4.16 ± 2.2 | -4.11 ± 2.18 | -4.2 ± 2.19 | -4.12 ± 2.21 | -4.16 ± 2.23 |
| | G | -4.11 ± 2.2 | -4.12 ± 2.22 | -4.15 ± 2.18 | -4.19 ± 2.24 | -4.16 ± 2.19 |
| | T | -9.59 ± 2.38 | -9.56 ± 2.4 | -9.55 ± 2.35 | -9.61 ± 2.39 | -9.6 ± 2.38 |
| MYC | A | -9.56 ± 2.44 | -9.46 ± 2.43 | -8.61 ± 2.08 | -9.43 ± 2.37 | -9.44 ± 2.42 |
| | C | -4.1 ± 2.24 | -4.06 ± 2.18 | -3.78 ± 2.0 | -4.01 ± 2.15 | -4.03 ± 2.21 |
| | G | -4.03 ± 2.17 | -3.98 ± 2.16 | -3.85 ± 2.03 | -4.05 ± 2.17 | -4.15 ± 2.22 |
| | T | -9.4 ± 2.39 | -9.44 ± 2.41 | -8.55 ± 2.11 | -9.55 ± 2.46 | -9.5 ± 2.42 |
| NANOG | A | -9.74 ± 2.38 | -9.68 ± 2.37 | -9.49 ± 2.21 | -9.68 ± 2.34 | -9.69 ± 2.36 |
| | C | -4.25 ± 2.21 | -4.2 ± 2.21 | -4.15 ± 2.23 | -4.16 ± 2.23 | -4.2 ± 2.24 |
| | G | -4.18 ± 2.2 | -4.13 ± 2.21 | -4.09 ± 2.24 | -4.18 ± 2.27 | -4.31 ± 2.28 |
| | T | -9.74 ± 2.39 | -9.68 ± 2.37 | -9.55 ± 2.2 | -9.69 ± 2.37 | -9.73 ± 2.36 |
| NFRKB | A | -9.61 ± 2.44 | -9.6 ± 2.43 | -9.21 ± 2.3 | -9.56 ± 2.38 | -9.56 ± 2.39 |
| | C | -4.19 ± 2.22 | -4.23 ± 2.23 | -4.02 ± 2.16 | -4.1 ± 2.21 | -4.09 ± 2.21 |
| | G | -4.13 ± 2.19 | -4.08 ± 2.19 | -4.06 ± 2.13 | -4.14 ± 2.2 | -4.15 ± 2.25 |
| | T | -9.55 ± 2.43 | -9.49 ± 2.37 | -9.18 ± 2.26 | -9.59 ± 2.4 | -9.66 ± 2.42 |
| RUNX1 | A | -9.33 ± 2.46 | -9.46 ± 2.4 | -8.77 ± 2.26 | -9.3 ± 2.34 | -9.31 ± 2.42 |
| | C | -4.08 ± 2.13 | -3.97 ± 2.1 | -3.84 ± 2.05 | -3.98 ± 2.19 | -4.06 ± 2.21 |
| | G | -4.03 ± 2.2 | -3.98 ± 2.2 | -3.88 ± 2.04 | -4.07 ± 2.16 | -4.12 ± 2.19 |
| | T | -9.48 ± 2.43 | -9.36 ± 2.35 | -8.9 ± 2.1 | -9.32 ± 2.4 | -9.45 ± 2.37 |
| SPI1 | A | -9.53 ± 2.38 | -9.52 ± 2.38 | -9.18 ± 2.22 | -9.52 ± 2.38 | -9.58 ± 2.4 |
| | C | -4.1 ± 2.21 | -4.13 ± 2.21 | -3.93 ± 2.08 | -4.11 ± 2.22 | -4.14 ± 2.24 |
| | G | -4.16 ± 2.2 | -4.13 ± 2.22 | -3.92 ± 2.11 | -4.19 ± 2.25 | -4.14 ± 2.23 |
| | T | -9.56 ± 2.38 | -9.54 ± 2.39 | -9.14 ± 2.2 | -9.57 ± 2.4 | -9.6 ± 2.38 |
| TP53 | A | -9.56 ± 2.45 | -9.47 ± 2.43 | -8.63 ± 2.16 | -9.39 ± 2.37 | -9.48 ± 2.42 |
| | C | -4.06 ± 2.17 | -4.05 ± 2.21 | -3.74 ± 1.95 | -3.98 ± 2.13 | -3.99 ± 2.19 |
| | G | -4.07 ± 2.18 | -3.99 ± 2.15 | -3.77 ± 1.98 | -4.06 ± 2.2 | -4.08 ± 2.21 |
| | T | -9.43 ± 2.41 | -9.38 ± 2.4 | -8.65 ± 2.15 | -9.47 ± 2.41 | -9.52 ± 2.39 |

**Table S6.** Average propeller twist values from selected positions relative to the binding site (prolonged peak center). The shape feature values were acquired using deepDNAshape from 50 bp windows.

| | | -3000 bp | -1500 bp | at binding site | 1500 bp | 3000 bp |
|---|---|---|---|---|---|---|
| CTCF | A | $5.1 \pm 0.65$ | $5.1 \pm 0.65$ | $5.28 \pm 0.52$ | $5.1 \pm 0.65$ | $5.09 \pm 0.65$ |
| | C | $5.11 \pm 0.5$ | $5.11 \pm 0.49$ | $5.17 \pm 0.41$ | $5.11 \pm 0.5$ | $5.11 \pm 0.5$ |
| | G | $5.11 \pm 0.5$ | $5.11 \pm 0.5$ | $5.17 \pm 0.42$ | $5.11 \pm 0.5$ | $5.11 \pm 0.49$ |
| | T | $5.1 \pm 0.66$ | $5.1 \pm 0.65$ | $5.27 \pm 0.53$ | $5.11 \pm 0.66$ | $5.11 \pm 0.65$ |
| FOXK2 | A | $5.09 \pm 0.66$ | $5.09 \pm 0.66$ | $5.18 \pm 0.58$ | $5.1 \pm 0.65$ | $5.09 \pm 0.66$ |
| | C | $5.11 \pm 0.5$ | $5.11 \pm 0.49$ | $5.16 \pm 0.46$ | $5.11 \pm 0.49$ | $5.11 \pm 0.5$ |
| | G | $5.11 \pm 0.5$ | $5.11 \pm 0.5$ | $5.17 \pm 0.47$ | $5.11 \pm 0.49$ | $5.11 \pm 0.49$ |
| | T | $5.09 \pm 0.66$ | $5.08 \pm 0.66$ | $5.18 \pm 0.58$ | $5.09 \pm 0.66$ | $5.1 \pm 0.66$ |
| IRF1 | A | $5.08 \pm 0.67$ | $5.1 \pm 0.66$ | $5.1 \pm 0.62$ | $5.09 \pm 0.66$ | $5.1 \pm 0.66$ |
| | C | $5.11 \pm 0.5$ | $5.11 \pm 0.49$ | $5.14 \pm 0.48$ | $5.12 \pm 0.49$ | $5.12 \pm 0.5$ |
| | G | $5.11 \pm 0.5$ | $5.1 \pm 0.5$ | $5.13 \pm 0.47$ | $5.11 \pm 0.5$ | $5.1 \pm 0.51$ |
| | T | $5.08 \pm 0.66$ | $5.09 \pm 0.67$ | $5.15 \pm 0.6$ | $5.09 \pm 0.67$ | $5.09 \pm 0.67$ |
| MEF2A | A | $5.1 \pm 0.67$ | $5.1 \pm 0.66$ | $5.1 \pm 0.63$ | $5.1 \pm 0.67$ | $5.08 \pm 0.66$ |
| | C | $5.11 \pm 0.51$ | $5.11 \pm 0.5$ | $5.14 \pm 0.5$ | $5.1 \pm 0.51$ | $5.11 \pm 0.51$ |
| | G | $5.11 \pm 0.51$ | $5.11 \pm 0.5$ | $5.13 \pm 0.5$ | $5.12 \pm 0.51$ | $5.12 \pm 0.51$ |
| | T | $5.08 \pm 0.67$ | $5.08 \pm 0.66$ | $5.1 \pm 0.64$ | $5.08 \pm 0.67$ | $5.09 \pm 0.66$ |
| MYC | A | $5.09 \pm 0.66$ | $5.1 \pm 0.65$ | $5.23 \pm 0.54$ | $5.09 \pm 0.66$ | $5.1 \pm 0.66$ |
| | C | $5.12 \pm 0.49$ | $5.12 \pm 0.49$ | $5.19 \pm 0.41$ | $5.12 \pm 0.48$ | $5.11 \pm 0.5$ |
| | G | $5.12 \pm 0.5$ | $5.12 \pm 0.48$ | $5.2 \pm 0.41$ | $5.12 \pm 0.49$ | $5.11 \pm 0.5$ |
| | T | $5.11 \pm 0.65$ | $5.09 \pm 0.65$ | $5.24 \pm 0.54$ | $5.09 \pm 0.67$ | $5.11 \pm 0.65$ |
| NANOG | A | $5.07 \pm 0.67$ | $5.09 \pm 0.67$ | $5.13 \pm 0.6$ | $5.1 \pm 0.66$ | $5.09 \pm 0.67$ |
| | C | $5.11 \pm 0.52$ | $5.1 \pm 0.52$ | $5.13 \pm 0.51$ | $5.1 \pm 0.51$ | $5.09 \pm 0.52$ |
| | G | $5.11 \pm 0.52$ | $5.09 \pm 0.51$ | $5.13 \pm 0.5$ | $5.1 \pm 0.52$ | $5.1 \pm 0.54$ |
| | T | $5.07 \pm 0.68$ | $5.08 \pm 0.68$ | $5.13 \pm 0.6$ | $5.07 \pm 0.67$ | $5.08 \pm 0.67$ |
| NFRKB | A | $5.08 \pm 0.66$ | $5.09 \pm 0.66$ | $5.13 \pm 0.6$ | $5.09 \pm 0.66$ | $5.08 \pm 0.66$ |
| | C | $5.1 \pm 0.5$ | $5.12 \pm 0.5$ | $5.14 \pm 0.46$ | $5.11 \pm 0.5$ | $5.11 \pm 0.5$ |
| | G | $5.1 \pm 0.5$ | $5.11 \pm 0.5$ | $5.13 \pm 0.47$ | $5.12 \pm 0.51$ | $5.11 \pm 0.51$ |
| | T | $5.09 \pm 0.66$ | $5.09 \pm 0.66$ | $5.14 \pm 0.61$ | $5.09 \pm 0.67$ | $5.08 \pm 0.67$ |
| RUNX1 | A | $5.09 \pm 0.65$ | $5.13 \pm 0.65$ | $5.25 \pm 0.56$ | $5.15 \pm 0.62$ | $5.11 \pm 0.65$ |
| | C | $5.12 \pm 0.52$ | $5.1 \pm 0.49$ | $5.17 \pm 0.45$ | $5.11 \pm 0.48$ | $5.11 \pm 0.49$ |
| | G | $5.11 \pm 0.49$ | $5.1 \pm 0.48$ | $5.17 \pm 0.45$ | $5.11 \pm 0.49$ | $5.16 \pm 0.48$ |
| | T | $5.1 \pm 0.66$ | $5.14 \pm 0.65$ | $5.25 \pm 0.53$ | $5.11 \pm 0.64$ | $5.12 \pm 0.63$ |
| SPI1 | A | $5.09 \pm 0.66$ | $5.1 \pm 0.66$ | $5.07 \pm 0.58$ | $5.11 \pm 0.66$ | $5.08 \pm 0.66$ |
| | C | $5.12 \pm 0.5$ | $5.11 \pm 0.5$ | $5.09 \pm 0.48$ | $5.11 \pm 0.5$ | $5.11 \pm 0.5$ |
| | G | $5.11 \pm 0.51$ | $5.11 \pm 0.5$ | $5.09 \pm 0.48$ | $5.11 \pm 0.51$ | $5.11 \pm 0.5$ |
| | T | $5.09 \pm 0.67$ | $5.1 \pm 0.66$ | $5.07 \pm 0.58$ | $5.1 \pm 0.66$ | $5.1 \pm 0.66$ |
| TP53 | A | $5.09 \pm 0.67$ | $5.1 \pm 0.65$ | $5.21 \pm 0.55$ | $5.11 \pm 0.64$ | $5.09 \pm 0.65$ |
| | C | $5.12 \pm 0.49$ | $5.11 \pm 0.49$ | $5.18 \pm 0.43$ | $5.11 \pm 0.49$ | $5.12 \pm 0.49$ |
| | G | $5.12 \pm 0.49$ | $5.12 \pm 0.49$ | $5.18 \pm 0.42$ | $5.11 \pm 0.48$ | $5.12 \pm 0.5$ |
| | T | $5.11 \pm 0.65$ | $5.1 \pm 0.66$ | $5.21 \pm 0.55$ | $5.1 \pm 0.65$ | $5.09 \pm 0.67$ |

**Table S7.** Average minor groove width values from selected positions relative to the binding site (prolonged peak center). The shape feature values were acquired using deepDNAshape from 50 bp windows.

| | | -3000 bp | -1500 bp | at binding site | 1500 bp | 3000 bp |
|---|---|---|---|---|---|---|
| CTCF | A | -0.88 ± 0.55 | -0.88 ± 0.55 | -0.73 ± 0.46 | -0.87 ± 0.55 | -0.88 ± 0.55 |
| | C | 0.31 ± 0.14 | 0.31 ± 0.14 | 0.3 ± 0.13 | 0.31 ± 0.14 | 0.31 ± 0.14 |
| | G | 0.31 ± 0.15 | 0.31 ± 0.14 | 0.31 ± 0.13 | 0.31 ± 0.15 | 0.31 ± 0.14 |
| | T | -0.87 ± 0.55 | -0.87 ± 0.55 | -0.74 ± 0.46 | -0.88 ± 0.55 | -0.88 ± 0.55 |
| FOXK2 | A | -0.89 ± 0.56 | -0.88 ± 0.56 | -0.85 ± 0.52 | -0.87 ± 0.56 | -0.88 ± 0.55 |
| | C | 0.31 ± 0.15 | 0.31 ± 0.15 | 0.32 ± 0.14 | 0.31 ± 0.14 | 0.31 ± 0.15 |
| | G | 0.31 ± 0.14 | 0.31 ± 0.15 | 0.32 ± 0.15 | 0.31 ± 0.15 | 0.31 ± 0.14 |
| | T | -0.88 ± 0.56 | -0.88 ± 0.56 | -0.85 ± 0.52 | -0.89 ± 0.56 | -0.9 ± 0.56 |
| IRF1 | A | -0.89 ± 0.56 | -0.89 ± 0.55 | -0.84 ± 0.53 | -0.88 ± 0.56 | -0.87 ± 0.55 |
| | C | 0.31 ± 0.14 | 0.31 ± 0.15 | 0.31 ± 0.14 | 0.31 ± 0.15 | 0.31 ± 0.14 |
| | G | 0.31 ± 0.15 | 0.31 ± 0.14 | 0.31 ± 0.14 | 0.31 ± 0.15 | 0.31 ± 0.15 |
| | T | -0.88 ± 0.56 | -0.89 ± 0.56 | -0.84 ± 0.53 | -0.89 ± 0.57 | -0.89 ± 0.56 |
| MEF2A | A | -0.89 ± 0.56 | -0.9 ± 0.56 | -0.88 ± 0.54 | -0.9 ± 0.56 | -0.9 ± 0.56 |
| | C | 0.31 ± 0.14 | 0.31 ± 0.14 | 0.31 ± 0.15 | 0.31 ± 0.14 | 0.31 ± 0.15 |
| | G | 0.31 ± 0.14 | 0.31 ± 0.15 | 0.31 ± 0.15 | 0.31 ± 0.15 | 0.31 ± 0.15 |
| | T | -0.89 ± 0.55 | -0.88 ± 0.57 | -0.89 ± 0.54 | -0.89 ± 0.56 | -0.89 ± 0.55 |
| MYC | A | -0.9 ± 0.57 | -0.88 ± 0.56 | -0.76 ± 0.47 | -0.86 ± 0.55 | -0.86 ± 0.56 |
| | C | 0.31 ± 0.14 | 0.31 ± 0.14 | 0.32 ± 0.13 | 0.31 ± 0.14 | 0.31 ± 0.14 |
| | G | 0.31 ± 0.14 | 0.31 ± 0.14 | 0.32 ± 0.13 | 0.31 ± 0.14 | 0.31 ± 0.15 |
| | T | -0.87 ± 0.55 | -0.86 ± 0.56 | -0.75 ± 0.47 | -0.9 ± 0.57 | -0.89 ± 0.56 |
| NANOG | A | -0.91 ± 0.56 | -0.91 ± 0.55 | -0.89 ± 0.52 | -0.9 ± 0.55 | -0.9 ± 0.56 |
| | C | 0.31 ± 0.15 | 0.31 ± 0.15 | 0.31 ± 0.14 | 0.31 ± 0.15 | 0.31 ± 0.14 |
| | G | 0.31 ± 0.14 | 0.31 ± 0.15 | 0.31 ± 0.14 | 0.31 ± 0.15 | 0.32 ± 0.15 |
| | T | -0.91 ± 0.56 | -0.9 ± 0.55 | -0.89 ± 0.52 | -0.9 ± 0.55 | -0.91 ± 0.56 |
| NFRKB | A | -0.9 ± 0.57 | -0.9 ± 0.56 | -0.84 ± 0.54 | -0.88 ± 0.55 | -0.89 ± 0.55 |
| | C | 0.32 ± 0.15 | 0.31 ± 0.15 | 0.32 ± 0.14 | 0.31 ± 0.15 | 0.31 ± 0.15 |
| | G | 0.31 ± 0.14 | 0.31 ± 0.15 | 0.32 ± 0.14 | 0.31 ± 0.15 | 0.31 ± 0.15 |
| | T | -0.89 ± 0.56 | -0.87 ± 0.55 | -0.82 ± 0.53 | -0.9 ± 0.56 | -0.91 ± 0.57 |
| RUNX1 | A | -0.84 ± 0.56 | -0.89 ± 0.57 | -0.81 ± 0.5 | -0.87 ± 0.55 | -0.85 ± 0.55 |
| | C | 0.3 ± 0.14 | 0.31 ± 0.15 | 0.31 ± 0.13 | 0.31 ± 0.14 | 0.31 ± 0.14 |
| | G | 0.31 ± 0.14 | 0.31 ± 0.14 | 0.32 ± 0.14 | 0.32 ± 0.15 | 0.31 ± 0.15 |
| | T | -0.88 ± 0.55 | -0.88 ± 0.54 | -0.83 ± 0.5 | -0.85 ± 0.53 | -0.88 ± 0.54 |
| SPI1 | A | -0.88 ± 0.56 | -0.88 ± 0.54 | -0.81 ± 0.53 | -0.89 ± 0.55 | -0.89 ± 0.56 |
| | C | 0.31 ± 0.14 | 0.31 ± 0.15 | 0.31 ± 0.14 | 0.31 ± 0.14 | 0.31 ± 0.15 |
| | G | 0.31 ± 0.14 | 0.31 ± 0.14 | 0.31 ± 0.13 | 0.32 ± 0.15 | 0.31 ± 0.15 |
| | T | -0.88 ± 0.55 | -0.89 ± 0.55 | -0.8 ± 0.52 | -0.9 ± 0.55 | -0.89 ± 0.55 |
| TP53 | A | -0.89 ± 0.57 | -0.88 ± 0.56 | -0.75 ± 0.49 | -0.87 ± 0.55 | -0.87 ± 0.56 |
| | C | 0.31 ± 0.15 | 0.32 ± 0.15 | 0.32 ± 0.13 | 0.31 ± 0.14 | 0.31 ± 0.14 |
| | G | 0.31 ± 0.14 | 0.31 ± 0.14 | 0.32 ± 0.13 | 0.31 ± 0.14 | 0.31 ± 0.15 |
| | T | -0.87 ± 0.55 | -0.86 ± 0.55 | -0.75 ± 0.48 | -0.88 ± 0.56 | -0.88 ± 0.55 |

**Table S8.** Average opening values from selected positions relative to the binding site (prolonged peak center). The shape feature values were acquired using deepDNAshape from 50 bp windows.

| | | -3000 bp | -1500 bp | at binding site | 1500 bp | 3000 bp |
|---|---|---|---|---|---|---|
| CTCF | A | -0.02 ± 0.01 | -0.02 ± 0.01 | -0.02 ± 0.01 | -0.02 ± 0.01 | -0.02 ± 0.01 |
| | C | -0.03 ± 0.01 | -0.03 ± 0.01 | -0.03 ± 0.01 | -0.03 ± 0.01 | -0.03 ± 0.01 |
| | G | -0.03 ± 0.01 | -0.03 ± 0.01 | -0.03 ± 0.01 | -0.03 ± 0.01 | -0.03 ± 0.01 |
| | T | -0.02 ± 0.01 | -0.02 ± 0.01 | -0.02 ± 0.01 | -0.02 ± 0.01 | -0.02 ± 0.01 |
| FOXK2 | A | -0.02 ± 0.01 | -0.02 ± 0.01 | -0.02 ± 0.01 | -0.02 ± 0.01 | -0.02 ± 0.01 |
| | C | -0.03 ± 0.01 | -0.03 ± 0.01 | -0.03 ± 0.01 | -0.03 ± 0.01 | -0.03 ± 0.01 |
| | G | -0.03 ± 0.01 | -0.03 ± 0.01 | -0.03 ± 0.01 | -0.03 ± 0.01 | -0.03 ± 0.01 |
| | T | -0.02 ± 0.01 | -0.02 ± 0.01 | -0.02 ± 0.01 | -0.02 ± 0.01 | -0.02 ± 0.01 |
| IRF1 | A | -0.02 ± 0.01 | -0.02 ± 0.01 | -0.02 ± 0.01 | -0.02 ± 0.01 | -0.02 ± 0.01 |
| | C | -0.03 ± 0.01 | -0.03 ± 0.01 | -0.03 ± 0.01 | -0.03 ± 0.01 | -0.03 ± 0.01 |
| | G | -0.03 ± 0.01 | -0.03 ± 0.01 | -0.03 ± 0.01 | -0.03 ± 0.01 | -0.03 ± 0.01 |
| | T | -0.02 ± 0.01 | -0.02 ± 0.01 | -0.02 ± 0.01 | -0.02 ± 0.01 | -0.02 ± 0.01 |
| MEF2A | A | -0.02 ± 0.01 | -0.02 ± 0.01 | -0.02 ± 0.01 | -0.02 ± 0.01 | -0.02 ± 0.01 |
| | C | -0.03 ± 0.01 | -0.03 ± 0.01 | -0.03 ± 0.01 | -0.03 ± 0.01 | -0.03 ± 0.01 |
| | G | -0.03 ± 0.01 | -0.03 ± 0.01 | -0.03 ± 0.01 | -0.03 ± 0.01 | -0.03 ± 0.01 |
| | T | -0.02 ± 0.01 | -0.02 ± 0.01 | -0.02 ± 0.01 | -0.02 ± 0.01 | -0.02 ± 0.01 |
| MYC | A | -0.02 ± 0.01 | -0.02 ± 0.01 | -0.02 ± 0.01 | -0.02 ± 0.01 | -0.02 ± 0.01 |
| | C | -0.03 ± 0.01 | -0.03 ± 0.01 | -0.03 ± 0.01 | -0.03 ± 0.01 | -0.03 ± 0.01 |
| | G | -0.03 ± 0.01 | -0.03 ± 0.01 | -0.03 ± 0.01 | -0.03 ± 0.01 | -0.03 ± 0.01 |
| | T | -0.02 ± 0.01 | -0.02 ± 0.01 | -0.02 ± 0.01 | -0.02 ± 0.01 | -0.02 ± 0.01 |
| NANOG | A | -0.02 ± 0.01 | -0.02 ± 0.01 | -0.02 ± 0.01 | -0.02 ± 0.01 | -0.02 ± 0.01 |
| | C | -0.03 ± 0.01 | -0.03 ± 0.01 | -0.03 ± 0.01 | -0.03 ± 0.01 | -0.03 ± 0.01 |
| | G | -0.03 ± 0.01 | -0.03 ± 0.01 | -0.03 ± 0.01 | -0.03 ± 0.01 | -0.03 ± 0.01 |
| | T | -0.02 ± 0.01 | -0.02 ± 0.01 | -0.02 ± 0.01 | -0.02 ± 0.01 | -0.02 ± 0.01 |
| NFRKB | A | -0.02 ± 0.01 | -0.02 ± 0.01 | -0.02 ± 0.01 | -0.02 ± 0.01 | -0.02 ± 0.01 |
| | C | -0.03 ± 0.01 | -0.03 ± 0.01 | -0.03 ± 0.01 | -0.03 ± 0.01 | -0.03 ± 0.01 |
| | G | -0.03 ± 0.01 | -0.03 ± 0.01 | -0.03 ± 0.01 | -0.03 ± 0.01 | -0.03 ± 0.01 |
| | T | -0.02 ± 0.01 | -0.02 ± 0.01 | -0.02 ± 0.01 | -0.02 ± 0.01 | -0.02 ± 0.01 |
| RUNX1 | A | -0.02 ± 0.01 | -0.02 ± 0.01 | -0.02 ± 0.01 | -0.02 ± 0.01 | -0.02 ± 0.01 |
| | C | -0.03 ± 0.01 | -0.03 ± 0.01 | -0.03 ± 0.01 | -0.03 ± 0.01 | -0.03 ± 0.01 |
| | G | -0.03 ± 0.01 | -0.03 ± 0.01 | -0.03 ± 0.01 | -0.03 ± 0.01 | -0.03 ± 0.01 |
| | T | -0.02 ± 0.01 | -0.02 ± 0.01 | -0.02 ± 0.01 | -0.02 ± 0.01 | -0.02 ± 0.01 |
| SPI1 | A | -0.02 ± 0.01 | -0.02 ± 0.01 | -0.02 ± 0.01 | -0.02 ± 0.01 | -0.02 ± 0.01 |
| | C | -0.03 ± 0.01 | -0.03 ± 0.01 | -0.03 ± 0.01 | -0.03 ± 0.01 | -0.03 ± 0.01 |
| | G | -0.03 ± 0.01 | -0.03 ± 0.01 | -0.03 ± 0.01 | -0.03 ± 0.01 | -0.03 ± 0.01 |
| | T | -0.02 ± 0.01 | -0.02 ± 0.01 | -0.02 ± 0.01 | -0.02 ± 0.01 | -0.02 ± 0.01 |
| TP53 | A | -0.02 ± 0.01 | -0.02 ± 0.01 | -0.02 ± 0.01 | -0.02 ± 0.01 | -0.02 ± 0.01 |
| | C | -0.03 ± 0.01 | -0.03 ± 0.01 | -0.03 ± 0.01 | -0.03 ± 0.01 | -0.03 ± 0.01 |
| | G | -0.03 ± 0.01 | -0.03 ± 0.01 | -0.03 ± 0.01 | -0.03 ± 0.01 | -0.03 ± 0.01 |
| | T | -0.02 ± 0.01 | -0.02 ± 0.01 | -0.02 ± 0.01 | -0.02 ± 0.01 | -0.02 ± 0.01 |

**Table S9.** Average stretch values from selected positions relative to the binding site (prolonged peak center). The shape feature values were acquired using deepDNAshape from 50 bp windows.

| | | -3000 bp | -1500 bp | at binding site | 1500 bp | 3000 bp |
|---|---|---|---|---|---|---|
| CTCF | A | -0.08 ± 0.11 | -0.08 ± 0.11 | -0.07 ± 0.1 | -0.08 ± 0.11 | -0.08 ± 0.11 |
| | C | 0.06 ± 0.07 | 0.06 ± 0.07 | 0.06 ± 0.07 | 0.06 ± 0.07 | 0.06 ± 0.07 |
| | G | 0.06 ± 0.07 | 0.06 ± 0.07 | 0.06 ± 0.07 | 0.06 ± 0.07 | 0.06 ± 0.07 |
| | T | -0.08 ± 0.11 | -0.08 ± 0.11 | -0.07 ± 0.1 | -0.08 ± 0.11 | -0.08 ± 0.11 |
| FOXK2 | A | -0.08 ± 0.11 | -0.08 ± 0.11 | -0.07 ± 0.11 | -0.08 ± 0.11 | -0.08 ± 0.11 |
| | C | 0.06 ± 0.07 | 0.06 ± 0.07 | 0.06 ± 0.07 | 0.06 ± 0.07 | 0.06 ± 0.07 |
| | G | 0.06 ± 0.07 | 0.06 ± 0.07 | 0.06 ± 0.07 | 0.06 ± 0.07 | 0.06 ± 0.07 |
| | T | -0.08 ± 0.11 | -0.08 ± 0.11 | -0.07 ± 0.11 | -0.08 ± 0.11 | -0.08 ± 0.11 |
| IRF1 | A | -0.08 ± 0.11 | -0.08 ± 0.11 | -0.08 ± 0.11 | -0.08 ± 0.11 | -0.08 ± 0.11 |
| | C | 0.06 ± 0.07 | 0.06 ± 0.07 | 0.06 ± 0.07 | 0.06 ± 0.07 | 0.06 ± 0.07 |
| | G | 0.06 ± 0.07 | 0.06 ± 0.07 | 0.06 ± 0.07 | 0.06 ± 0.07 | 0.06 ± 0.07 |
| | T | -0.08 ± 0.11 | -0.08 ± 0.11 | -0.08 ± 0.11 | -0.08 ± 0.11 | -0.08 ± 0.11 |
| MEF2A | A | -0.08 ± 0.11 | -0.08 ± 0.11 | -0.08 ± 0.11 | -0.08 ± 0.11 | -0.08 ± 0.11 |
| | C | 0.06 ± 0.07 | 0.06 ± 0.07 | 0.06 ± 0.07 | 0.06 ± 0.07 | 0.06 ± 0.07 |
| | G | 0.06 ± 0.07 | 0.06 ± 0.07 | 0.06 ± 0.07 | 0.06 ± 0.07 | 0.06 ± 0.07 |
| | T | -0.08 ± 0.11 | -0.08 ± 0.11 | -0.08 ± 0.11 | -0.08 ± 0.11 | -0.08 ± 0.11 |
| MYC | A | -0.08 ± 0.11 | -0.08 ± 0.11 | -0.07 ± 0.1 | -0.08 ± 0.11 | -0.08 ± 0.11 |
| | C | 0.06 ± 0.07 | 0.06 ± 0.07 | 0.06 ± 0.07 | 0.06 ± 0.07 | 0.06 ± 0.07 |
| | G | 0.06 ± 0.07 | 0.06 ± 0.07 | 0.06 ± 0.07 | 0.06 ± 0.07 | 0.06 ± 0.07 |
| | T | -0.08 ± 0.1 | -0.08 ± 0.1 | -0.06 ± 0.1 | -0.08 ± 0.11 | -0.08 ± 0.11 |
| NANOG | A | -0.09 ± 0.11 | -0.08 ± 0.11 | -0.09 ± 0.11 | -0.09 ± 0.11 | -0.08 ± 0.11 |
| | C | 0.06 ± 0.07 | 0.06 ± 0.07 | 0.06 ± 0.07 | 0.07 ± 0.07 | 0.06 ± 0.07 |
| | G | 0.06 ± 0.07 | 0.06 ± 0.07 | 0.06 ± 0.07 | 0.06 ± 0.07 | 0.06 ± 0.07 |
| | T | -0.08 ± 0.11 | -0.08 ± 0.11 | -0.09 ± 0.11 | -0.08 ± 0.11 | -0.09 ± 0.11 |
| NFRKB | A | -0.08 ± 0.11 | -0.08 ± 0.11 | -0.08 ± 0.11 | -0.08 ± 0.11 | -0.08 ± 0.11 |
| | C | 0.06 ± 0.07 | 0.06 ± 0.07 | 0.06 ± 0.07 | 0.06 ± 0.07 | 0.06 ± 0.07 |
| | G | 0.06 ± 0.07 | 0.06 ± 0.07 | 0.06 ± 0.07 | 0.06 ± 0.07 | 0.06 ± 0.07 |
| | T | -0.08 ± 0.11 | -0.08 ± 0.11 | -0.08 ± 0.11 | -0.08 ± 0.11 | -0.08 ± 0.11 |
| RUNX1 | A | -0.08 ± 0.1 | -0.08 ± 0.1 | -0.07 ± 0.1 | -0.08 ± 0.11 | -0.08 ± 0.1 |
| | C | 0.06 ± 0.07 | 0.06 ± 0.07 | 0.06 ± 0.07 | 0.06 ± 0.07 | 0.06 ± 0.07 |
| | G | 0.06 ± 0.07 | 0.06 ± 0.07 | 0.06 ± 0.07 | 0.06 ± 0.07 | 0.06 ± 0.07 |
| | T | -0.08 ± 0.11 | -0.07 ± 0.11 | -0.07 ± 0.11 | -0.08 ± 0.11 | -0.08 ± 0.11 |
| SPI1 | A | -0.08 ± 0.11 | -0.08 ± 0.11 | -0.08 ± 0.11 | -0.08 ± 0.11 | -0.08 ± 0.11 |
| | C | 0.06 ± 0.07 | 0.06 ± 0.07 | 0.06 ± 0.07 | 0.06 ± 0.07 | 0.06 ± 0.07 |
| | G | 0.06 ± 0.07 | 0.06 ± 0.07 | 0.06 ± 0.07 | 0.06 ± 0.07 | 0.06 ± 0.07 |
| | T | -0.08 ± 0.11 | -0.08 ± 0.11 | -0.08 ± 0.11 | -0.08 ± 0.11 | -0.08 ± 0.11 |
| TP53 | A | -0.08 ± 0.11 | -0.08 ± 0.11 | -0.07 ± 0.1 | -0.08 ± 0.11 | -0.08 ± 0.1 |
| | C | 0.06 ± 0.07 | 0.06 ± 0.07 | 0.06 ± 0.07 | 0.06 ± 0.07 | 0.06 ± 0.07 |
| | G | 0.06 ± 0.07 | 0.06 ± 0.07 | 0.06 ± 0.07 | 0.06 ± 0.07 | 0.06 ± 0.07 |
| | T | -0.08 ± 0.11 | -0.08 ± 0.1 | -0.07 ± 0.1 | -0.08 ± 0.11 | -0.08 ± 0.11 |

**Table S10.** Average stagger values from selected positions relative to the binding site (prolonged peak center). The shape feature values were acquired using deepDNAshape from 50 bp windows.

**Table S11.** Average roll values from selected positions relative to the binding site (prolonged peak center). The shape feature values were acquired using deepDNAshape from 50 bp windows.

| | | -3000 bp | -1500 bp | at binding site | 1500 bp | 3000 bp |
|---|---|---|---|---|---|---|
| CTCF | AA | -3.31 ± 1.07 | -3.31 ± 1.07 | -3.24 ± 0.92 | -3.3 ± 1.04 | -3.32 ± 1.06 |
| | AC | -3.0 ± 0.94 | -3.01 ± 0.94 | -2.84 ± 0.75 | -3.0 ± 0.97 | -3.01 ± 0.96 |
| | AG | -2.39 ± 0.75 | -2.42 ± 0.77 | -2.22 ± 0.71 | -2.39 ± 0.77 | -2.39 ± 0.77 |
| | AT | -4.67 ± 1.17 | -4.67 ± 1.19 | -4.54 ± 1.01 | -4.69 ± 1.2 | -4.68 ± 1.22 |
| | CA | 4.06 ± 1.2 | 4.05 ± 1.19 | 3.72 ± 1.08 | 4.05 ± 1.2 | 4.07 ± 1.18 |
| | CC | -1.63 ± 0.7 | -1.63 ± 0.7 | -1.63 ± 0.69 | -1.62 ± 0.7 | -1.63 ± 0.7 |
| | CG | 3.42 ± 1.27 | 3.31 ± 1.24 | 3.0 ± 1.1 | 3.31 ± 1.26 | 3.4 ± 1.29 |
| | CT | -2.4 ± 0.77 | -2.39 ± 0.76 | -2.23 ± 0.71 | -2.4 ± 0.77 | -2.39 ± 0.76 |
| | GA | -1.11 ± 0.8 | -1.11 ± 0.77 | -1.15 ± 0.71 | -1.1 ± 0.78 | -1.11 ± 0.77 |
| | GC | -2.17 ± 0.9 | -2.19 ± 0.9 | -1.92 ± 0.74 | -2.16 ± 0.9 | -2.16 ± 0.88 |
| | GG | -1.64 ± 0.7 | -1.63 ± 0.69 | -1.64 ± 0.68 | -1.64 ± 0.7 | -1.62 ± 0.7 |
| | GT | -3.01 ± 0.98 | -3.01 ± 0.96 | -2.86 ± 0.78 | -3.01 ± 0.96 | -3.01 ± 0.97 |
| | TA | 5.39 ± 1.27 | 5.43 ± 1.28 | 5.09 ± 1.1 | 5.42 ± 1.28 | 5.44 ± 1.31 |
| | TC | -1.13 ± 0.79 | -1.12 ± 0.78 | -1.14 ± 0.71 | -1.09 ± 0.79 | -1.1 ± 0.78 |
| | TG | 4.07 ± 1.19 | 4.04 ± 1.2 | 3.69 ± 1.07 | 4.06 ± 1.19 | 4.07 ± 1.19 |
| | TT | -3.32 ± 1.06 | -3.31 ± 1.06 | -3.25 ± 0.94 | -3.33 ± 1.05 | -3.32 ± 1.07 |
| FOXK2 | AA | -3.31 ± 1.06 | -3.32 ± 1.08 | -3.14 ± 1.02 | -3.3 ± 1.04 | -3.34 ± 1.07 |
| | AC | -3.02 ± 0.97 | -3.0 ± 0.96 | -2.82 ± 0.86 | -3.0 ± 0.93 | -3.03 ± 0.95 |
| | AG | -2.39 ± 0.76 | -2.39 ± 0.76 | -2.38 ± 0.74 | -2.39 ± 0.79 | -2.4 ± 0.76 |
| | AT | -4.72 ± 1.21 | -4.7 ± 1.17 | -4.56 ± 1.02 | -4.71 ± 1.18 | -4.71 ± 1.21 |
| | CA | 4.07 ± 1.22 | 4.0 ± 1.2 | 4.13 ± 1.16 | 4.06 ± 1.19 | 4.08 ± 1.17 |
| | CC | -1.63 ± 0.68 | -1.61 ± 0.68 | -1.58 ± 0.7 | -1.6 ± 0.7 | -1.62 ± 0.68 |
| | CG | 3.38 ± 1.26 | 3.45 ± 1.28 | 3.2 ± 1.22 | 3.31 ± 1.28 | 3.34 ± 1.27 |
| | CT | -2.4 ± 0.78 | -2.4 ± 0.76 | -2.37 ± 0.72 | -2.39 ± 0.77 | -2.36 ± 0.75 |
| | GA | -1.12 ± 0.78 | -1.1 ± 0.79 | -1.11 ± 0.74 | -1.1 ± 0.77 | -1.11 ± 0.75 |
| | GC | -2.19 ± 0.89 | -2.16 ± 0.91 | -2.01 ± 0.84 | -2.17 ± 0.91 | -2.16 ± 0.86 |
| | GG | -1.6 ± 0.68 | -1.64 ± 0.7 | -1.56 ± 0.7 | -1.62 ± 0.7 | -1.61 ± 0.69 |
| | GT | -3.05 ± 0.98 | -3.03 ± 0.97 | -2.78 ± 0.84 | -2.99 ± 0.97 | -3.01 ± 0.95 |
| | TA | 5.44 ± 1.27 | 5.33 ± 1.29 | 5.35 ± 1.17 | 5.36 ± 1.26 | 5.42 ± 1.28 |
| | TC | -1.11 ± 0.81 | -1.07 ± 0.8 | -1.14 ± 0.73 | -1.11 ± 0.8 | -1.13 ± 0.78 |
| | TG | 4.03 ± 1.18 | 4.02 ± 1.2 | 4.11 ± 1.15 | 4.05 ± 1.2 | 4.04 ± 1.22 |
| | TT | -3.32 ± 1.07 | -3.35 ± 1.08 | -3.17 ± 1.04 | -3.32 ± 1.05 | -3.31 ± 1.08 |
| MEF2A | AA | -3.34 ± 1.05 | -3.31 ± 1.05 | -3.3 ± 1.05 | -3.31 ± 1.08 | -3.34 ± 1.03 |
| | AC | -3.0 ± 0.98 | -2.99 ± 0.94 | -3.02 ± 0.96 | -3.0 ± 0.97 | -3.0 ± 0.96 |
| | AG | -2.4 ± 0.78 | -2.38 ± 0.78 | -2.4 ± 0.77 | -2.37 ± 0.78 | -2.39 ± 0.79 |
| | AT | -4.72 ± 1.24 | -4.7 ± 1.19 | -4.79 ± 1.17 | -4.69 ± 1.21 | -4.72 ± 1.19 |
| | CA | 4.06 ± 1.22 | 4.06 ± 1.19 | 4.14 ± 1.15 | 4.1 ± 1.22 | 4.08 ± 1.19 |
| | CC | -1.61 ± 0.71 | -1.6 ± 0.71 | -1.6 ± 0.69 | -1.62 ± 0.7 | -1.63 ± 0.7 |
| | CG | 3.42 ± 1.34 | 3.34 ± 1.21 | 3.17 ± 1.22 | 3.43 ± 1.28 | 3.42 ± 1.24 |
| | CT | -2.41 ± 0.82 | -2.39 ± 0.78 | -2.37 ± 0.78 | -2.4 ± 0.75 | -2.41 ± 0.78 |
| | GA | -1.12 ± 0.81 | -1.11 ± 0.8 | -1.11 ± 0.76 | -1.07 ± 0.79 | -1.09 ± 0.8 |
| | GC | -2.16 ± 0.89 | -2.09 ± 0.89 | -2.08 ± 0.9 | -2.1 ± 0.87 | -2.15 ± 0.88 |
| | GG | -1.62 ± 0.7 | -1.59 ± 0.7 | -1.59 ± 0.71 | -1.62 ± 0.71 | -1.58 ± 0.69 |
| | GT | -3.01 ± 1.01 | -3.03 ± 0.98 | -3.0 ± 1.0 | -3.0 ± 0.95 | -3.01 ± 0.97 |
| | TA | 5.35 ± 1.27 | 5.4 ± 1.38 | 5.26 ± 1.29 | 5.48 ± 1.26 | 5.41 ± 1.26 |
| | TC | -1.1 ± 0.81 | -1.11 ± 0.79 | -1.1 ± 0.79 | -1.11 ± 0.81 | -1.11 ± 0.8 |
| | TG | 4.11 ± 1.2 | 4.11 ± 1.17 | 4.08 ± 1.16 | 4.11 ± 1.16 | 4.05 ± 1.2 |
| | TT | -3.32 ± 1.06 | -3.31 ± 1.07 | -3.26 ± 1.04 | -3.34 ± 1.07 | -3.29 ± 1.05 |
| MYC | AA | -3.33 ± 1.07 | -3.33 ± 1.06 | -3.25 ± 0.91 | -3.37 ± 1.04 | -3.33 ± 1.08 |
| | AC | -3.02 ± 0.95 | -3.03 ± 0.98 | -2.88 ± 0.78 | -3.03 ± 0.97 | -3.02 ± 0.93 |
| | AG | -2.39 ± 0.75 | -2.43 ± 0.75 | -2.35 ± 0.69 | -2.4 ± 0.78 | -2.4 ± 0.77 |

| | | | | | | |
|---|---|---|---|---|---|---|
| | AT | -4.71 ± 1.2 | -4.72 ± 1.22 | -4.64 ± 1.04 | -4.76 ± 1.22 | -4.77 ± 1.21 |
| | CA | 4.08 ± 1.21 | 4.0 ± 1.22 | 3.73 ± 1.08 | 4.0 ± 1.19 | 4.04 ± 1.22 |
| | CC | -1.63 ± 0.71 | -1.63 ± 0.69 | -1.58 ± 0.64 | -1.62 ± 0.7 | -1.64 ± 0.68 |
| | CG | 3.3 ± 1.22 | 3.24 ± 1.19 | 3.12 ± 1.2 | 3.32 ± 1.29 | 3.36 ± 1.18 |
| | CT | -2.4 ± 0.75 | -2.39 ± 0.76 | -2.36 ± 0.69 | -2.38 ± 0.73 | -2.41 ± 0.76 |
| | GA | -1.14 ± 0.81 | -1.12 ± 0.77 | -1.19 ± 0.67 | -1.14 ± 0.79 | -1.12 ± 0.8 |
| | GC | -2.16 ± 0.91 | -2.12 ± 0.86 | -1.87 ± 0.75 | -2.15 ± 0.89 | -2.15 ± 0.85 |
| | GG | -1.61 ± 0.68 | -1.63 ± 0.69 | -1.58 ± 0.66 | -1.61 ± 0.71 | -1.6 ± 0.7 |
| | GT | -3.01 ± 0.95 | -3.02 ± 0.93 | -2.85 ± 0.78 | -2.98 ± 0.92 | -3.01 ± 0.96 |
| | TA | 5.37 ± 1.25 | 5.36 ± 1.27 | 5.13 ± 1.16 | 5.43 ± 1.24 | 5.36 ± 1.27 |
| | TC | -1.09 ± 0.76 | -1.13 ± 0.76 | -1.21 ± 0.68 | -1.15 ± 0.77 | -1.09 ± 0.77 |
| | TG | 3.99 ± 1.19 | 3.98 ± 1.18 | 3.72 ± 1.09 | 4.0 ± 1.19 | 4.04 ± 1.2 |
| | TT | -3.32 ± 1.04 | -3.34 ± 1.07 | -3.28 ± 0.94 | -3.36 ± 1.05 | -3.32 ± 1.04 |
| NANOG | AA | -3.31 ± 1.07 | -3.29 ± 1.07 | -3.13 ± 1.01 | -3.27 ± 1.05 | -3.24 ± 1.07 |
| | AC | -3.03 ± 0.96 | -3.08 ± 1.01 | -2.95 ± 0.93 | -3.01 ± 0.99 | -3.01 ± 1.0 |
| | AG | -2.41 ± 0.78 | -2.41 ± 0.82 | -2.43 ± 0.76 | -2.39 ± 0.76 | -2.45 ± 0.77 |
| | AT | -4.63 ± 1.2 | -4.66 ± 1.23 | -4.75 ± 1.06 | -4.68 ± 1.23 | -4.69 ± 1.26 |
| | CA | 4.18 ± 1.21 | 4.19 ± 1.19 | 4.2 ± 1.19 | 4.12 ± 1.19 | 4.15 ± 1.17 |
| | CC | -1.63 ± 0.73 | -1.63 ± 0.71 | -1.69 ± 0.72 | -1.59 ± 0.69 | -1.57 ± 0.71 |
| | CG | 3.5 ± 1.15 | 3.52 ± 1.25 | 3.16 ± 1.15 | 3.4 ± 1.28 | 3.31 ± 1.26 |
| | CT | -2.38 ± 0.76 | -2.41 ± 0.82 | -2.39 ± 0.77 | -2.4 ± 0.79 | -2.4 ± 0.78 |
| | GA | -1.1 ± 0.8 | -1.11 ± 0.81 | -1.07 ± 0.83 | -1.09 ± 0.82 | -1.08 ± 0.83 |
| | GC | -2.22 ± 0.97 | -2.2 ± 0.91 | -2.19 ± 0.97 | -2.21 ± 0.94 | -2.17 ± 0.93 |
| | GG | -1.64 ± 0.7 | -1.63 ± 0.7 | -1.66 ± 0.71 | -1.63 ± 0.7 | -1.64 ± 0.72 |
| | GT | -3.02 ± 0.95 | -3.03 ± 1.0 | -2.97 ± 0.95 | -3.03 ± 1.01 | -3.06 ± 1.03 |
| | TA | 5.54 ± 1.32 | 5.41 ± 1.31 | 5.5 ± 1.25 | 5.42 ± 1.29 | 5.44 ± 1.28 |
| | TC | -1.08 ± 0.8 | -1.11 ± 0.83 | -1.05 ± 0.78 | -1.07 ± 0.78 | -1.06 ± 0.81 |
| | TG | 4.27 ± 1.19 | 4.17 ± 1.19 | 4.17 ± 1.25 | 4.17 ± 1.23 | 4.14 ± 1.17 |
| | TT | -3.3 ± 1.07 | -3.32 ± 1.06 | -3.1 ± 0.97 | -3.31 ± 1.04 | -3.29 ± 1.06 |
| NFRKB | AA | -3.34 ± 1.07 | -3.36 ± 1.07 | -3.33 ± 1.02 | -3.34 ± 1.06 | -3.36 ± 1.06 |
| | AC | -3.01 ± 0.95 | -2.99 ± 0.99 | -2.95 ± 0.92 | -3.04 ± 0.97 | -3.04 ± 1.0 |
| | AG | -2.41 ± 0.77 | -2.41 ± 0.77 | -2.38 ± 0.68 | -2.38 ± 0.77 | -2.4 ± 0.75 |
| | AT | -4.7 ± 1.22 | -4.69 ± 1.19 | -4.67 ± 1.1 | -4.68 ± 1.18 | -4.74 ± 1.22 |
| | CA | 4.06 ± 1.22 | 4.13 ± 1.21 | 4.0 ± 1.18 | 4.06 ± 1.18 | 4.08 ± 1.22 |
| | CC | -1.6 ± 0.68 | -1.58 ± 0.71 | -1.58 ± 0.7 | -1.63 ± 0.71 | -1.61 ± 0.68 |
| | CG | 3.36 ± 1.25 | 3.44 ± 1.28 | 3.19 ± 1.17 | 3.25 ± 1.28 | 3.36 ± 1.3 |
| | CT | -2.4 ± 0.8 | -2.39 ± 0.78 | -2.38 ± 0.72 | -2.39 ± 0.75 | -2.4 ± 0.77 |
| | GA | -1.12 ± 0.77 | -1.08 ± 0.77 | -1.12 ± 0.74 | -1.12 ± 0.8 | -1.08 ± 0.78 |
| | GC | -2.2 ± 0.88 | -2.13 ± 0.88 | -2.04 ± 0.83 | -2.16 ± 0.89 | -2.18 ± 0.89 |
| | GG | -1.65 ± 0.68 | -1.65 ± 0.68 | -1.58 ± 0.71 | -1.6 ± 0.69 | -1.61 ± 0.71 |
| | GT | -3.02 ± 0.98 | -3.03 ± 0.98 | -2.97 ± 0.95 | -3.05 ± 1.03 | -3.02 ± 0.98 |
| | TA | 5.41 ± 1.24 | 5.36 ± 1.32 | 5.25 ± 1.31 | 5.42 ± 1.22 | 5.35 ± 1.25 |
| | TC | -1.11 ± 0.74 | -1.08 ± 0.82 | -1.16 ± 0.76 | -1.13 ± 0.78 | -1.11 ± 0.82 |
| | TG | 4.07 ± 1.2 | 4.0 ± 1.22 | 4.0 ± 1.11 | 4.07 ± 1.18 | 4.09 ± 1.18 |
| | TT | -3.32 ± 1.07 | -3.34 ± 1.04 | -3.29 ± 1.03 | -3.32 ± 1.06 | -3.33 ± 1.08 |
| RUNX1 | AA | -3.36 ± 1.11 | -3.28 ± 1.02 | -3.12 ± 0.92 | -3.25 ± 1.01 | -3.33 ± 1.11 |
| | AC | -3.01 ± 0.91 | -2.88 ± 0.9 | -2.9 ± 0.88 | -3.01 ± 0.92 | -2.94 ± 0.92 |
| | AG | -2.33 ± 0.79 | -2.42 ± 0.82 | -2.37 ± 0.65 | -2.35 ± 0.79 | -2.43 ± 0.74 |
| | AT | -4.81 ± 1.17 | -4.69 ± 1.14 | -4.58 ± 1.03 | -4.68 ± 1.19 | -4.62 ± 1.18 |
| | CA | 3.88 ± 1.2 | 3.93 ± 1.14 | 3.88 ± 1.18 | 3.9 ± 1.2 | 4.18 ± 1.25 |
| | CC | -1.55 ± 0.68 | -1.68 ± 0.69 | -1.53 ± 0.68 | -1.59 ± 0.72 | -1.68 ± 0.69 |
| | CG | 3.36 ± 1.22 | 3.28 ± 1.07 | 2.91 ± 1.12 | 3.16 ± 1.29 | 3.11 ± 1.28 |
| | CT | -2.42 ± 0.8 | -2.39 ± 0.84 | -2.35 ± 0.71 | -2.37 ± 0.7 | -2.41 ± 0.74 |
| | GA | -1.17 ± 0.76 | -1.13 ± 0.77 | -1.17 ± 0.68 | -1.12 ± 0.73 | -1.06 ± 0.77 |
| | GC | -2.17 ± 0.84 | -2.08 ± 0.77 | -1.89 ± 0.76 | -2.12 ± 0.84 | -2.09 ± 0.9 |
| | GG | -1.65 ± 0.72 | -1.63 ± 0.73 | -1.54 ± 0.7 | -1.53 ± 0.69 | -1.6 ± 0.68 |

| | | | | | | |
|---|---|---|---|---|---|---|
| | GT | -3.03 ± 0.98 | -3.05 ± 0.92 | -2.96 ± 0.89 | -3.0 ± 0.96 | -2.96 ± 0.91 |
| | TA | 5.32 ± 1.49 | 5.41 ± 1.22 | 5.25 ± 1.24 | 5.38 ± 1.25 | 5.25 ± 1.32 |
| | TC | -1.09 ± 0.81 | -1.12 ± 0.78 | -1.1 ± 0.68 | -1.05 ± 0.68 | -1.1 ± 0.76 |
| | TG | 3.97 ± 1.22 | 4.01 ± 1.1 | 3.89 ± 1.16 | 3.95 ± 1.14 | 4.16 ± 1.2 |
| | TT | -3.35 ± 1.1 | -3.37 ± 1.03 | -3.16 ± 0.94 | -3.32 ± 1.12 | -3.28 ± 1.03 |
| SPI1 | AA | -3.33 ± 1.07 | -3.29 ± 1.06 | -3.41 ± 0.95 | -3.29 ± 1.04 | -3.35 ± 1.06 |
| | AC | -3.06 ± 0.97 | -3.02 ± 0.94 | -3.14 ± 0.91 | -3.0 ± 0.94 | -3.03 ± 0.95 |
| | AG | -2.39 ± 0.76 | -2.4 ± 0.79 | -2.43 ± 0.72 | -2.39 ± 0.79 | -2.38 ± 0.79 |
| | AT | -4.71 ± 1.19 | -4.68 ± 1.18 | -4.77 ± 1.08 | -4.69 ± 1.21 | -4.72 ± 1.22 |
| | CA | 4.07 ± 1.2 | 4.09 ± 1.2 | 3.91 ± 1.19 | 4.07 ± 1.17 | 4.1 ± 1.2 |
| | CC | -1.65 ± 0.71 | -1.62 ± 0.7 | -1.65 ± 0.7 | -1.62 ± 0.69 | -1.64 ± 0.72 |
| | CG | 3.3 ± 1.25 | 3.3 ± 1.28 | 3.09 ± 1.2 | 3.33 ± 1.28 | 3.38 ± 1.29 |
| | CT | -2.37 ± 0.76 | -2.39 ± 0.76 | -2.42 ± 0.71 | -2.42 ± 0.77 | -2.39 ± 0.75 |
| | GA | -1.09 ± 0.79 | -1.11 ± 0.79 | -1.06 ± 0.68 | -1.15 ± 0.79 | -1.12 ± 0.78 |
| | GC | -2.16 ± 0.87 | -2.18 ± 0.89 | -2.09 ± 0.9 | -2.19 ± 0.88 | -2.16 ± 0.86 |
| | GG | -1.6 ± 0.71 | -1.65 ± 0.69 | -1.65 ± 0.71 | -1.59 ± 0.69 | -1.63 ± 0.7 |
| | GT | -3.02 ± 0.96 | -3.02 ± 1.0 | -3.12 ± 0.93 | -3.0 ± 0.97 | -3.03 ± 0.98 |
| | TA | 5.4 ± 1.3 | 5.39 ± 1.34 | 5.23 ± 1.21 | 5.44 ± 1.23 | 5.42 ± 1.27 |
| | TC | -1.12 ± 0.8 | -1.1 ± 0.77 | -1.11 ± 0.67 | -1.09 ± 0.79 | -1.11 ± 0.78 |
| | TG | 4.08 ± 1.19 | 4.07 ± 1.19 | 3.89 ± 1.21 | 4.05 ± 1.19 | 4.09 ± 1.2 |
| | TT | -3.33 ± 1.08 | -3.32 ± 1.06 | -3.38 ± 0.94 | -3.31 ± 1.09 | -3.29 ± 1.06 |
| TP53 | AA | -3.33 ± 1.09 | -3.36 ± 1.03 | -3.27 ± 0.93 | -3.3 ± 1.07 | -3.34 ± 1.04 |
| | AC | -2.98 ± 0.93 | -3.02 ± 0.99 | -2.94 ± 0.81 | -2.96 ± 0.91 | -3.01 ± 0.96 |
| | AG | -2.4 ± 0.75 | -2.4 ± 0.73 | -2.38 ± 0.68 | -2.42 ± 0.75 | -2.4 ± 0.76 |
| | AT | -4.74 ± 1.22 | -4.71 ± 1.19 | -4.7 ± 1.07 | -4.76 ± 1.17 | -4.76 ± 1.2 |
| | CA | 4.03 ± 1.2 | 4.03 ± 1.2 | 3.75 ± 1.11 | 3.95 ± 1.21 | 4.02 ± 1.18 |
| | CC | -1.62 ± 0.68 | -1.66 ± 0.7 | -1.61 ± 0.63 | -1.63 ± 0.7 | -1.64 ± 0.68 |
| | CG | 3.23 ± 1.21 | 3.28 ± 1.23 | 2.99 ± 1.16 | 3.31 ± 1.22 | 3.2 ± 1.27 |
| | CT | -2.38 ± 0.77 | -2.39 ± 0.73 | -2.38 ± 0.7 | -2.4 ± 0.77 | -2.39 ± 0.74 |
| | GA | -1.1 ± 0.75 | -1.12 ± 0.76 | -1.16 ± 0.71 | -1.12 ± 0.78 | -1.14 ± 0.8 |
| | GC | -2.16 ± 0.91 | -2.13 ± 0.89 | -1.92 ± 0.77 | -2.17 ± 0.87 | -2.14 ± 0.94 |
| | GG | -1.63 ± 0.7 | -1.62 ± 0.7 | -1.59 ± 0.64 | -1.62 ± 0.68 | -1.58 ± 0.68 |
| | GT | -2.99 ± 0.92 | -3.0 ± 0.93 | -2.9 ± 0.81 | -3.0 ± 0.94 | -3.01 ± 0.98 |
| | TA | 5.4 ± 1.29 | 5.29 ± 1.25 | 5.14 ± 1.22 | 5.37 ± 1.23 | 5.38 ± 1.3 |
| | TC | -1.15 ± 0.78 | -1.14 ± 0.75 | -1.18 ± 0.7 | -1.15 ± 0.77 | -1.15 ± 0.77 |
| | TG | 4.01 ± 1.18 | 4.03 ± 1.19 | 3.74 ± 1.1 | 4.01 ± 1.2 | 4.04 ± 1.17 |
| | TT | -3.33 ± 1.02 | -3.38 ± 1.06 | -3.28 ± 0.93 | -3.32 ± 1.08 | -3.38 ± 1.07 |

**Table S12.** Average helical twist values from selected positions relative to the binding site (prolonged peak center). The shape feature values were acquired using deepDNAshape from 50 bp windows.

| | | -3000 bp | -1500 bp | at binding site | 1500 bp | 3000 bp |
|---|---|---|---|---|---|---|
| CTCF | AA | 35.85 ± 0.9 | 35.84 ± 0.9 | 35.57 ± 0.85 | 35.85 ± 0.89 | 35.86 ± 0.91 |
| | AC | 34.51 ± 0.77 | 34.53 ± 0.78 | 34.32 ± 0.64 | 34.52 ± 0.79 | 34.52 ± 0.78 |
| | AG | 31.84 ± 0.66 | 31.85 ± 0.66 | 31.63 ± 0.51 | 31.84 ± 0.66 | 31.84 ± 0.65 |
| | AT | 32.67 ± 0.8 | 32.68 ± 0.82 | 32.43 ± 0.71 | 32.68 ± 0.81 | 32.69 ± 0.82 |
| | CA | 34.62 ± 0.57 | 34.62 ± 0.56 | 34.44 ± 0.52 | 34.62 ± 0.57 | 34.62 ± 0.56 |
| | CC | 33.9 ± 0.59 | 33.88 ± 0.58 | 33.81 ± 0.53 | 33.9 ± 0.58 | 33.91 ± 0.6 |
| | CG | 32.99 ± 0.6 | 32.96 ± 0.6 | 32.74 ± 0.55 | 32.94 ± 0.58 | 32.96 ± 0.6 |
| | CT | 31.83 ± 0.65 | 31.83 ± 0.65 | 31.64 ± 0.52 | 31.84 ± 0.64 | 31.84 ± 0.64 |
| | GA | 35.78 ± 0.68 | 35.78 ± 0.66 | 35.73 ± 0.58 | 35.77 ± 0.67 | 35.78 ± 0.67 |
| | GC | 36.67 ± 0.62 | 36.69 ± 0.61 | 36.57 ± 0.53 | 36.67 ± 0.62 | 36.68 ± 0.62 |
| | GG | 33.91 ± 0.59 | 33.89 ± 0.58 | 33.81 ± 0.52 | 33.89 ± 0.58 | 33.89 ± 0.58 |
| | GT | 34.52 ± 0.79 | 34.52 ± 0.79 | 34.33 ± 0.66 | 34.53 ± 0.79 | 34.52 ± 0.78 |
| | TA | 34.66 ± 0.46 | 34.65 ± 0.46 | 34.52 ± 0.4 | 34.63 ± 0.45 | 34.64 ± 0.45 |
| | TC | 35.79 ± 0.69 | 35.77 ± 0.67 | 35.71 ± 0.59 | 35.76 ± 0.67 | 35.78 ± 0.67 |
| | TG | 34.61 ± 0.56 | 34.61 ± 0.55 | 34.43 ± 0.51 | 34.62 ± 0.56 | 34.62 ± 0.56 |
| | TT | 35.86 ± 0.9 | 35.85 ± 0.89 | 35.58 ± 0.86 | 35.86 ± 0.9 | 35.86 ± 0.9 |
| FOXK2 | AA | 35.91 ± 0.89 | 35.88 ± 0.92 | 35.71 ± 0.83 | 35.87 ± 0.91 | 35.87 ± 0.9 |
| | AC | 34.53 ± 0.8 | 34.53 ± 0.79 | 34.42 ± 0.71 | 34.51 ± 0.76 | 34.53 ± 0.78 |
| | AG | 31.83 ± 0.65 | 31.84 ± 0.65 | 31.81 ± 0.61 | 31.84 ± 0.65 | 31.83 ± 0.66 |
| | AT | 32.69 ± 0.81 | 32.68 ± 0.81 | 32.49 ± 0.69 | 32.68 ± 0.81 | 32.7 ± 0.82 |
| | CA | 34.6 ± 0.56 | 34.62 ± 0.56 | 34.67 ± 0.57 | 34.62 ± 0.56 | 34.61 ± 0.55 |
| | CC | 33.89 ± 0.58 | 33.88 ± 0.57 | 33.84 ± 0.53 | 33.88 ± 0.57 | 33.91 ± 0.59 |
| | CG | 33.0 ± 0.6 | 32.97 ± 0.59 | 32.91 ± 0.57 | 32.92 ± 0.58 | 32.97 ± 0.59 |
| | CT | 31.83 ± 0.65 | 31.84 ± 0.64 | 31.8 ± 0.6 | 31.84 ± 0.65 | 31.84 ± 0.65 |
| | GA | 35.76 ± 0.68 | 35.78 ± 0.68 | 35.74 ± 0.62 | 35.78 ± 0.67 | 35.77 ± 0.67 |
| | GC | 36.68 ± 0.62 | 36.67 ± 0.63 | 36.56 ± 0.62 | 36.67 ± 0.62 | 36.67 ± 0.61 |
| | GG | 33.9 ± 0.58 | 33.89 ± 0.57 | 33.84 ± 0.54 | 33.9 ± 0.58 | 33.91 ± 0.59 |
| | GT | 34.53 ± 0.78 | 34.53 ± 0.8 | 34.4 ± 0.69 | 34.51 ± 0.77 | 34.52 ± 0.78 |
| | TA | 34.62 ± 0.45 | 34.65 ± 0.46 | 34.58 ± 0.42 | 34.65 ± 0.46 | 34.64 ± 0.45 |
| | TC | 35.79 ± 0.69 | 35.74 ± 0.67 | 35.75 ± 0.62 | 35.78 ± 0.69 | 35.79 ± 0.68 |
| | TG | 34.61 ± 0.55 | 34.61 ± 0.57 | 34.67 ± 0.56 | 34.6 ± 0.55 | 34.6 ± 0.55 |
| | TT | 35.87 ± 0.91 | 35.89 ± 0.9 | 35.71 ± 0.83 | 35.89 ± 0.92 | 35.87 ± 0.91 |
| MEF2A | AA | 35.85 ± 0.9 | 35.92 ± 0.89 | 35.84 ± 0.85 | 35.89 ± 0.91 | 35.89 ± 0.9 |
| | AC | 34.5 ± 0.78 | 34.5 ± 0.78 | 34.55 ± 0.8 | 34.54 ± 0.81 | 34.54 ± 0.8 |
| | AG | 31.87 ± 0.65 | 31.85 ± 0.64 | 31.85 ± 0.65 | 31.86 ± 0.65 | 31.88 ± 0.67 |
| | AT | 32.71 ± 0.83 | 32.69 ± 0.82 | 32.71 ± 0.83 | 32.68 ± 0.84 | 32.7 ± 0.82 |
| | CA | 34.64 ± 0.57 | 34.61 ± 0.56 | 34.65 ± 0.56 | 34.62 ± 0.57 | 34.63 ± 0.55 |
| | CC | 33.92 ± 0.6 | 33.9 ± 0.56 | 33.88 ± 0.59 | 33.89 ± 0.59 | 33.91 ± 0.58 |
| | CG | 32.98 ± 0.61 | 32.92 ± 0.55 | 32.87 ± 0.56 | 32.99 ± 0.63 | 33.01 ± 0.6 |
| | CT | 31.87 ± 0.67 | 31.85 ± 0.65 | 31.84 ± 0.64 | 31.89 ± 0.66 | 31.86 ± 0.67 |
| | GA | 35.82 ± 0.69 | 35.8 ± 0.69 | 35.74 ± 0.67 | 35.77 ± 0.69 | 35.8 ± 0.7 |
| | GC | 36.69 ± 0.62 | 36.63 ± 0.63 | 36.62 ± 0.65 | 36.63 ± 0.62 | 36.67 ± 0.62 |
| | GG | 33.91 ± 0.61 | 33.9 ± 0.6 | 33.91 ± 0.59 | 33.91 ± 0.6 | 33.91 ± 0.6 |
| | GT | 34.54 ± 0.83 | 34.55 ± 0.78 | 34.56 ± 0.83 | 34.56 ± 0.79 | 34.54 ± 0.8 |
| | TA | 34.65 ± 0.45 | 34.64 ± 0.47 | 34.67 ± 0.48 | 34.62 ± 0.46 | 34.65 ± 0.46 |
| | TC | 35.78 ± 0.68 | 35.79 ± 0.68 | 35.77 ± 0.67 | 35.78 ± 0.7 | 35.79 ± 0.7 |
| | TG | 34.64 ± 0.56 | 34.62 ± 0.56 | 34.62 ± 0.53 | 34.63 ± 0.56 | 34.62 ± 0.56 |
| | TT | 35.89 ± 0.9 | 35.88 ± 0.92 | 35.82 ± 0.87 | 35.88 ± 0.9 | 35.87 ± 0.88 |
| MYC | AA | 35.87 ± 0.93 | 35.85 ± 0.9 | 35.54 ± 0.88 | 35.85 ± 0.91 | 35.86 ± 0.91 |
| | AC | 34.52 ± 0.79 | 34.51 ± 0.79 | 34.32 ± 0.67 | 34.54 ± 0.79 | 34.52 ± 0.77 |
| | AG | 31.83 ± 0.64 | 31.82 ± 0.64 | 31.69 ± 0.55 | 31.85 ± 0.66 | 31.83 ± 0.65 |

| | | | | | | |
|---|---|---|---|---|---|---|
| | AT | 32.71 ± 0.83 | 32.69 ± 0.84 | 32.47 ± 0.73 | 32.72 ± 0.84 | 32.7 ± 0.82 |
| | CA | 34.61 ± 0.55 | 34.6 ± 0.55 | 34.48 ± 0.51 | 34.59 ± 0.54 | 34.6 ± 0.56 |
| | CC | 33.87 ± 0.58 | 33.89 ± 0.59 | 33.75 ± 0.47 | 33.86 ± 0.55 | 33.87 ± 0.57 |
| | CG | 32.95 ± 0.58 | 32.89 ± 0.55 | 32.86 ± 0.57 | 32.94 ± 0.61 | 32.97 ± 0.59 |
| | CT | 31.81 ± 0.63 | 31.82 ± 0.63 | 31.63 ± 0.51 | 31.82 ± 0.64 | 31.82 ± 0.66 |
| | GA | 35.74 ± 0.68 | 35.76 ± 0.67 | 35.66 ± 0.58 | 35.76 ± 0.67 | 35.76 ± 0.69 |
| | GC | 36.68 ± 0.62 | 36.64 ± 0.6 | 36.48 ± 0.58 | 36.66 ± 0.62 | 36.67 ± 0.6 |
| | GG | 33.9 ± 0.58 | 33.87 ± 0.57 | 33.76 ± 0.48 | 33.85 ± 0.56 | 33.89 ± 0.59 |
| | GT | 34.49 ± 0.78 | 34.52 ± 0.77 | 34.3 ± 0.67 | 34.51 ± 0.75 | 34.52 ± 0.79 |
| | TA | 34.63 ± 0.45 | 34.62 ± 0.44 | 34.54 ± 0.4 | 34.62 ± 0.44 | 34.64 ± 0.45 |
| | TC | 35.76 ± 0.68 | 35.77 ± 0.67 | 35.68 ± 0.58 | 35.76 ± 0.66 | 35.74 ± 0.66 |
| | TG | 34.57 ± 0.55 | 34.58 ± 0.56 | 34.48 ± 0.51 | 34.59 ± 0.55 | 34.58 ± 0.55 |
| | TT | 35.83 ± 0.89 | 35.85 ± 0.92 | 35.57 ± 0.87 | 35.89 ± 0.94 | 35.85 ± 0.91 |
| NANOG | AA | 35.91 ± 0.91 | 35.89 ± 0.87 | 35.71 ± 0.76 | 35.86 ± 0.86 | 35.87 ± 0.85 |
| | AC | 34.56 ± 0.76 | 34.61 ± 0.83 | 34.47 ± 0.72 | 34.55 ± 0.81 | 34.55 ± 0.78 |
| | AG | 31.89 ± 0.67 | 31.89 ± 0.67 | 31.87 ± 0.67 | 31.86 ± 0.64 | 31.9 ± 0.65 |
| | AT | 32.69 ± 0.82 | 32.69 ± 0.8 | 32.64 ± 0.74 | 32.7 ± 0.82 | 32.72 ± 0.87 |
| | CA | 34.66 ± 0.56 | 34.68 ± 0.58 | 34.67 ± 0.55 | 34.66 ± 0.56 | 34.66 ± 0.56 |
| | CC | 33.92 ± 0.6 | 33.9 ± 0.62 | 33.86 ± 0.57 | 33.94 ± 0.6 | 33.91 ± 0.6 |
| | CG | 33.02 ± 0.61 | 33.06 ± 0.6 | 32.84 ± 0.48 | 33.02 ± 0.64 | 32.96 ± 0.58 |
| | CT | 31.88 ± 0.68 | 31.88 ± 0.69 | 31.88 ± 0.67 | 31.87 ± 0.65 | 31.92 ± 0.69 |
| | GA | 35.79 ± 0.7 | 35.84 ± 0.69 | 35.82 ± 0.67 | 35.79 ± 0.69 | 35.77 ± 0.72 |
| | GC | 36.69 ± 0.65 | 36.67 ± 0.64 | 36.64 ± 0.66 | 36.71 ± 0.64 | 36.65 ± 0.63 |
| | GG | 33.9 ± 0.59 | 33.93 ± 0.62 | 33.87 ± 0.6 | 33.96 ± 0.59 | 33.93 ± 0.63 |
| | GT | 34.57 ± 0.8 | 34.57 ± 0.77 | 34.5 ± 0.78 | 34.56 ± 0.81 | 34.58 ± 0.82 |
| | TA | 34.65 ± 0.45 | 34.65 ± 0.47 | 34.58 ± 0.45 | 34.66 ± 0.46 | 34.67 ± 0.47 |
| | TC | 35.78 ± 0.71 | 35.81 ± 0.69 | 35.78 ± 0.67 | 35.79 ± 0.69 | 35.79 ± 0.72 |
| | TG | 34.7 ± 0.57 | 34.64 ± 0.55 | 34.66 ± 0.56 | 34.69 ± 0.6 | 34.66 ± 0.57 |
| | TT | 35.89 ± 0.91 | 35.89 ± 0.89 | 35.73 ± 0.78 | 35.89 ± 0.87 | 35.9 ± 0.88 |
| NFRKB | AA | 35.89 ± 0.91 | 35.9 ± 0.92 | 35.77 ± 0.9 | 35.87 ± 0.88 | 35.9 ± 0.88 |
| | AC | 34.55 ± 0.79 | 34.52 ± 0.78 | 34.46 ± 0.72 | 34.53 ± 0.8 | 34.51 ± 0.79 |
| | AG | 31.86 ± 0.67 | 31.86 ± 0.66 | 31.79 ± 0.59 | 31.84 ± 0.67 | 31.86 ± 0.66 |
| | AT | 32.68 ± 0.81 | 32.69 ± 0.81 | 32.61 ± 0.78 | 32.69 ± 0.78 | 32.71 ± 0.81 |
| | CA | 34.62 ± 0.57 | 34.64 ± 0.58 | 34.58 ± 0.55 | 34.61 ± 0.55 | 34.6 ± 0.55 |
| | CC | 33.93 ± 0.6 | 33.89 ± 0.6 | 33.82 ± 0.52 | 33.93 ± 0.58 | 33.91 ± 0.6 |
| | CG | 32.95 ± 0.61 | 32.95 ± 0.58 | 32.86 ± 0.53 | 32.92 ± 0.6 | 32.95 ± 0.58 |
| | CT | 31.86 ± 0.66 | 31.85 ± 0.64 | 31.79 ± 0.62 | 31.85 ± 0.63 | 31.84 ± 0.67 |
| | GA | 35.81 ± 0.68 | 35.83 ± 0.67 | 35.75 ± 0.64 | 35.76 ± 0.67 | 35.76 ± 0.69 |
| | GC | 36.71 ± 0.61 | 36.64 ± 0.63 | 36.58 ± 0.6 | 36.67 ± 0.62 | 36.7 ± 0.61 |
| | GG | 33.91 ± 0.6 | 33.86 ± 0.59 | 33.84 ± 0.54 | 33.88 ± 0.58 | 33.9 ± 0.6 |
| | GT | 34.53 ± 0.78 | 34.51 ± 0.79 | 34.48 ± 0.79 | 34.54 ± 0.81 | 34.55 ± 0.81 |
| | TA | 34.65 ± 0.48 | 34.64 ± 0.45 | 34.6 ± 0.43 | 34.63 ± 0.45 | 34.65 ± 0.45 |
| | TC | 35.79 ± 0.69 | 35.75 ± 0.69 | 35.79 ± 0.66 | 35.79 ± 0.68 | 35.79 ± 0.7 |
| | TG | 34.6 ± 0.56 | 34.58 ± 0.56 | 34.6 ± 0.55 | 34.62 ± 0.56 | 34.66 ± 0.56 |
| | TT | 35.87 ± 0.92 | 35.87 ± 0.89 | 35.73 ± 0.9 | 35.88 ± 0.9 | 35.92 ± 0.91 |
| RUNX1 | AA | 35.92 ± 0.95 | 35.85 ± 0.92 | 35.63 ± 0.91 | 35.85 ± 0.85 | 35.9 ± 0.92 |
| | AC | 34.47 ± 0.74 | 34.43 ± 0.74 | 34.4 ± 0.84 | 34.46 ± 0.77 | 34.49 ± 0.75 |
| | AG | 31.8 ± 0.65 | 31.87 ± 0.72 | 31.61 ± 0.52 | 31.81 ± 0.63 | 31.79 ± 0.62 |
| | AT | 32.69 ± 0.76 | 32.65 ± 0.8 | 32.52 ± 0.66 | 32.64 ± 0.79 | 32.65 ± 0.78 |
| | CA | 34.54 ± 0.55 | 34.61 ± 0.55 | 34.55 ± 0.57 | 34.56 ± 0.54 | 34.61 ± 0.57 |
| | CC | 33.9 ± 0.58 | 33.88 ± 0.59 | 33.85 ± 0.54 | 33.94 ± 0.58 | 33.88 ± 0.57 |
| | CG | 32.93 ± 0.64 | 32.85 ± 0.52 | 32.78 ± 0.44 | 32.9 ± 0.56 | 32.8 ± 0.6 |
| | CT | 31.86 ± 0.69 | 31.87 ± 0.67 | 31.74 ± 0.64 | 31.77 ± 0.64 | 31.82 ± 0.66 |
| | GA | 35.74 ± 0.62 | 35.8 ± 0.6 | 35.67 ± 0.64 | 35.81 ± 0.73 | 35.71 ± 0.67 |
| | GC | 36.69 ± 0.57 | 36.64 ± 0.58 | 36.44 ± 0.58 | 36.69 ± 0.6 | 36.6 ± 0.61 |
| | GG | 33.88 ± 0.57 | 33.87 ± 0.57 | 33.82 ± 0.55 | 33.83 ± 0.5 | 33.91 ± 0.58 |

| | | | | | | |
|---|---|---|---|---|---|---|
| | GT | 34.52 ± 0.81 | 34.52 ± 0.79 | 34.46 ± 0.87 | 34.5 ± 0.76 | 34.48 ± 0.77 |
| | TA | 34.64 ± 0.5 | 34.58 ± 0.42 | 34.58 ± 0.36 | 34.61 ± 0.44 | 34.64 ± 0.48 |
| | TC | 35.73 ± 0.7 | 35.71 ± 0.65 | 35.72 ± 0.59 | 35.75 ± 0.66 | 35.78 ± 0.64 |
| | TG | 34.58 ± 0.53 | 34.57 ± 0.54 | 34.59 ± 0.47 | 34.56 ± 0.51 | 34.62 ± 0.57 |
| | TT | 35.89 ± 0.96 | 35.83 ± 0.92 | 35.55 ± 0.71 | 35.86 ± 0.92 | 35.81 ± 0.89 |
| SPI1 | AA | 35.88 ± 0.92 | 35.86 ± 0.89 | 35.68 ± 0.83 | 35.86 ± 0.87 | 35.88 ± 0.89 |
| | AC | 34.56 ± 0.79 | 34.55 ± 0.79 | 34.58 ± 0.78 | 34.51 ± 0.76 | 34.54 ± 0.77 |
| | AG | 31.85 ± 0.63 | 31.85 ± 0.67 | 31.89 ± 0.63 | 31.84 ± 0.65 | 31.86 ± 0.66 |
| | AT | 32.69 ± 0.82 | 32.68 ± 0.81 | 32.63 ± 0.77 | 32.69 ± 0.83 | 32.72 ± 0.83 |
| | CA | 34.6 ± 0.56 | 34.63 ± 0.56 | 34.51 ± 0.52 | 34.61 ± 0.55 | 34.64 ± 0.57 |
| | CC | 33.89 ± 0.6 | 33.9 ± 0.59 | 33.9 ± 0.55 | 33.92 ± 0.59 | 33.92 ± 0.58 |
| | CG | 32.95 ± 0.59 | 32.89 ± 0.56 | 32.85 ± 0.56 | 32.97 ± 0.59 | 32.98 ± 0.6 |
| | CT | 31.84 ± 0.64 | 31.84 ± 0.63 | 31.89 ± 0.63 | 31.86 ± 0.67 | 31.86 ± 0.65 |
| | GA | 35.76 ± 0.68 | 35.77 ± 0.67 | 35.72 ± 0.61 | 35.79 ± 0.69 | 35.78 ± 0.69 |
| | GC | 36.67 ± 0.61 | 36.68 ± 0.62 | 36.62 ± 0.63 | 36.69 ± 0.62 | 36.69 ± 0.61 |
| | GG | 33.91 ± 0.6 | 33.91 ± 0.6 | 33.89 ± 0.55 | 33.9 ± 0.61 | 33.92 ± 0.6 |
| | GT | 34.57 ± 0.81 | 34.53 ± 0.78 | 34.56 ± 0.76 | 34.54 ± 0.79 | 34.55 ± 0.79 |
| | TA | 34.66 ± 0.46 | 34.64 ± 0.45 | 34.57 ± 0.42 | 34.65 ± 0.46 | 34.66 ± 0.46 |
| | TC | 35.82 ± 0.68 | 35.79 ± 0.67 | 35.76 ± 0.61 | 35.75 ± 0.68 | 35.79 ± 0.69 |
| | TG | 34.62 ± 0.57 | 34.61 ± 0.56 | 34.51 ± 0.54 | 34.62 ± 0.56 | 34.64 ± 0.55 |
| | TT | 35.87 ± 0.9 | 35.87 ± 0.9 | 35.66 ± 0.82 | 35.88 ± 0.9 | 35.86 ± 0.89 |
| TP53 | AA | 35.87 ± 0.92 | 35.84 ± 0.91 | 35.54 ± 0.85 | 35.85 ± 0.87 | 35.86 ± 0.91 |
| | AC | 34.5 ± 0.75 | 34.51 ± 0.81 | 34.4 ± 0.69 | 34.51 ± 0.79 | 34.51 ± 0.79 |
| | AG | 31.85 ± 0.67 | 31.8 ± 0.64 | 31.66 ± 0.57 | 31.83 ± 0.65 | 31.81 ± 0.64 |
| | AT | 32.72 ± 0.83 | 32.68 ± 0.8 | 32.53 ± 0.79 | 32.7 ± 0.81 | 32.73 ± 0.81 |
| | CA | 34.58 ± 0.54 | 34.59 ± 0.54 | 34.46 ± 0.51 | 34.56 ± 0.54 | 34.59 ± 0.56 |
| | CC | 33.89 ± 0.6 | 33.88 ± 0.59 | 33.75 ± 0.49 | 33.88 ± 0.59 | 33.87 ± 0.59 |
| | CG | 32.88 ± 0.57 | 32.89 ± 0.55 | 32.78 ± 0.5 | 32.9 ± 0.54 | 32.92 ± 0.59 |
| | CT | 31.81 ± 0.63 | 31.79 ± 0.63 | 31.68 ± 0.58 | 31.84 ± 0.64 | 31.81 ± 0.63 |
| | GA | 35.76 ± 0.64 | 35.74 ± 0.66 | 35.68 ± 0.59 | 35.77 ± 0.66 | 35.78 ± 0.69 |
| | GC | 36.65 ± 0.63 | 36.65 ± 0.62 | 36.51 ± 0.58 | 36.69 ± 0.63 | 36.64 ± 0.63 |
| | GG | 33.88 ± 0.61 | 33.89 ± 0.58 | 33.74 ± 0.47 | 33.89 ± 0.59 | 33.85 ± 0.57 |
| | GT | 34.51 ± 0.78 | 34.5 ± 0.78 | 34.37 ± 0.7 | 34.51 ± 0.81 | 34.5 ± 0.81 |
| | TA | 34.63 ± 0.45 | 34.62 ± 0.46 | 34.57 ± 0.41 | 34.59 ± 0.44 | 34.64 ± 0.44 |
| | TC | 35.76 ± 0.66 | 35.74 ± 0.66 | 35.7 ± 0.59 | 35.78 ± 0.66 | 35.82 ± 0.68 |
| | TG | 34.59 ± 0.56 | 34.6 ± 0.56 | 34.46 ± 0.51 | 34.6 ± 0.55 | 34.61 ± 0.55 |
| | TT | 35.84 ± 0.92 | 35.85 ± 0.95 | 35.59 ± 0.9 | 35.85 ± 0.93 | 35.88 ± 0.91 |

### CTCF

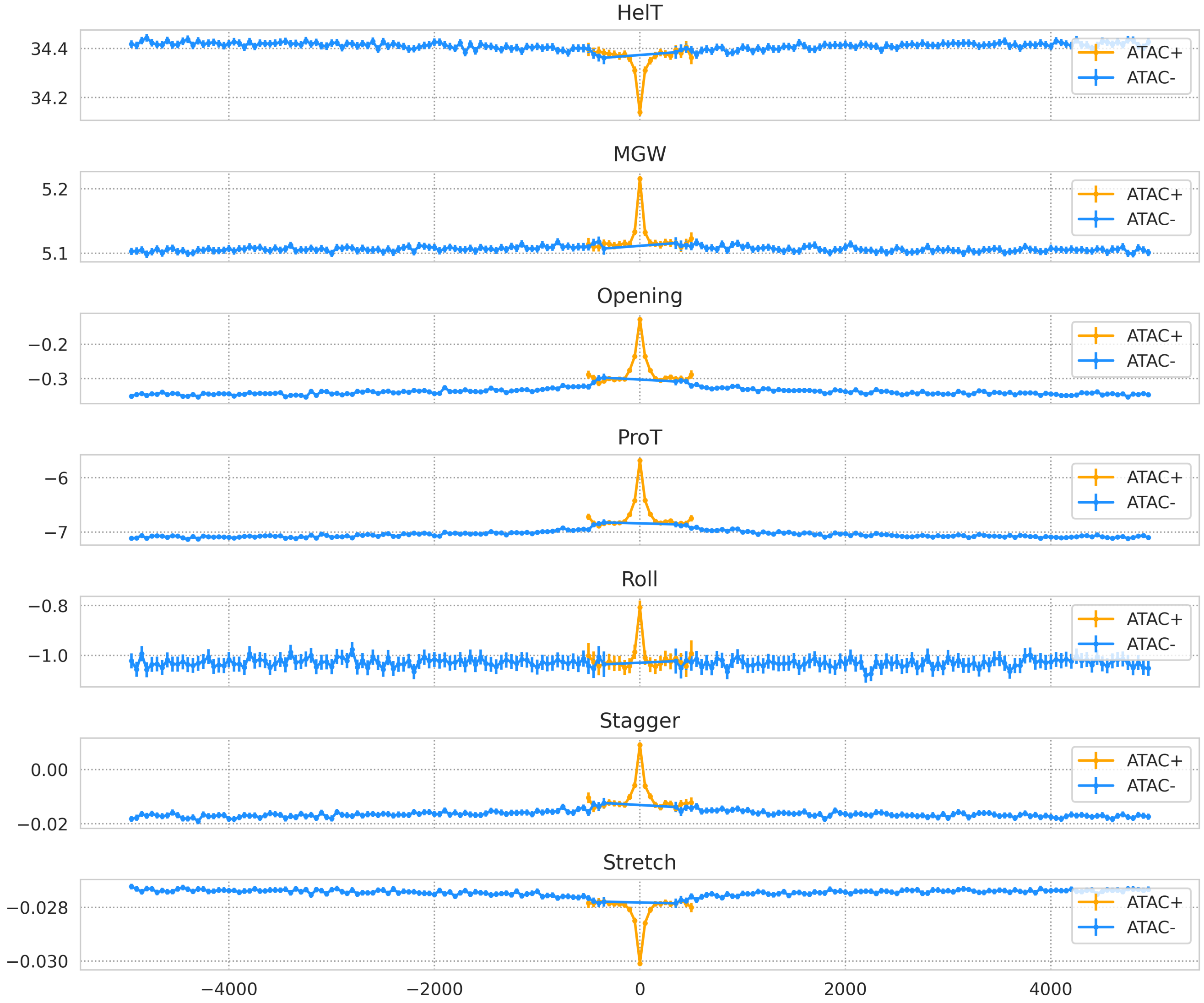


**Fig. S24.** Average DNA shape values (from top to bottom: helical twist, minor groove width, opening, propeller twist, roll, stagger, stretch) per every 25th offset for prolonged peaks in experiments targeting CTCF. We distinguish ATAC+ (orange, k-mers overlapping with an ATAC-seq peak in the same cell line) and ATAC- (blue, k-mers overlapping with **no** ATAC-seq peak in the same cell line) k-mers. Value is shown if at least 25% peaks at position are seen open/closed.

### FOXK2

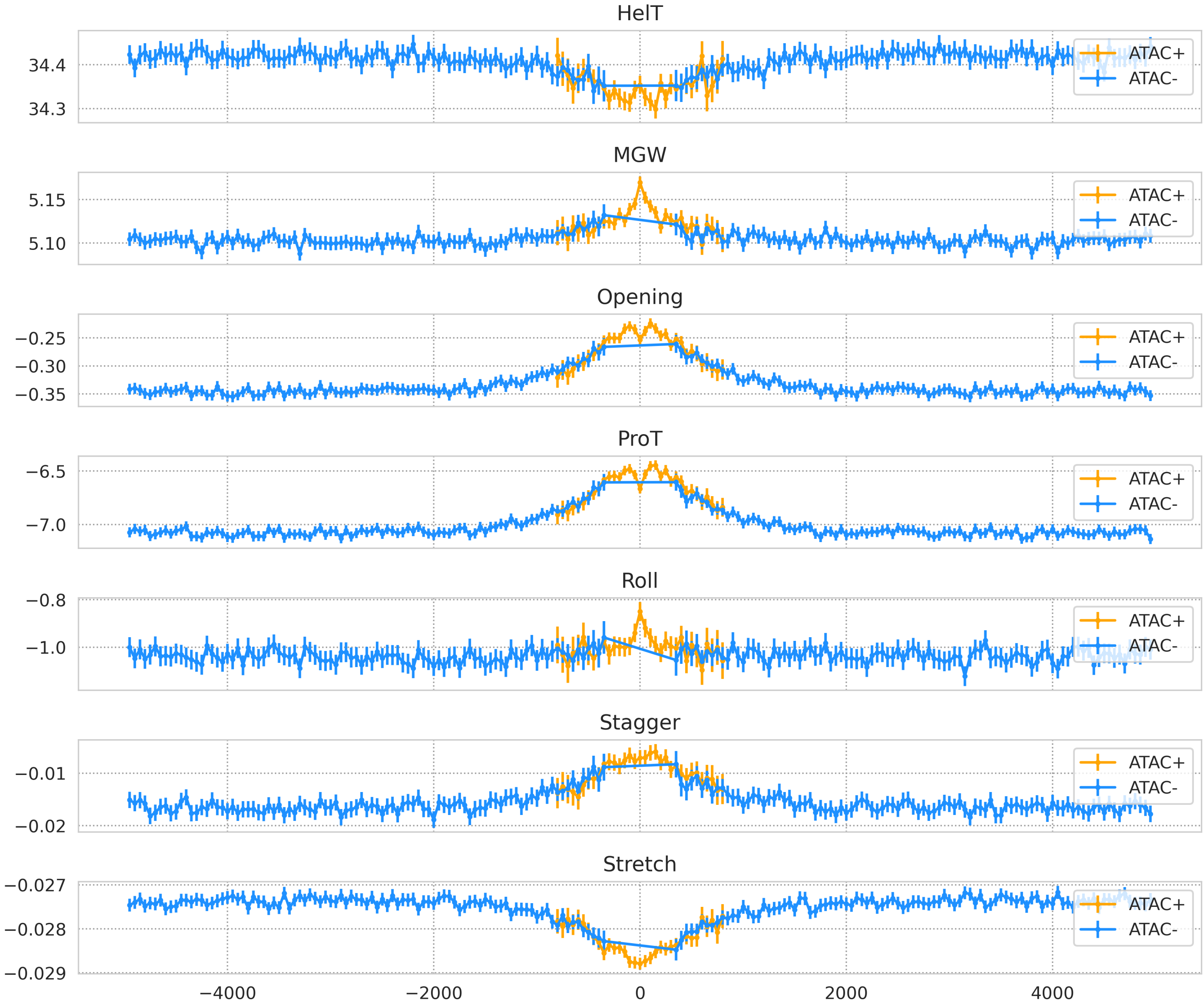


**Fig. S25.** Average DNA shape values (from top to bottom: helical twist, minor groove width, opening, propeller twist, roll, stagger, stretch) per every 25th offset for prolonged peaks in experiments targeting FOXK2. We distinguish ATAC+ (orange, k-mers overlapping with an ATAC-seq peak in the same cell line) and ATAC- (blue, k-mers overlapping with **no** ATAC-seq peak in the same cell line) k-mers. Value is shown if at least 25% peaks at position are seen open/closed.

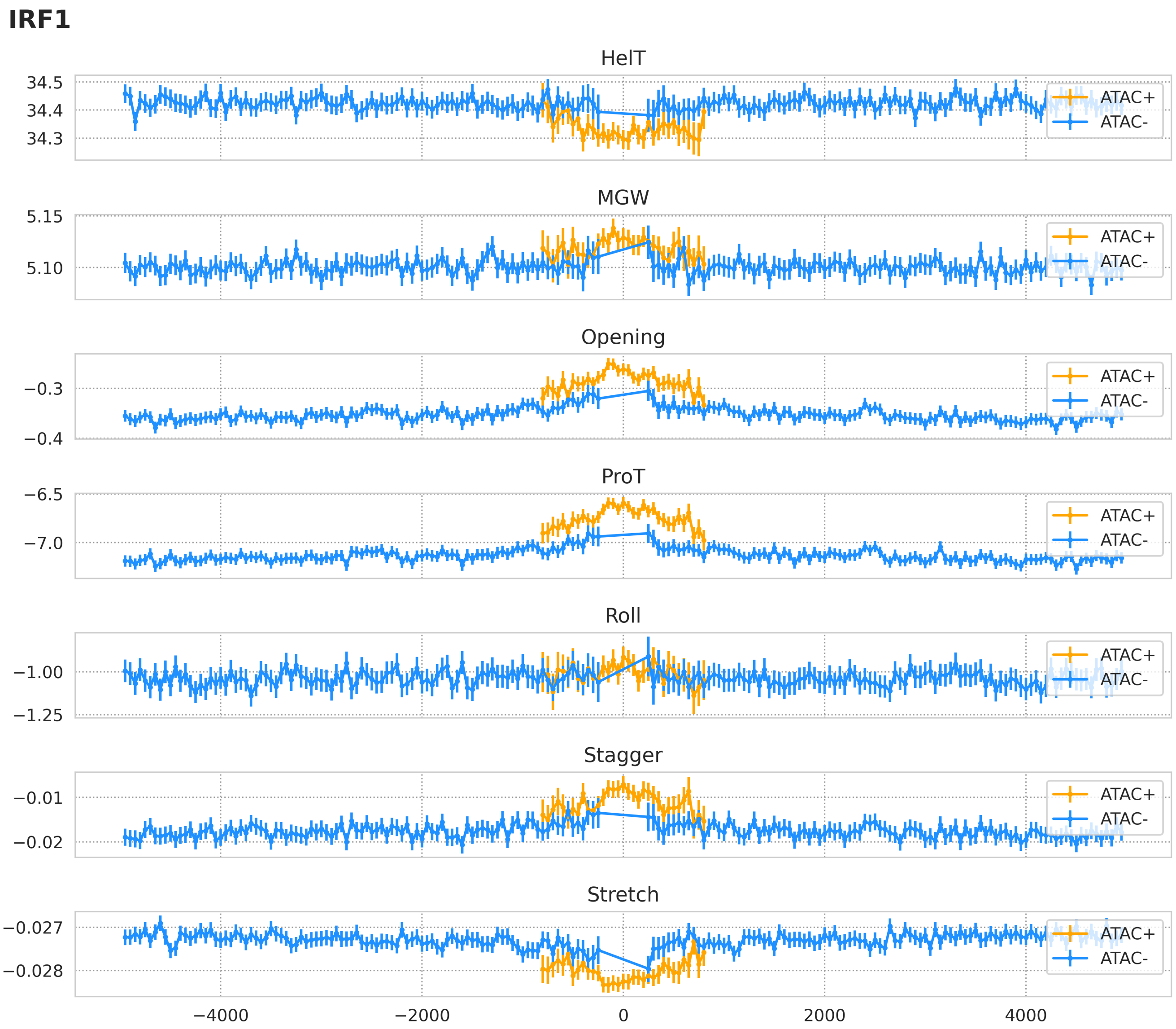


**Fig. S26.** Average DNA shape values (from top to bottom: helical twist, minor groove width, opening, propeller twist, roll, stagger, stretch) per every 25th offset for prolonged peaks in experiments targeting IRF1. We distinguish ATAC+ (orange, k-mers overlapping with an ATAC-seq peak in the same cell line) and ATAC- (blue, k-mers overlapping with **no** ATAC-seq peak in the same cell line) k-mers. Value is shown if at least 25% peaks at position are seen open/closed.

### MEF2A

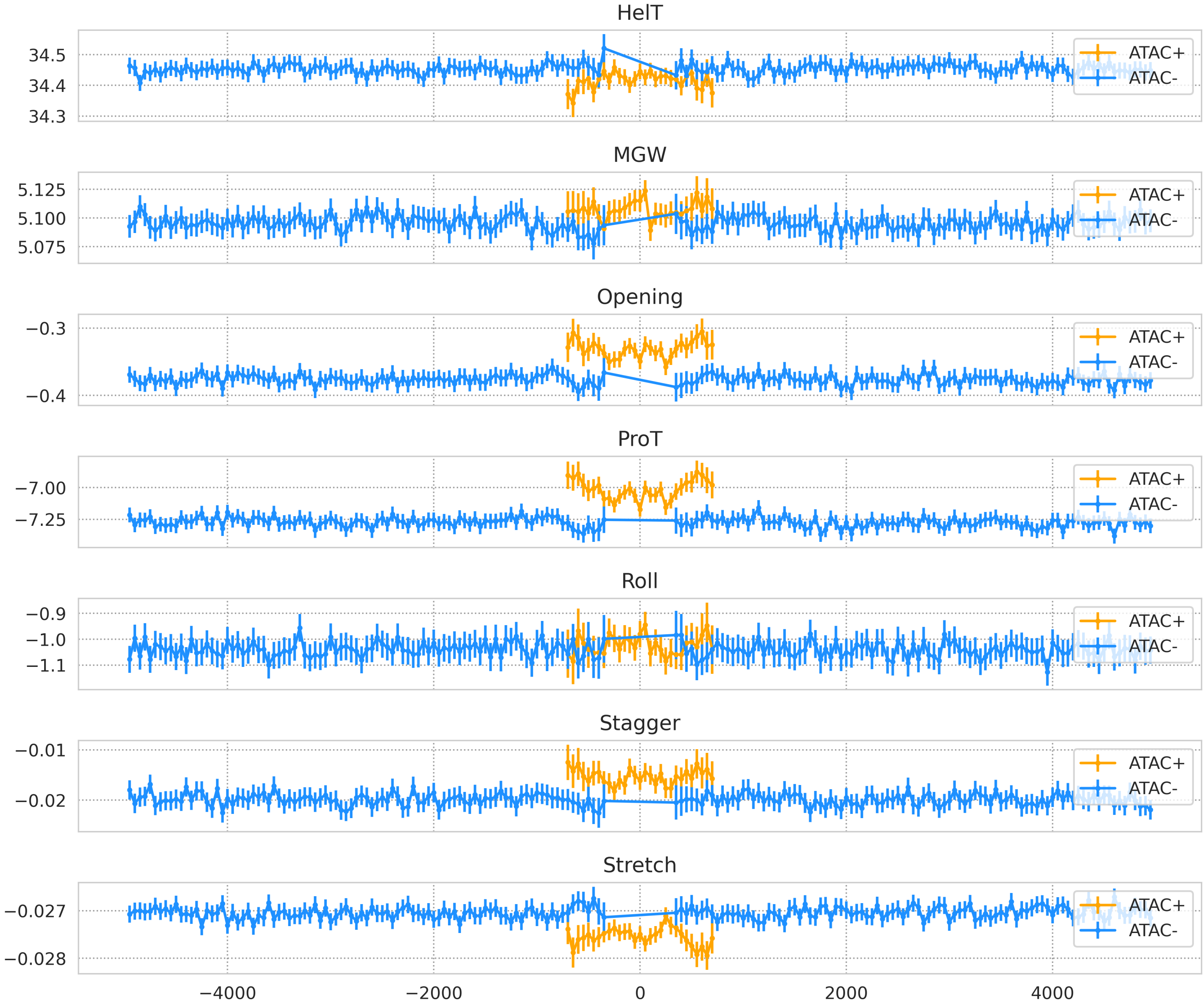


**Fig. S27.** Average DNA shape values (from top to bottom: helical twist, minor groove width, opening, propeller twist, roll, stagger, stretch) per every 25th offset for prolonged peaks in experiments targeting MEF2A. We distinguish ATAC+ (orange, k-mers overlapping with an ATAC-seq peak in the same cell line) and ATAC- (blue, k-mers overlapping with **no** ATAC-seq peak in the same cell line) k-mers. Value is shown if at least 25% peaks at position are seen open/closed.

### MYC

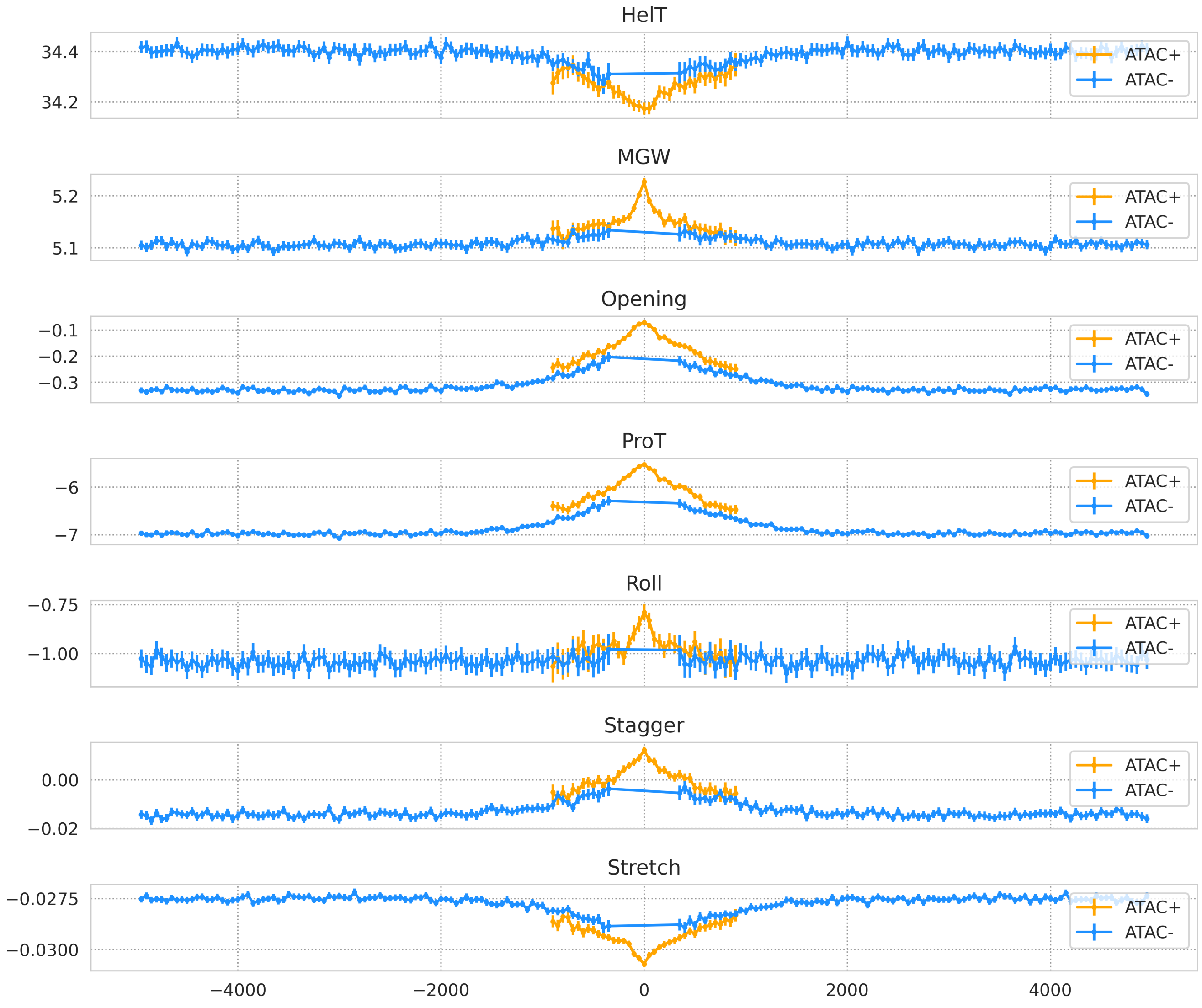


**Fig. S28.** Average DNA shape values (from top to bottom: helical twist, minor groove width, opening, propeller twist, roll, stagger, stretch) per every 25th offset for prolonged peaks in experiments targeting MYC. We distinguish ATAC+ (orange, k-mers overlapping with an ATAC-seq peak in the same cell line) and ATAC- (blue, k-mers overlapping with **no** ATAC-seq peak in the same cell line) k-mers. Value is shown if at least 25% peaks at position are seen open/closed.

### NANOG

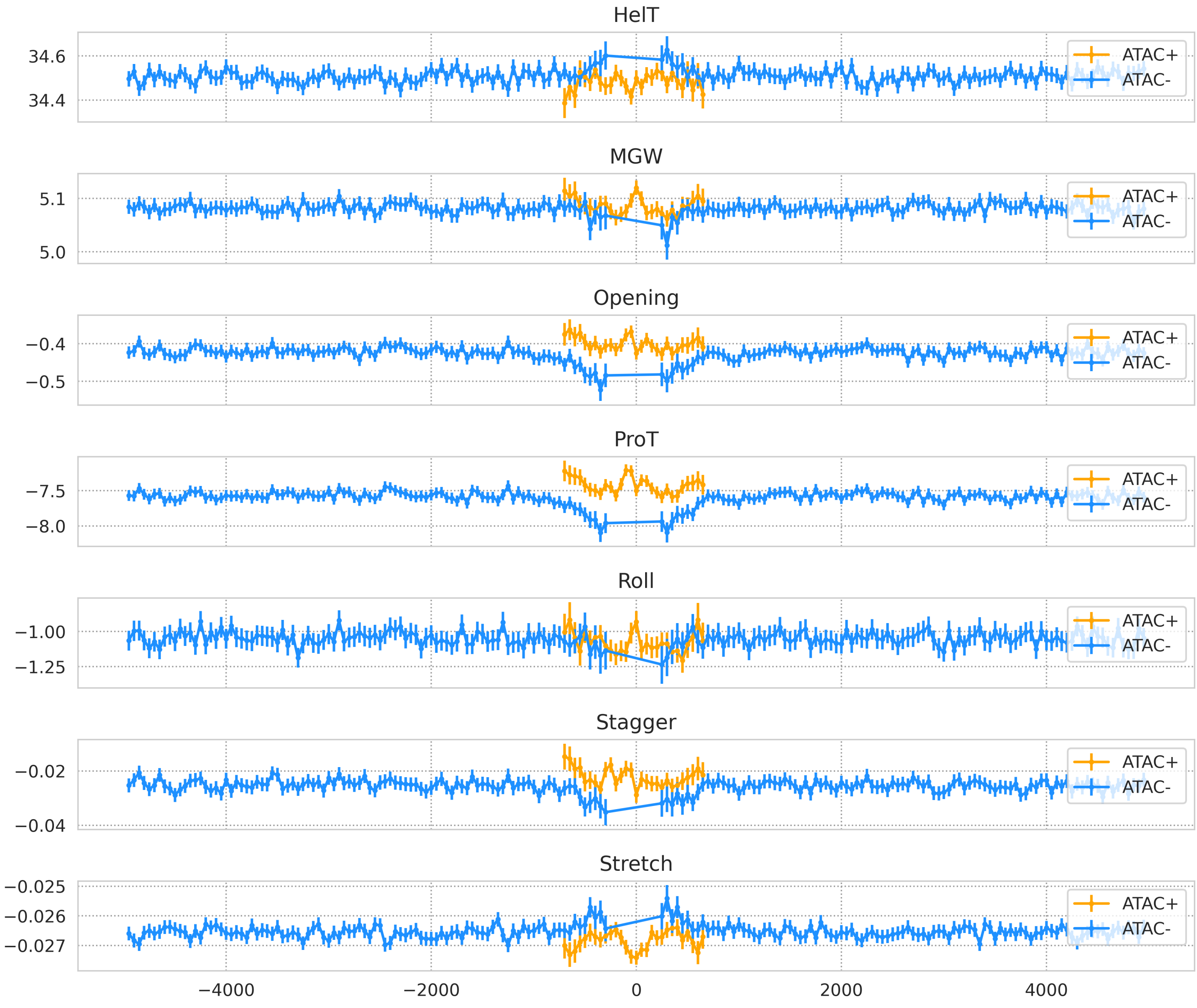


**Fig. S29.** Average DNA shape values (from top to bottom: helical twist, minor groove width, opening, propeller twist, roll, stagger, stretch) per every 25th offset for prolonged peaks in experiments targeting NANOG. We distinguish ATAC+ (orange, k-mers overlapping with an ATAC-seq peak in the same cell line) and ATAC- (blue, k-mers overlapping with **no** ATAC-seq peak in the same cell line) k-mers. Value is shown if at least 25% peaks at position are seen open/closed.

### NFRKB

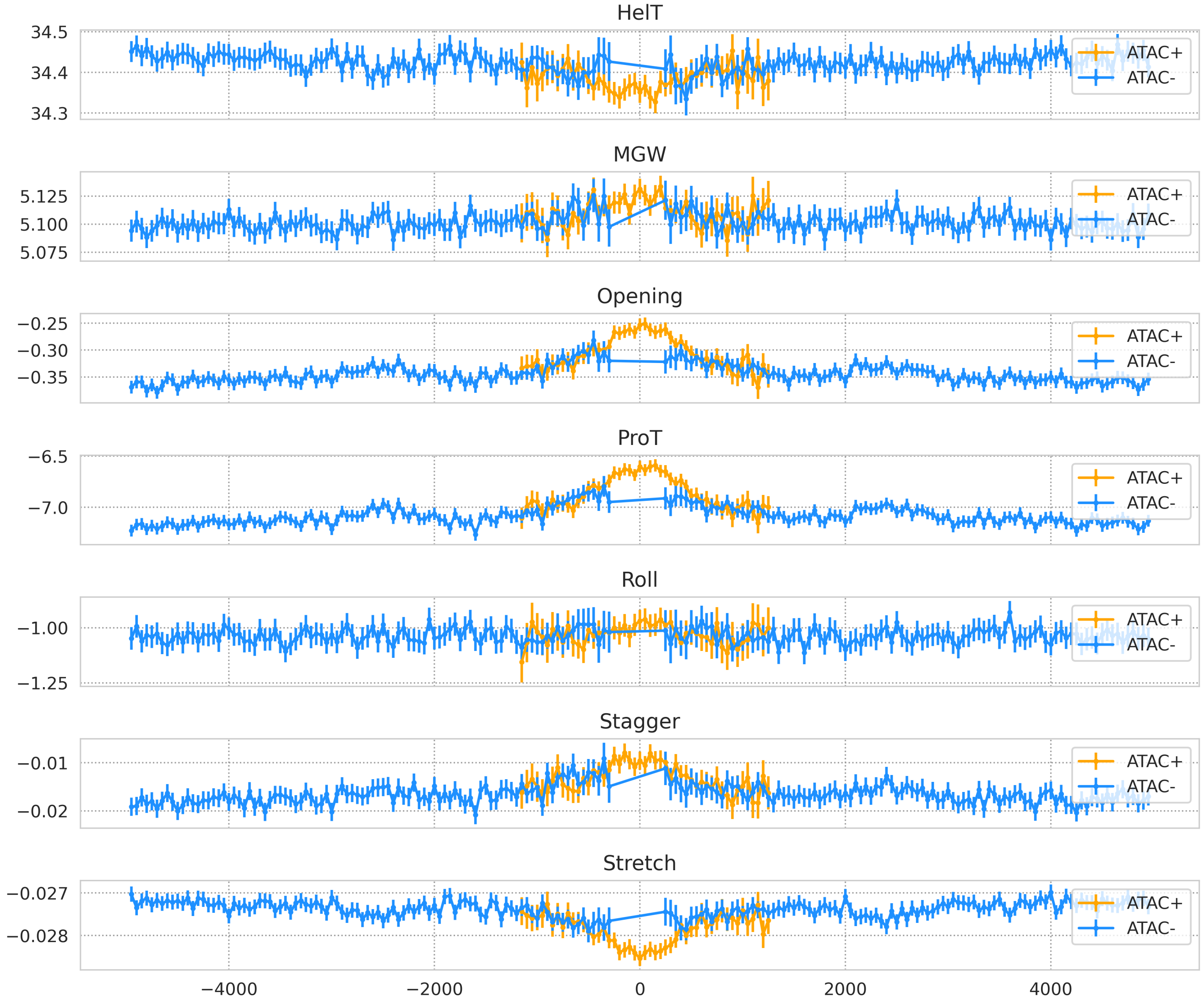


**Fig. S30.** Average DNA shape values (from top to bottom: helical twist, minor groove width, opening, propeller twist, roll, stagger, stretch) per every 25th offset for prolonged peaks in experiments targeting NFRKB. We distinguish ATAC+ (orange, k-mers overlapping with an ATAC-seq peak in the same cell line) and ATAC- (blue, k-mers overlapping with **no** ATAC-seq peak in the same cell line) k-mers. Value is shown if at least 25% peaks at position are seen open/closed.

### RUNX1

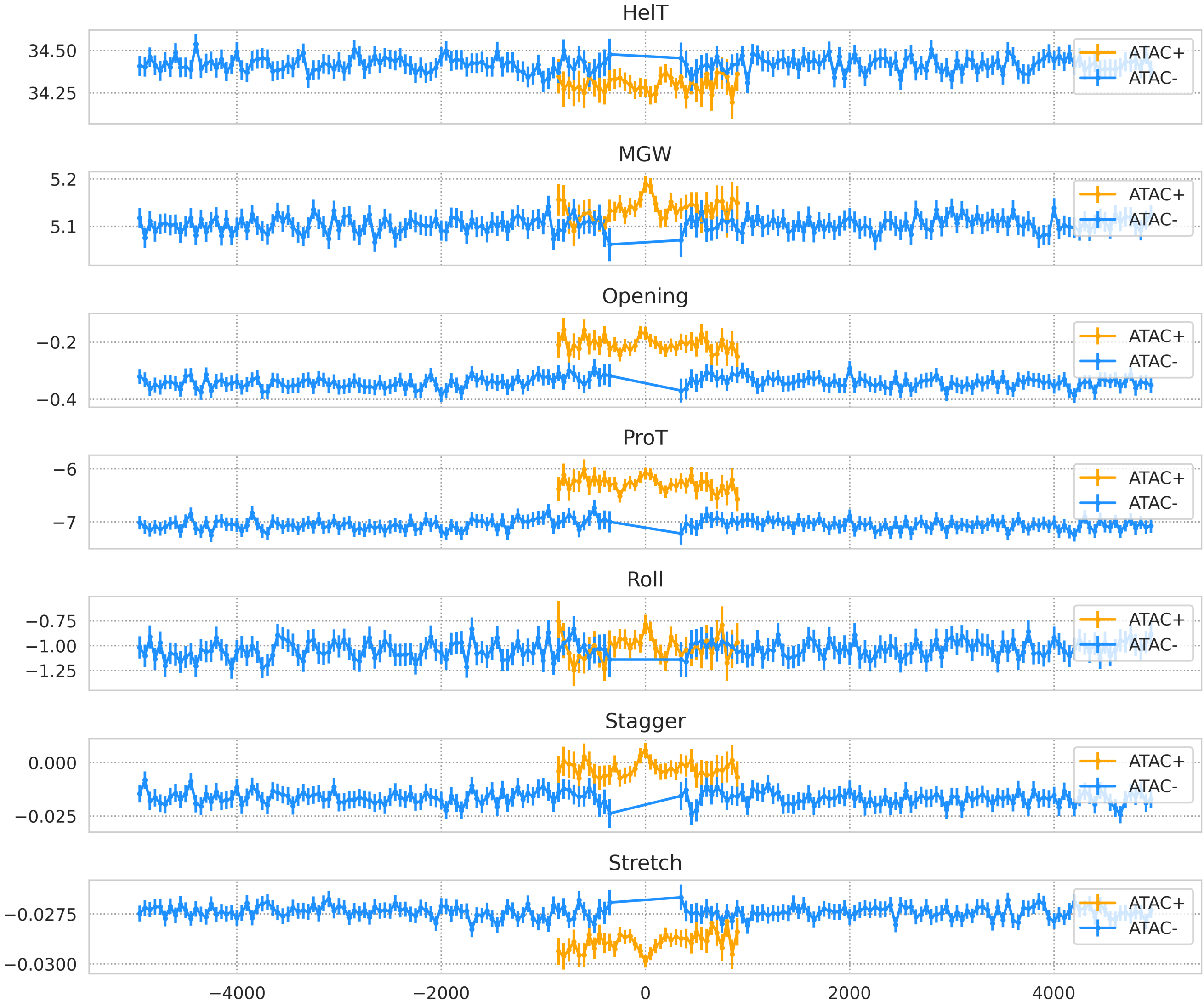


**Fig. S31.** Average DNA shape values (from top to bottom: helical twist, minor groove width, opening, propeller twist, roll, stagger, stretch) per every 25th offset for prolonged peaks in experiments targeting RUNX1. We distinguish ATAC+ (orange, k-mers overlapping with an ATAC-seq peak in the same cell line) and ATAC- (blue, k-mers overlapping with **no** ATAC-seq peak in the same cell line) k-mers. Value is shown if at least 25% peaks at position are seen open/closed.

### SPI1

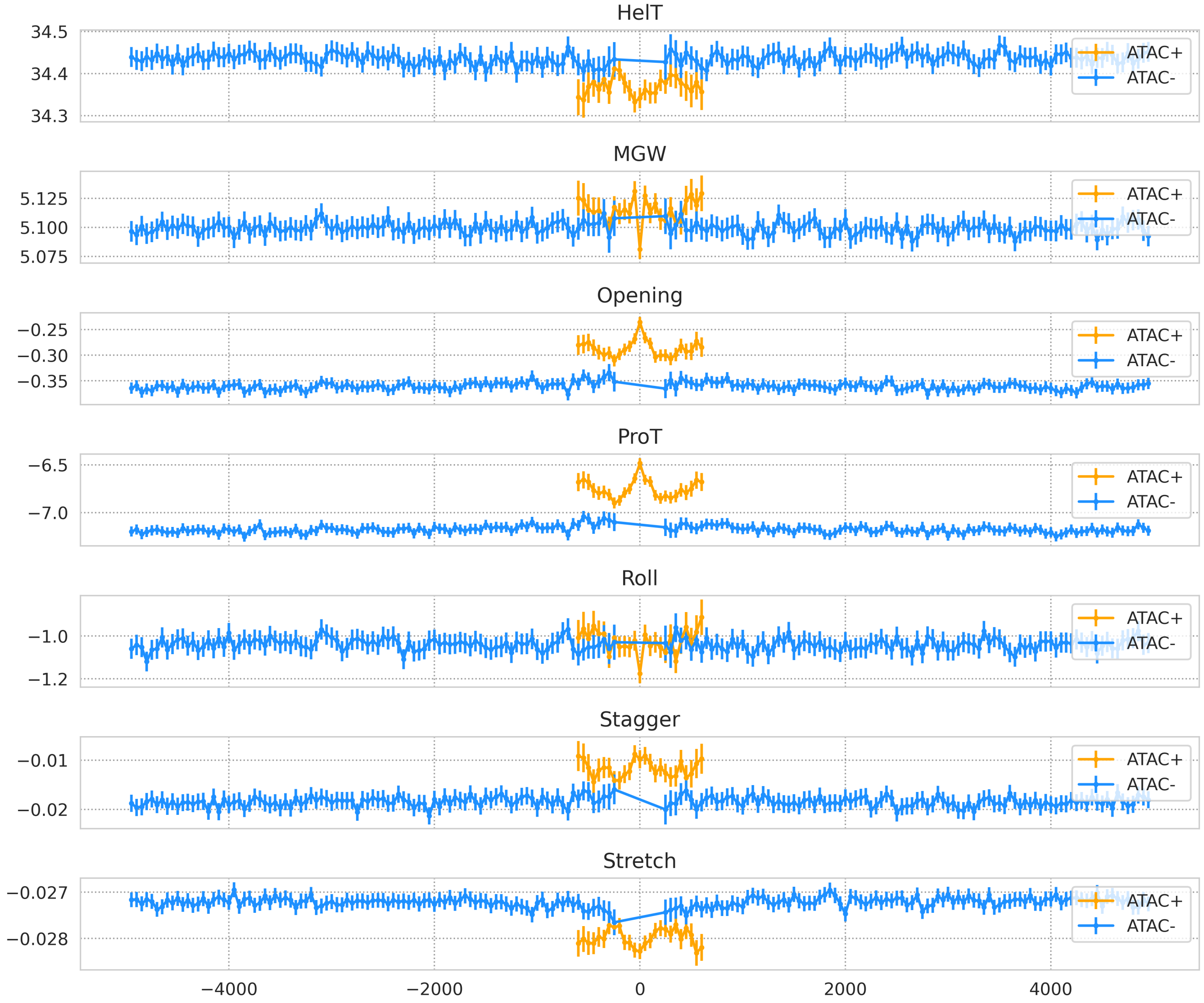


**Fig. S32.** Average DNA shape values (from top to bottom: helical twist, minor groove width, opening, propeller twist, roll, stagger, stretch) per every 25th offset for prolonged peaks in experiments targeting SPI1. We distinguish ATAC+ (orange, k-mers overlapping with an ATAC-seq peak in the same cell line) and ATAC- (blue, k-mers overlapping with **no** ATAC-seq peak in the same cell line) k-mers. Value is shown if at least 25% peaks at position are seen open/closed.

## TP53

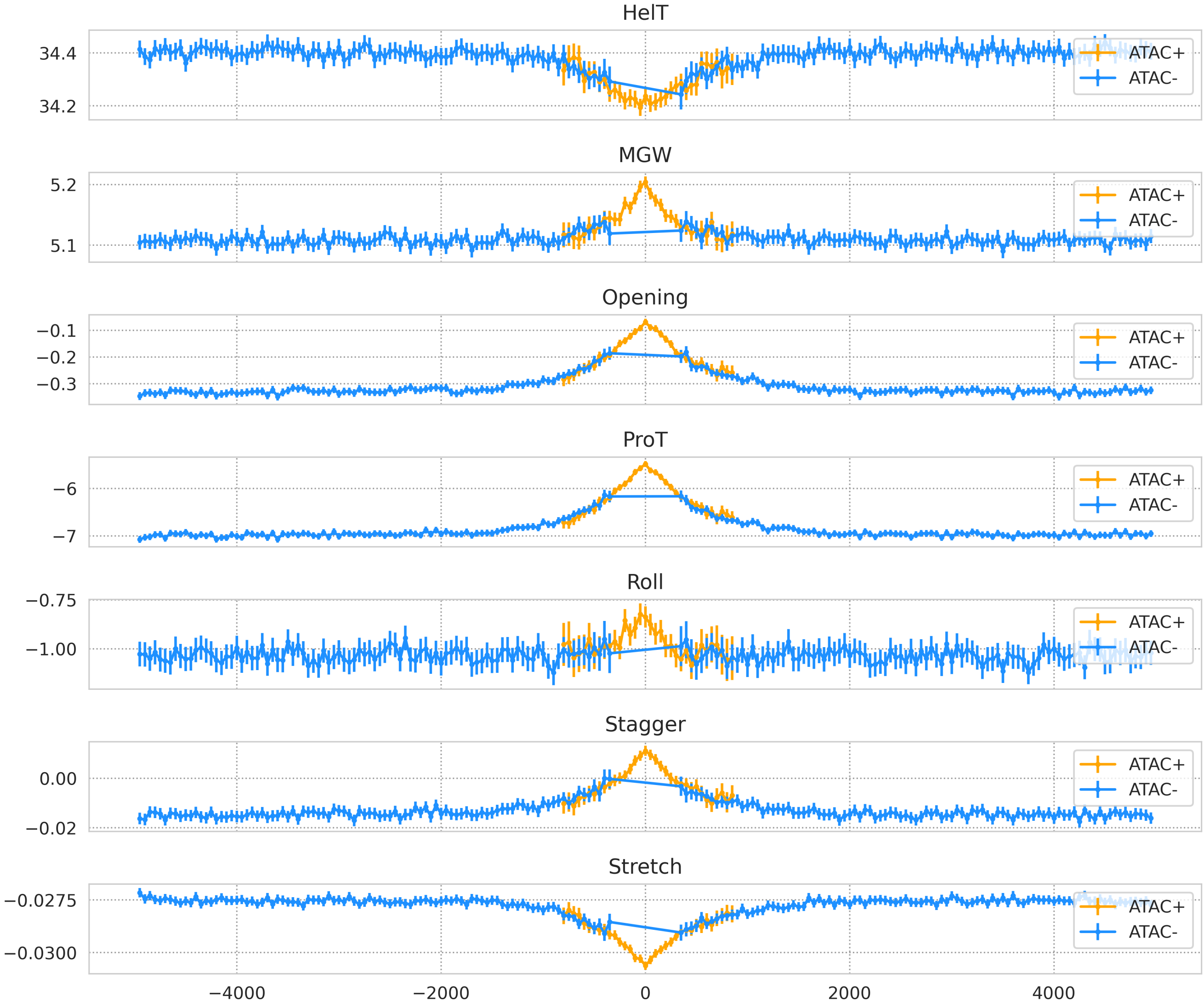


**Fig. S33.** Average DNA shape values (from top to bottom: helical twist, minor groove width, opening, propeller twist, roll, stagger, stretch) per every 25th offset for prolonged peaks in experiments targeting TP53. We distinguish ATAC+ (orange, k-mers overlapping with an ATAC-seq peak in the same cell line) and ATAC- (blue, k-mers overlapping with **no** ATAC-seq peak in the same cell line) k-mers. Value is shown if at least 25% peaks at position are seen open/closed.

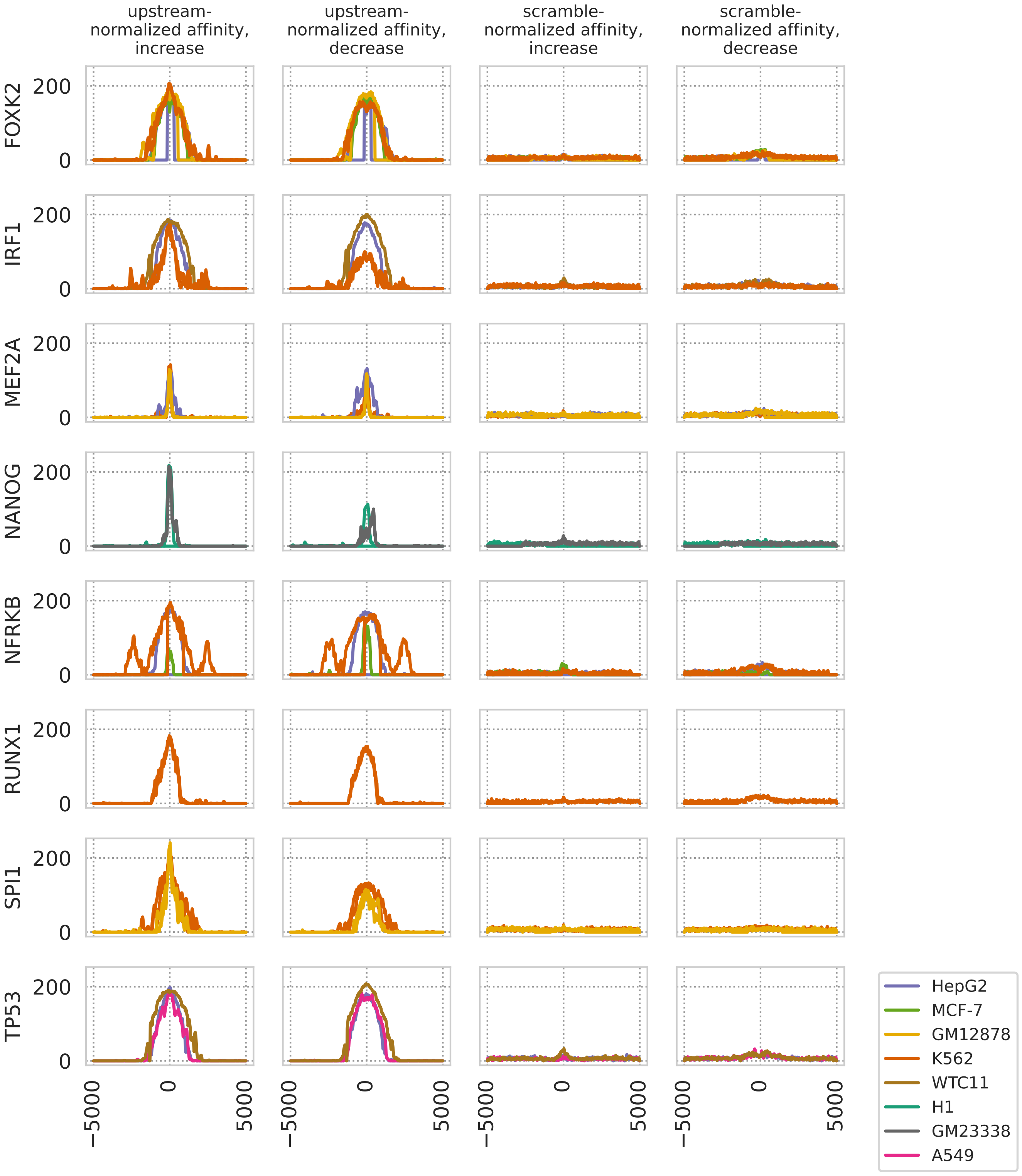


**Fig. S34.** First column: counts of TFs from the HOCOMOCO with increased upstream-normalized affinity when compared to prolonged peak edges, for each tested TF. Second column: counts of TFs from the HOCOMOCO with decreased upstream-normalized affinity when compared to prolonged peak edges, for each tested TF. Third column: counts of TFs from the HOCOMOCO with increased scramble-normalized affinity when compared to prolonged peak edges, for each tested TF. Fourth column: counts of TFs from the HOCOMOCO with decreased scramble-normalized affinity when compared to prolonged peak edges, for each tested TF. Each line color represents a different cell line. For a TF from the HOCOMOCO database, we assess the normalized affinity increases and decreases using the cluster analysis (Methods). The upstream-normalized affinity score is calculated as the k-mer affinity z-score with regards to affinity to sequences 5000 bases upstream of annotated transcription starts of RefSeq genes (hg19). The scramble-normalized affinity score is calculated using a bootstrapped set of k-mers found at a given offset from the putative binding site. Each of these k-mers is then randomly shuffled.

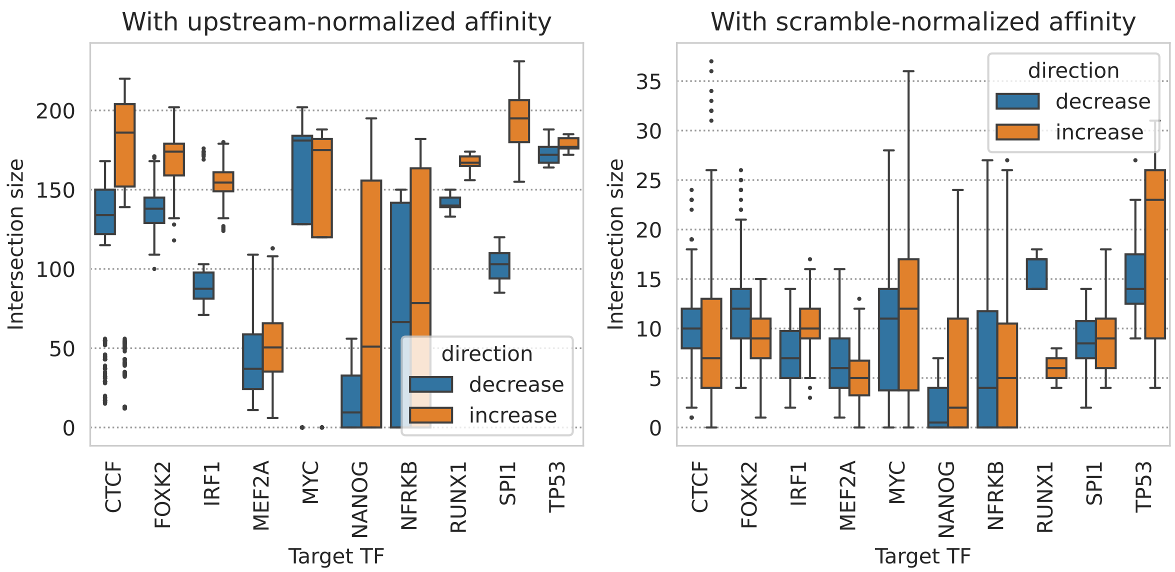


**Fig. S35.** Boxplots showing the distribution of intersection sizes for TFs from the HOCOMOCO database with altered normalized affinity between pairs of experiments targeting the same TF. Each data point represents the maximum number of overlapping TFs exhibiting differential affinity between two such experiments, for offsets up to 100 bp from the putative binding size. The left plot corresponds to the upstream-normalized affinity values, the right one to the scramble-normalized affinity values.

**Table S13.** Portion of experiments that displayed increased/decreased scramble-normalized affinity for a given HOCOMOCO TF at a given offset from the putative binding site. Only HOCOMOCO TFs that were observed inat least 75% of experiments (for the target TF) for at least one offset value are shown. We limit ourselves to the experiment that target TFs in three of more cell lines.

| | | | offset from binding site | | | | | | | | |
|---|---|---|---|---|---|---|---|---|---|---|---|
| Target TF | Change direction | HOCOMOCO TF | -100 | -75 | -50 | -25 | 0 | 25 | 50 | 75 | 100 |
| CTCF | decrease | CPEB1 | 10/11 | 9/11 | 10/11 | 10/11 | 10/11 | 10/11 | 10/11 | 10/11 | 8/11 |
| | decrease | ZN562 | 9/11 | 9/11 | 9/11 | 5/11 | 2/11 | 7/11 | 9/11 | 6/11 | 7/11 |
| | decrease | ZN649 | 9/11 | 9/11 | 6/11 | 4/11 | 4/11 | 3/11 | 3/11 | 7/11 | 8/11 |
| | decrease | ZNF90 | 8/11 | 9/11 | 7/11 | 6/11 | 5/11 | 3/11 | 6/11 | 7/11 | 7/11 |
| | increase | OZF | - | 2/11 | 3/11 | 6/11 | 9/11 | 9/11 | 3/11 | 1/11 | 1/11 |
| | increase | ZBT7A | 5/11 | 4/11 | 9/11 | 8/11 | 7/11 | 8/11 | 8/11 | 6/11 | 3/11 |
| | increase | ZFP57 | - | - | 1/11 | 8/11 | 9/11 | 7/11 | 1/11 | - | - |
| | increase | ZN611 | 1/11 | 3/11 | 6/11 | 8/11 | 10/11 | 9/11 | 5/11 | 3/11 | 2/11 |
| FOXK2 | decrease | CPEB1 | 7/8 | 7/8 | 7/8 | 5/8 | 3/8 | 4/8 | 5/8 | 7/8 | 7/8 |
| | decrease | OZF | 6/8 | 3/8 | 2/8 | 4/8 | 4/8 | 2/8 | 2/8 | 2/8 | 3/8 |
| | decrease | ZN550 | 6/8 | 4/8 | 3/8 | 3/8 | 4/8 | 3/8 | 5/8 | 4/8 | 2/8 |
| | decrease | ZNF90 | 5/8 | 7/8 | 6/8 | 4/8 | 5/8 | 5/8 | 5/8 | 6/8 | 4/8 |
| | increase | MYCN | 5/8 | 3/8 | 5/8 | 5/8 | 6/8 | 5/8 | 6/8 | 5/8 | 5/8 |
| | increase | ZBT7A | 3/8 | 4/8 | 4/8 | 5/8 | 6/8 | 6/8 | 6/8 | 5/8 | 4/8 |
| IRF1 | decrease | HXD10 | 1/4 | - | 3/4 | 3/4 | 2/4 | - | 2/4 | 2/4 | 3/4 |
| | decrease | SIX6 | 1/4 | 1/4 | 2/4 | 2/4 | 3/4 | 3/4 | 2/4 | 1/4 | - |
| | decrease | ZN443 | 1/4 | - | - | 1/4 | 1/4 | 2/4 | 3/4 | 2/4 | 1/4 |
| | decrease | ZN649 | 1/4 | 1/4 | 2/4 | 1/4 | 1/4 | - | 2/4 | 3/4 | 2/4 |
| | decrease | ZNF90 | - | 1/4 | 2/4 | 1/4 | 2/4 | 2/4 | 1/4 | 2/4 | 3/4 |
| | increase | MYCN | 2/4 | 2/4 | 1/4 | 2/4 | 3/4 | 3/4 | 3/4 | 2/4 | 2/4 |
| | increase | OZF | 2/4 | 2/4 | 2/4 | 3/4 | 3/4 | 3/4 | 2/4 | 3/4 | 2/4 |
| | increase | ZN302 | - | 2/4 | 2/4 | 2/4 | 2/4 | 2/4 | 2/4 | 2/4 | 3/4 |
| | increase | ZN430 | 2/4 | 2/4 | 3/4 | 3/4 | 3/4 | 3/4 | 1/4 | - | 1/4 |
| | increase | ZN443 | 3/4 | 2/4 | 2/4 | 2/4 | 1/4 | 1/4 | 1/4 | 1/4 | 2/4 |
| | increase | ZN611 | 2/4 | 3/4 | 4/4 | 4/4 | 2/4 | 3/4 | 2/4 | 3/4 | 3/4 |
| | increase | ZN623 | - | 3/4 | 3/4 | 1/4 | 2/4 | 3/4 | 2/4 | 2/4 | 1/4 |

| | | | | | | | | | | | |
|---|---|---|---|---|---|---|---|---|---|---|---|
| | increase | ZN649 | 2/4 | 2/4 | 2/4 | 2/4 | 3/4 | 2/4 | - | 1/4 | 1/4 |
| MEF2A | decrease | HMGA1 | 1/5 | 2/5 | 2/5 | 1/5 | 1/5 | 1/5 | 2/5 | 2/5 | 4/5 |
| | decrease | OZF | 4/5 | 4/5 | 3/5 | 2/5 | 2/5 | 1/5 | 2/5 | 2/5 | 4/5 |
| | decrease | ZFP57 | 2/5 | - | - | - | 2/5 | 2/5 | 1/5 | 2/5 | 4/5 |
| | decrease | ZN248 | 5/5 | 3/5 | 3/5 | 2/5 | 2/5 | 2/5 | 3/5 | 4/5 | 4/5 |
| | decrease | ZN562 | 3/5 | 4/5 | 3/5 | 4/5 | 4/5 | 3/5 | 4/5 | 3/5 | 3/5 |
| | decrease | ZN649 | 4/5 | 3/5 | 3/5 | 3/5 | 2/5 | 2/5 | 3/5 | 4/5 | 5/5 |
| | increase | MYCN | 3/5 | 3/5 | 2/5 | 2/5 | 3/5 | 5/5 | 5/5 | 2/5 | - |
| | increase | ZN443 | 4/5 | 2/5 | 1/5 | 1/5 | - | - | 1/5 | 2/5 | 2/5 |
| | increase | ZN550 | 3/5 | 1/5 | 1/5 | 3/5 | 4/5 | 2/5 | 3/5 | 3/5 | 1/5 |
| | increase | ZN611 | 1/5 | 1/5 | 3/5 | 3/5 | 5/5 | 4/5 | 1/5 | 1/5 | 1/5 |
| | increase | ZN623 | - | - | 3/5 | 4/5 | 4/5 | 3/5 | 1/5 | - | 1/5 |
| MYC | decrease | CPEB1 | 7/8 | 5/8 | 5/8 | 6/8 | 6/8 | 7/8 | 6/8 | 5/8 | 6/8 |
| | decrease | IRX2 | 4/8 | 4/8 | 5/8 | 4/8 | 6/8 | 6/8 | 5/8 | 4/8 | 4/8 |
| | decrease | ZN580 | 6/8 | 6/8 | 4/8 | 5/8 | 6/8 | 5/8 | 6/8 | 6/8 | 4/8 |
| | decrease | ZN736 | 5/8 | 3/8 | 1/8 | 3/8 | 5/8 | 7/8 | 5/8 | 6/8 | 5/8 |
| | decrease | ZNF2 | 2/8 | 2/8 | 4/8 | 5/8 | 6/8 | 4/8 | 3/8 | 4/8 | 4/8 |
| | decrease | ZNF90 | 6/8 | 4/8 | 2/8 | 3/8 | 3/8 | 3/8 | 4/8 | 4/8 | 5/8 |
| | increase | MYCN | 6/8 | 6/8 | 6/8 | 5/8 | 4/8 | 4/8 | 6/8 | 6/8 | 6/8 |
| | increase | OZF | 3/8 | 3/8 | 4/8 | 3/8 | 6/8 | 5/8 | 5/8 | 3/8 | 2/8 |
| | increase | ZN302 | 4/8 | 2/8 | 2/8 | 4/8 | 5/8 | 5/8 | 5/8 | 6/8 | 3/8 |
| | increase | ZN454 | 3/8 | 7/8 | 7/8 | 6/8 | 7/8 | 7/8 | 7/8 | 5/8 | 2/8 |
| | increase | ZN578 | 2/8 | 3/8 | 5/8 | 6/8 | 7/8 | 5/8 | 4/8 | 4/8 | 4/8 |
| | increase | ZN611 | 4/8 | 5/8 | 7/8 | 7/8 | 6/8 | 6/8 | 6/8 | 4/8 | 3/8 |
| TP53 | decrease | CPEB1 | 3/3 | 3/3 | 3/3 | 3/3 | 3/3 | 3/3 | 3/3 | 3/3 | 3/3 |
| | decrease | HMGA1 | 1/3 | 1/3 | 3/3 | 2/3 | 1/3 | 1/3 | 1/3 | 2/3 | 2/3 |
| | decrease | ZN248 | 3/3 | 2/3 | 2/3 | 2/3 | 2/3 | 2/3 | 1/3 | 3/3 | 3/3 |
| | decrease | ZN550 | 1/3 | 1/3 | 2/3 | 2/3 | 1/3 | 2/3 | 3/3 | 3/3 | 3/3 |
| | decrease | ZN562 | 2/3 | 2/3 | 3/3 | 3/3 | 2/3 | 1/3 | 2/3 | 3/3 | 2/3 |
| | decrease | ZN580 | 2/3 | 2/3 | 3/3 | 1/3 | 2/3 | 2/3 | 2/3 | 3/3 | 2/3 |
| | decrease | ZNF90 | 2/3 | 3/3 | 2/3 | 2/3 | 1/3 | 2/3 | 2/3 | 2/3 | 2/3 |
| | increase | MYCN | 3/3 | 2/3 | 2/3 | 2/3 | 3/3 | 3/3 | 3/3 | 2/3 | 3/3 |
| | increase | ZN302 | 2/3 | 2/3 | 2/3 | 2/3 | 3/3 | 2/3 | 2/3 | 3/3 | 2/3 |
| | increase | ZN623 | 1/3 | - | - | 1/3 | 2/3 | 3/3 | 3/3 | 3/3 | 1/3 |
